# Reticulate evolution and climate driven diversification shaped the origin and geographic structure of *Linum bienne*

**DOI:** 10.64898/2026.06.17.732954

**Authors:** Beatrice Landoni, Juan Viruel, Yann Bourgeois, Robin G. Allaby, Adrian C. Brennan, Rocio Pérez-Barrales

## Abstract

▯ Hybridization, incomplete lineage sorting and climatic oscillations can interact across evolutionary timescales, but their combined effects on plant diversification remain difficult to resolve. We investigated how these processes shaped the origin and geographic structure of *Linum bienne*, the putative progenitor of cultivated flax.
▯ We integrated genus-level Angiosperms353 phylogenomics across *Linum* with plastome analyses, low-depth nuclear resequencing, demographic inference and environmental niche modelling across the range of *L. bienne*. This framework allowed us to assess phylogenetic discordance, test for reticulation and reconstruct lineage history through time.
▯ Phylogenetic discordance was widespread across our genus-level sampling and largely consistent with incomplete lineage sorting. However, branch-length and maximum-likelihood network analyses detected additional signal of gene flow at the node including *L. bienne*, cultivated flax and the closest relatives. Within *L. bienne,* plastid and nuclear data recovered four geographically structured lineages across the species range, with cytonuclear discordance and demographic analyses supporting repeated secondary contact.
▯ Our results show that the evolutionary history of L*. bienne* reflects the interaction between deep reticulation, incomplete lineage sorting and Pleistocene range dynamics. This multiscale perspective highlights *L. bienne* as a genetically complex species and illustrates how reticulate evolution and climatic change can jointly shape plant diversification.

## Introduction

Whole genome duplications (WGD), hybridization, and introgression are recurrent features of plant evolution often associated with climatic and geological change, contributing to lineage persistence, diversification, and adaptation (Cai et al., 2019; Leroy et al., 2019; Wagner et al., 2019; Kadereit & Abbot, 2021; Lovell et al., 2021; Stull et al., 2023; Chen et al., 2025). Yet, understanding how WGD, gene flow, lineage divergence, and incomplete lineage sorting (ILS) interact across taxonomic and temporal scales remains challenging (Stull et al., 2023). Integrating phylogenomics, phylogeography, and population genomics provides a powerful framework to connect macro- and microevolutionary processes, disentangling multiple sources of phylogenetic conflict (Edwards et al., 2022).

The Mediterranean Basin offers a natural context for understanding how these evolutionary processes interact during plant diversification (Myers et al., 2000; Medail & Diadema, 2009; Perret et al., 2023). Its rugged topography, combined with isolation and reconnection cycles driven by the Messinian Salinity Crisis and Pleistocene glaciations, promoted geographic isolation, range shifts, and complex phylogeographic structure across plant lineages at genus and species level (Hewitt, 1999; Comes & Abbott, 2001; Magri et al., 2006, 2008; Santos-Gally et al., 2012; Nieto Feliner, 2014, 2023; Scheunert & Heubl, 2014; Alonso-Blanco et al., 2016; Díaz-Pérez et al., 2018; Migliore et al., 2012, 2018; Sancho et al., 2018; Toledo et al., 2020; Viruel et al., 2020b; García-Verdugo et al., 2021; Kadereit & Abbot, 2021). These dynamics created opportunities for secondary contact and reticulate evolution, raising the question of how climatic oscillations shaped both divergence and subsequent gene flow in Mediterranean plants (Scheunert & Heubl, 2014; Díaz-Pérez et al., 2018; Sancho et al., 2018; Nieto Feliner, 2014; Kadereit & Abbot, 2021). Furthermore, hybridization and polyploidization in several Mediterranean lineages are associated with environmental heterogeneity, niche differentiation, and historical climate dynamics (Lopez-Jurado et al., 2019; Valdés-Florido et al., 2024a and 2024b).

This evolutionary complexity is reflected in the region’s exceptional diversity of crop wild relatives (Vincent et al., 2019). Many Mediterranean crop wild relatives show strong phylogeographic structure and histories shaped by climatic fluctuations, dispersal, and gene flow with domesticated relatives (Diez et al., 2014; Mousavi-Derazmahalleh et al., 2018; Gutaker et al., 2019; Smykal et al., 2017; Zunino et al., 2024). Such processes can influence the distribution of genetic variation and provide variation relevant for adaptation, crop improvement, and range expansion (Cortes et al 2013; Hufford et al., 2013; Burgarella et al., 2019; Viruel et al., 2021; Grabowski et al., 2024; Chen et al., 2026). Thus, Mediterranean crop wild relatives provide a valuable framework for linking historical range dynamics, reticulate evolution, and the evolutionary potential of domesticated lineages.

Within Mediterranean lineages including crops, *Linum* stands out for studying how phylogeographic structuring, chromosome change, and reproductive diversification interact during plant evolution. The genus displays marked diversity in floral morphology, reproductive strategies, life-history traits, and chromosome number, and includes cultivated flax or *Linum usitatissimum* (Armbruster et al., 2006; Ruiz-Martín et al., 2018; Maguilla et al., 2021; Pérez-Barrales & Armbruster, 2023; Gutiérrez-Valencia et al., 2022). Chromosome evolution has partly driven the diversification of the genus, with ploidy variation associated with life-history traits and colonization patterns (Afonso et al. 2021, 2023; Valdés-Florido et al. 2023; Vanrell et al. 2024; but see also Ockendon, 1968). Transcriptomic analyses also support an ancient genome duplication in the lineage containing cultivated flax (Sveinsson et al., 2014). Conversely, evidence for hybridization or reticulation in *Linum* remains limited at the genus level, relying largely on karyotypic evidence from a few lineages (Bolsheva et al., 2017, 2022). More broadly, because phylogenetic inference in *Linum* has relied mostly on a limited set of plastid and nuclear ITS markers (McDill et al., 2009; Ruiz-Martín et al., 2018; Maguilla et al., 2021), the extent of phylogenetic conflict and reticulate evolution across *Linum* largely unresolved.

*Linum bienne*, the closest wild relative and putative progenitor of cultivated flax (Fu & Allaby, 2010; Gutaker et al., 2019; Weiss & Zohary, 2011), occupies a key position within the genus. Together with *Linum villarianum* and *L. usitatissimum*, it forms a clade within section *Linum* whose origin and phylogenetic relationships remain poorly resolved (Ruiz-Martín et al., 2018; Maguilla et al., 2021; Villalvazo-Hernandez et al., 2022). Cytogenetic and molecular evidence suggests a reticulate origin for this lineage involving hybridization between ancestors with similar chromosome numbers, possibly related to *L. decumbens* and/or *L. narbonense,* followed by a late Miocene WGD around 5–9 million years ago (Bolsheva et al., 2017; Cai et al., 2019; Wang et al., 2012). Conflicts between nuclear and plastid markers in resolving the relationship between *L. bienne* and *L. villarianum* further suggest that reticulate processes may have contributed to the history of this lineage (Villalvazo-Hernandez et al., 2022).

*Linum bienne* also provides an opportunity to investigate how climatic gradients and historical climate change have shaped population structure and intraspecific diversification. Distributed across the Mediterranean Basin and Atlantic Europe, populations show evidence of genetic and phenotypic differentiation across latitudinal, elevational, and climatic gradients (Uysal et al., 2010, 2012; Soto-Cerda et al., 2014; Fu, 2023; Landoni et al., 2024). Documented secondary contact and gene flow between *L. bienne* and cultivated flax (Gutaker et al., 2019) raise questions about the timing, frequency, and evolutionary consequences of gene flow both before and during the process of domestication. The evolutionary significance of this process is further underscored by evidence that introgression from wild *L. bienne* populations to cultivated flax might have contributed to the northwards expansion of the crop by facilitating adaptation to shorter growing seasons following domestication in the Near-Eastern Fertile Crescent (Gutaker et al., 2019).

This study investigates how ILS, reticulation, and climatic oscillations interact across taxonomic scales to shape lineage origin, diversification, and intraspecific structure. We integrate genus-level phylogenomics in *Linum*, population genomics in *L. bienne*, and past-to-present environmental niche modeling. First, using target-capture data, we reconstruct phylogenetic relationships across *Linum* to evaluate the relative contributions of ILS and gene flow to phylogenetic conflict. Second, we analyze plastid and low-depth nuclear genomes to infer lineage structure, demographics, and gene flow across *L. bienne* distribution, while including *L. usitatissimum*. Finally, we model Pliocene-to-present climatic suitability to assess its impact on lineage persistence, range shifts, and secondary contact in *L. bienne*. Combined, these multi-scale data test how Mediterranean climate history and genomic complexity jointly shaped the evolution of a crop wild relative.

## Material & Methods

### Genus-level phylogenomics, phylogenetic conflict, and divergence dating in *Linum*

#### Taxon sampling, sequencing, and recovery of nuclear and plastid loci

Nineteen *Linum* species were selected to infer a genus-wide phylogeny using Angiosperms353 data (Johnson et al., 2019) to date major nodes and investigate reticulation involving *L. bienne* (Table S1). Sampling represented the two main *Linum* clades, its taxonomic sections, and the closest relatives of *L. bienne* (Ruiz-Martín et al., 2018). We also included three geographically distinct *L. bienne* individuals and one European *L. usitatissimum* cultivar (Table S1). Sampling combined ten species from the Tree of Life repository (Baker et al., 2022) with newly generated sequences for seven additional species. Following a comprehensive Malpighiales phylogeny by Xi et al. (2012), *Phyllanthus* spp. and *Euphorbia mesembryanthemifolia* were included as outgroups. For new samples, DNA was extracted using a modified CTAB protocol (Viruel et al., 2019), and genomic libraries were prepared, enriched using the Angiosperms353 kit following Razanamaro et al. (2025), and sequenced on an Illumina HiSeq X platform. Reads for all samples were quality-filtered using FastQC v0.11.9 (Andrews, 2010) and fastp v1 (Chen et al., 2018) with settings: --trim_poly_g and --trim_poly_x, and discarding of unpaired reads. The HybPiper pipeline v2.3.2 (Johnson et al., 2016) was used to recover plastid (Pokorny et al., 2024) and single-copy Angiosperms353 nuclear genes (Johnson et al., 2019; MacLay et al., 2021). Low-quality or problematic samples and genes flagged with paralog warnings were removed following strict quality control thresholds based on hybpiper statistics (Table S2). For more information about filtering, see Supplementary Information.

#### Phylogenetic inference, discordance and reticulation analyses, and divergence dating

The resulting 208 nuclear and 36 plastid gene DNA sequences were aligned separately with MAFFT v7.310 (Katoh & Standley, 2013) with settings: --nuc --localpair --maxiterate 1000. Alignments were trimmed in trimAl v1.2 with “automated1” option (Capella-Gutiérrez et al., 2009). Two supermatrices partitioned by gene were generated for plastid and nuclear datasets respectively, by concatenating gene alignments (https://github.com/santiagosnchez/ConcatFasta). Maximum likelihood (ML) phylogenetic inference was done using IQ-TREE v1.6.12 (Nguyen et al., 2015) with a GTR+G model on: (1) individual nuclear gene alignments; (2) with the nuclear supermatrix; (3) with the plastid supermatrix, each with 1000 bootstrap replicates. The GTR+G model was chosen because it provides a flexible and robust model for different types of datasets, including Angiosperms353 and plastid datasets (Abadi et al., 2019; Pokorny et al., 2024; Zuntini et al., 2024), while maintaining comparability with previous phylogenetic studies in *Linum* (Ruiz-Martin et al., 2018). Unrooted best ML gene trees were summarized with ASTRAL-III v5.7.1 (Zhang et al., 2018). All best ML trees and the coalescent Astral tree were then rooted in Phyx v1.3 (Brown et al., 2017) with *E. mesembryanthemifolia* as outgroup. Rooted plastid and nuclear best ML (supermatrix) or coalescent (Astral) trees were compared to assess if cytonuclear discordance in *Linum*. Topologies retrieved previously (Bolsheva et al., 2017; Bolsheva et al., 2022; McDill et al., 2009; Patel et al., 2026; Ruiz-Martin et al., 2018; Svenisson et al., 2014; Villalvazo et al., 2022) were compared with present ones.

ASTRAL nuclear gene-tree discordance was quantified with PhyParts and visualised with phypartspiecharts.py (https://github.com/mossmatters), complemented by a DensiTree of supermatrix bootstrap replicates. Phytop v0.3.2 (Hong-Yun et al., 2025) and QuIBL (Edelman et al., 2019) were used to assess ILS and gene flow at genus level based on the Astral tree, while *L. bienne* hybrid origin was tested via ML in PhyloNet v3.8.2 (Than et al., 2008) focusing on the *L. bienne* node (including *L. narbonense*, *L. decumbens* and *L. grandiflorum*). Additionally, we also tested if individual plastid genes could vary in their support of topologies found in the present and previous studies at the *L. bienne* node to assess whether sampling of specific plastid genes in previous works might have also resulted in apparent cytonuclear discordance. Detailed description of the material and methods regarding Phytop, QuIBL, PhyloNet, and plastid gene analyses are reported in Supplementary Information.

Finally, the supermatrix ML and Astral nuclear trees, and the plastid ML tree were time-calibrated in treePL v1.0 (Smith & O’Meara, 2012) using a *Linum* crown fossil constraint of 33.9–37.2 Ma (Cavagnetto & Anadón, 1996; Ruiz-Martín et al., 2018) and empirically chosen calibration settings (available at https://github.com/beaLando/linomics). All figures and statistical analyses in the present and following sections were performed using R v4.3.0 (R Core Team, 2021). All phylogenies were plotted using the R packages ape v5.6-1 (Paradis & Schliep, 2019), phytools (Revell, 2012), and ggplot2 v3.3.3 (Wickham, 2016), and post-processed in Inkscape v1.1.1 (Inkscape Project, 2020).

### Range-wide population genomics, plastome variation, and demographic history of *L. bienne*

#### Population sampling, library preparation, and genome sequencing

Sampling encompassed the native range of *L. bienne*, covering 56 populations (88 individuals), five *L. usitatissimum* cultivars, and one *L. narbonense* individual (Table S3). DNA extraction and shotgun libraries followed Viruel et al. (2019). Equimolar pooled (150 × 150 bp) libraries were sequenced at Novogene (Beijing, China) in an Illumina HiSeq X Lane (Illumina, San Diego, California, USA) achieving ∼ 3 million reads/sample (1-2X). Reads were quality-filtered and trimmed in FastQC v0.11.9 (Andrews, 2010) and fastp v1 (Chen et al., 2018) with settings: --trim_poly_g and --trim_poly_x, –dedup, quality ≥20, length ≥40, and discarding of unpaired reads. Clean reads were then mapped separately to *L. usitatissimum* plastid and nuclear genomes to conduct phylogeography and population genomic analyses. Three individuals (L65, L30, L18) representing the main *L. bienne* lineages identified here (see Results) were re-sequenced at high-depth (∼ 30 million reads; 30X) to improve variant calling and demographic inference based on nuclear data (see Supplementary Information). DNA extraction and sequencing of these three individuals was outsourced at Eurofins Genomics (Germany) and conducted with Illumina HiSeq technology (NovaSeq 6000 S4 PE150 XP).

#### Nuclear population structure, spatial genetic diversity, and demographic inference

Trimmed short reads were mapped to the *L. usitatissimum* reference genome (ASM1066527v2; Zhang et al., 2020) with BWA MEM v0.7.17 (Li et al., 2009). Genotype likelihoods for low-coverage data were estimated using ANGSD v0.940 (Korneliussen et al., 2014), and variants were filtered with Samtools (options -GL 1 -uniqueOnly 1 -remove_bads 1 -trim 0 -C 50 -baq 1 -minMapQ 20 -minQ 20 -skipTriallelic 1) so that only biallelic loci with a mapping quality of 20 or more were retained. Details about SNP calling and masking of variants based on mappability and artefactual heterozygosity using individuals sequenced at high depth can be found in Supplementary Information.

For population structure and relatedness analyses, SNPs were further filtered based on: p-value >= 1×10-6, missing data/site <= 66%, coverage depth <= 250x. Linkage disequilibrium was addressed by subsampling SNPs every 10kb. Finally, 16,890 polymorphic nuclear sites were retained. A genetic distance tree was obtained in ngsDist (Vieira et al., 2016), with pairwise deletion of missing data and estimation of branch support (100 block-bootstrap replicates, 20 SNPs/block). We used ngsADMIX (Skotte et al., 2013) to calculate each individual’s assignment probability across one to nine genetic clusters, henceforth termed “lineages” for consistency with plastome analyses. Individuals were assigned to a cluster when having a percentage of assignment larger than 60%. A Principal Component Analysis was run in PCANGSD v1.36.1 (Meisner & Albrechtsen, 2018) given genotype likelihoods. Because *L. bienne* is a highly selfing species, individual heterozygosity cannot work as a proxy for local genetic diversity. First, RAB, a relatedness coefficient robust to inbreeding (Hedrick & Lacy, 2015), was estimated in ngsRelate v2 (Hanghøj et al., 2019) to identify and exclude putative clones (RAB > 0.75). Then, we used the matrix of pairwise genetic distances to compute the average genetic distance between all individuals found in a predefined radius of five latitude and longitude degrees.

Given *L. bienne* lineages, demographic inference was performed using likelihood in fastsimcoal2.8 (Excoffier et al., 2013), and using coalescence rate estimation along genomes in MSMC2 and MSMC-IM (Schiffels & Wang, 2020; Wang et al., 2020). In individual pairs with RAB > 0.25 (closely related), the individual with highest depth coverage was retained. *Linum bienne* central lineage (see Results) was not included due to its low sample size. Imputation of missing genotypes was run in STITCH v1.7.3 (Davies et al., 2016) and genome-wide substitution rates were estimated based on LASTZ v1.04.22 (Harris, 2007) estimation of genome divergence after aligning genomes of different *Linum* species closely related to *L. bienne* (see Supplementary Information). This resulted in an estimated mutation rate of 3.3 x 10^−9^ substitutions/site/generation and a 1yr generation time, used for demographic inference with an inbreeding coefficient of 0.95. Ten random haplotype pairs were selected per lineage to reduce computational load. For fastsimcoal, we used frequency spectra inferred by ANGSD (see Supplementary Information). Three demographic models were compared: (1) no gene flow between the three lineages (strict isolation - SI); (2) constant and asymmetric gene flow between populations (constant migration - IM); (3) secondary contact after the split between lineages (secondary contact - SC). For MSMC2 and MSMC-IM, imputed genotypes were used to estimate coalescence rates after additional variant filtering based on coverage depth (see Supplementary Information). Coalescence rates were converted to demographic parameters based on mutation rates and generation times reported above.

#### Plastome phylogeography, cytonuclear discordance, and divergence dating within L. bienne

Filtered reads were mapped to the *L. usitatissimum* plastid genome (NCBI accession: NC_036356.1; de Santana-Lopes, 2018) with Bowtie2 v2.3.5 (Langmead & Salzberg, 2013). SNPs were called with bcftools v1.9 (Li et al., 2009) assuming a haploid genome, then filtered (quality ≥30; depth 100–354). Consensus plastid genomes were aligned with MAFFT and cleaned with trimAL (“automated1”), yielding 1,852 variants. Plastome alignments were analysed with ML in IQ-TREE (GTR+ G), with *L. narbonense* as outgroup based on previous Ruiz-Martín et al., 2018. The best plastome ML tree was compared to the nuclear genetic distance tree (see above) to detect cytonuclear discordance. Additionally, to test if *L. villarianum* was part of any *L. bienne* lineage, an ML tree was reconstructed in IQ-TREE (GTR+ G) using a concatenated matrix of plastid genes (*matK*, *ndhF*, *trnL-trnF*) retrieved from Villalvazo-Hernandez et al. (2022) for *L. villarianum* and by extracting the same regions from our *L. bienne* plastome alignment via Muscle v5 (Edgar, 2004).

Divergence times among *L. bienne* lineages were estimated from the plastome phylogeny using BEAST v1.10.4 (Drummond & Rambaut, 2007) under a GTR+G model, a Yule prior, an uncorrelated lognormal clock, and 1,000,000,000-chain length, to accommodate variable branch evolutionary rates. Calibration used secondary points derived from the treePL *Linum* dated phylogeny. However, dating of plastid and Astral or supermatrix nuclear trees produced different ages for the *L. bienne+L. usitatissimum* crown node (set as monophyletic), so that three BEAST runs were performed with the following priors for such node: 14.5 My (SD = 1); 0.4 My (SD = 1); 2.5 My (SD = 1). An age of 15 My or 18 My (SD = 2) was set as prior in both supermatrix (nuclear, plastid) or Astral (nuclear) calibrated runs respectively for the root node (*L. narbonense+L. bienne+L. usitatissimum*). These priors were assumed to be normally distributed being secondary calibration points. A 10% burn-in was applied in TreeAnnotator, and convergence visually inspected in Tracer. If effective sample sizes (ESS) for BEAST parameters were > 200, they were considered indicative of sufficient posterior sampling. The parameters’ distribution was plotted with the R package rwty (Warren et al., 2017).

### Environmental niche of *L. bienne* from the Pliocene to present

*Linum bienne* occurrence data were retrieved from GBIF.org (https://doi.org/10.15468/dl.nhbuhr accessed on 03/07/2025) and bioclimatic layers were retrieved for the present via WorldClim (Fick & Hijmans, 2017) and for eight past time-periods (Holocene to Pliocene) via pastclim (Barreto et al., 2023; Holden et al., 2019). Details about data preparation for environmental niche models (ENMs) are described in Supplementary Information. After data preparation, Maxent (dismo v1.3-5; Hijmans et al., 2017), GLM, and GAM ENMs were fitted in R using GBIF occurrence and present bioclimatic data for a restricted, well-sampled area (Portugal, Spain, France). Model performance was evaluated using AUC (Area Under the Curve), MCS (Miller Calibration Slope), TSS (True Skill Statistic), and climate dissimilarity was assessed with MESS (modeEva v3.0; Barbosa et al., 2016). After evaluation, GAM was used to predict favourability across the whole species’ distribution for present and past climatic layers. Favourability was then averaged across time-periods, while also calculating its standard deviation (SD) as a proxy for climatic stability in time. Within areas where mean favourability > 0.1 (considered non-negligible for *L. bienne* occurrence), if SD > 0.18 (85^th^ quantile) the area was considered climatically unstable, while SD < 0.18 indicated climatically stable areas, likely important for long-term persistence.

## Results

### Genus-level phylogenomics, phylogenetic conflict, and divergence dating in *Linum*

Hybpiper statistics with sample and gene filtering details are reported in Supplementary Information and Table S1-S2. In brief, 208 out of 353 nuclear Angiosperms353 genes and 24 out of 26 individuals (all 19 *Linum* species; Table S1) were retained based on hybpiper statistics (Table S2). Instead, based on plastid data, 36 out of 72 genes and 21 individuals were retained (17 *Linum* species with exclusion of *L. decumbens* and *L. hologynum* given data quality; Table S1-S2).

The supermatrix and Astral nuclear trees, as well as the plastid tree, showed largely congruent, well-supported deep nodes (bootstrap > 90%), dividing the genus into two major clades (Figure 1A-C). Clade 1 contained *L. tenue*, *L. flavum*, *L. tenuifolium*, *L. strictum*, and *L. macraei*, whereas Clade 2 was represented by two *Linum* sections: sect. *Dasylinum*, with *L. viscosum* and *L. hirsutum,* and sect. *Linum*, itself divided into two main nodes: (1) *L. perenne*, *L. lewisii*, *L. leoni*, and (2) *L. narbonense, L. decumbens, L. grandiflorum, L. hologynum, L. marginale, L. bienne* (hereafter called *L. bienne* node). Within (2), *L. narbonense* was sister to *L. decumbens* + *L. grandiflorum*. The latter were then sister to *L. bienne* + *L. usitatissimum* and *L. marginale* + *L. hologynum*. Despite overall agreement and high bootstrap support at most nodes (> 90%), conflicting phylogenetic signal in nuclear data increased toward younger nodes according to phyparts, accompanied by disagreement between bootstrap trees (Figure S1), HybPiper paralog warnings, frequent for the *L. bienne* node (Table S1), and mismatches between the nuclear supermatrix versus the Astral and plastid tree topologies for *L bienne* samples (Figure 1A-C).

**Figure 1.**
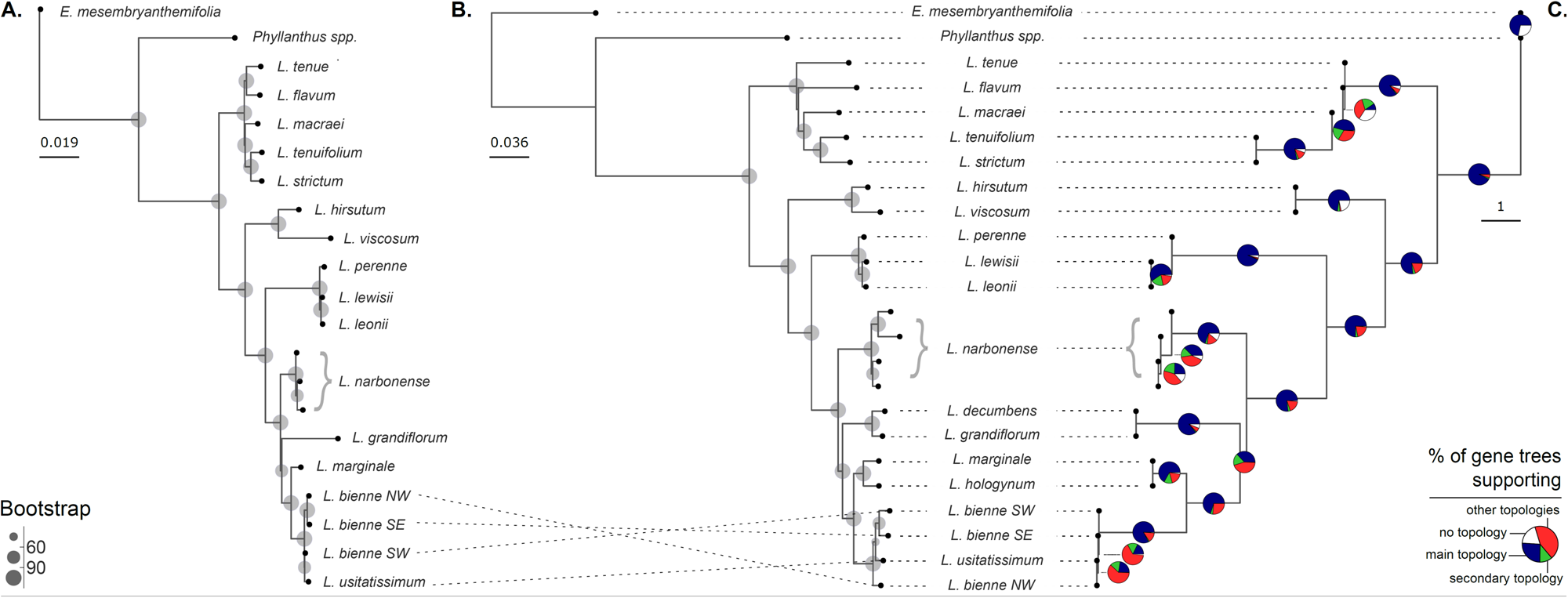
Genus-level phylogenetic relationships and gene-tree discordance in *Linum*. Phylogenetic relationships inferred from (A) 36 plastid genes, and (B) 208 Angiosperms353 nuclear genes, concatenated into a supermatrix using maximum likelihood in IQ-TREE, (C) 208 Angiosperms353 nuclear gene trees summarized under the multispecies coalescent in ASTRAL. Circles in (A) and (B) indicate bootstrap support from 1000 replicates. Pie charts in (C) show the proportion of nuclear gene trees supporting the main topology, the second most frequent topology, alternative topologies, or missing data at each node. The three analyses recovered broadly congruent deeper relationships within *Linum*, but phylogenetic discordance increased towards shallower nodes, particularly around the node including *L. bienne*, cultivated flax (*L. usitatissimum*) and their closest sampled relatives.

Phytop identified ILS as the major driver of nuclear gene tree discordance based on quartet analysis (Figure S2A). ILS was particularly high at the *L. bienne* node (i-ILS > 84%) suggesting that topologies where *L. narbonense* was not sister to all other species at this node were also frequent. This aligns with the branch subtending the split between *L. narbonense* and the other species at the *L. bienne* node being very short across plastid and nuclear phylogenies (Figure 1A-C), possibly indicating rapid diversification of these lineages. QuIBL confirmed this pattern based on branch lengths, with ca. 66.7% discordant triplets (572 out of 858), but only 0.9% of triplets (8 out of 858) supporting an ILS+introgression model (deltaBIC > 10; Table S4). Moderate levels of average introgression were found within *Linum* sections, with higher average introgression estimated for *L. bienne* (Figure S2B).

Interestingly, all statistically significant discordant triplets were placed at the *L. bienne* node, with *L. bienne* often found as outgroup of *L. narbonense* and *L. decumbens/L. grandiflorum*, or as outgroup of *L. decumbens/L. grandiflorum* and *L. marginale/L. hologynum*. The proportion of gene trees supporting ILS+introgression discordant triplets at this node ranged between 9% and 28%, being more frequent for combinations of *L. narbonense* and *L. decumbens*/*L. grandiflorum* (Table S4).

Despite strong ILS, PhyloNet ML network analysis of the *L. bienne* node showed that adding 1-2 hybrid nodes resulted in a more likely topology than the backbone tree, with a plateau in likelihood reached for 3 hybrid nodes (Figure 2A). Additionally, networks constructed explicitly specifying *L. bienne* as a hybrid were always more likely than networks without any hybrid taxa specified. We then examined topologies for the ML analysis where *L. bienne* was specified as a hybrid taxon. Different topologies with similar likelihood were found for 1-2 hybrid nodes (Figure 2B-E). Ignoring hybrid nodes for which the minor hybrid edge had y < 0.1, as suggested by Solís-Lemus & Ané (2016), indicated that *L. bienne* always derived from a hybrid node between either ancestors of *L. bienne + L. decumbens + L. grandiflorum* and *L. decumbens + L. grandiflorum* (Figure 2D-E). A minor hybrid edge with y < 0.1 was often also found between a ghost lineage sister to all taxa of interest and *L. bienne* (Figure 2B-C, 2E). Interestingly, we did not find any apparent evidence of swapped plastid-nuclear topology at this node (Figure 1A-C), often considered a sign of past hybridization. This is possible if hybridization occurred between lineages that had already partly diverged from *L. narbonense* as shown in Figure 2 based on PhyloNet. Instead, we show that individual plastid markers can support a swapped topology relative to the main one retrieved at this node (Table S5). Such swapped topology is common in previous studies using a small number of plastid and ITS markers, with *L. decumbens + L. grandiflorum* as outgroup to *L. narbonense + L. marginale + L. bienne* (Table S6).

**Figure 2.**
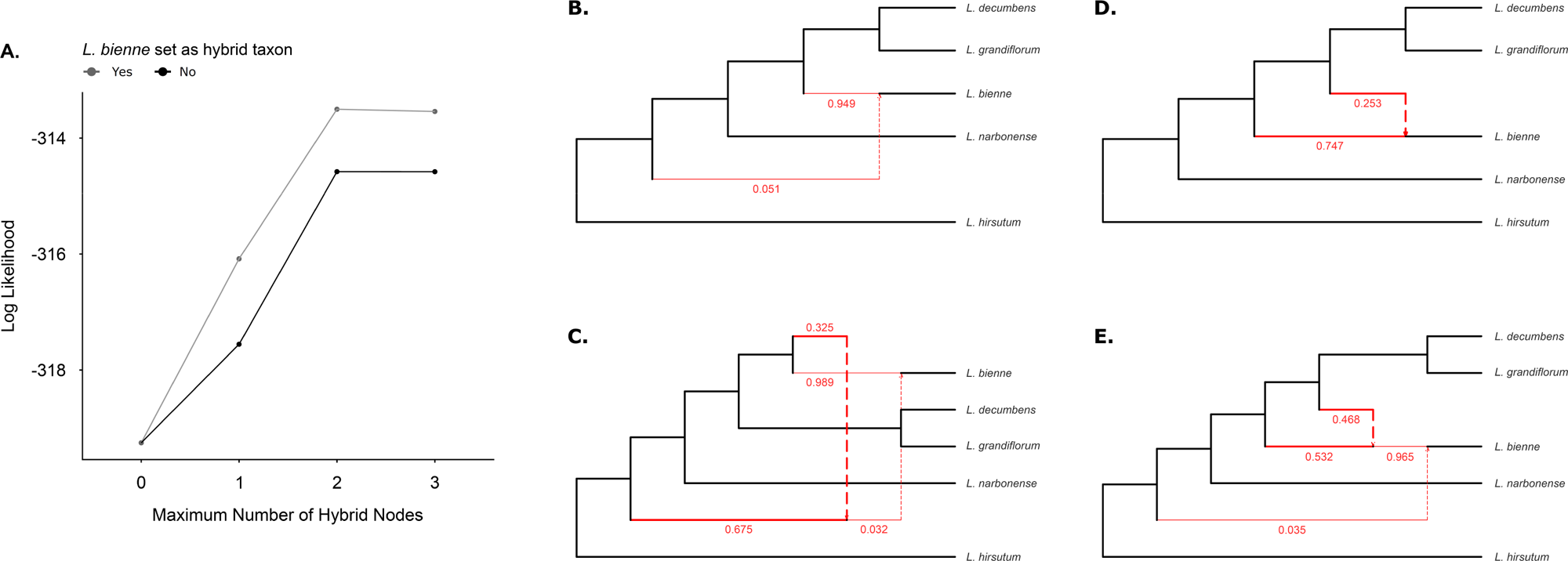
Network analyses support reticulation at the *Linum bienne* node. Maximum-likelihood (ML) phylogenetic network analyses were conducted in PhyloNet for the node including *L. narbonense, L. grandiflorum, L. decumbens* and *L. bienne*, using 190 best ML nuclear gene trees as input (the 208 initial nuclear gene trees were filtered based on presence of taxa needed for these analyses). *Linum bienne* was represented by southwestern and southeastern individuals assigned to the same taxon. (A) Network likelihoods increased when one or two hybrid nodes were added to the backbone tree (derived by pruning the ASTRAL tree), and likelihoods were higher when *L. bienne* was specified as the hybrid taxon than when hybrid nodes were inferred freely. (B–E) Representative network topologies inferred when *L. bienne* was specified as the hybrid taxon. Black indicates the backbone tree, while red indicates hybrid network edges. Hybrid edges are thicker if y > 0.1 (major contribution to hybrid node), while they are thinner if y < 0.1 (minor contribution/spurious). (B-C) show network topologies with the highest likelihood for one (B) and two (C) hybrid nodes, although they both contain minor hybrid edges (y < 0.1). Thus, the top 10 alternative network topologies were explored. with (D-E) show topologies for one or two hybrid nodes respectively for which the hybrid edges leading to *L. bienne* both gave a contribution greater than 20% (y > 0.1).

Penalized likelihood dating of the supermatrix and Astral nuclear trees and the plastid tree sometimes also resulted in conflicting outcomes, as highlighted in Figure S3A-C and Table 1, where dating of nodes of interest in *Linum* are reported for the three phylogenetic trees. Note that the plastid tree did not contain all species given data quality (see Methods). In brief, across datasets, the age of Clade 1 was more variable, possibly because represented by a sparse number of species, while the age of Clade 2 was consistent across datasets (Table 1). The diversification of Clade 2 occurred mostly throughout the Miocene (Figure S3A-C). Within this clade, the *L. bienne* crown node was dated consistently across datasets (15-18 My), also within the range of previous reports (Table 1). For the crown node of *L. hologynum + L.marginale* + *L. bienne + L. usitatissimum* the age range retrieved (between 8.5 and 14 My, towards the end of the Miocene) was instead quite large and variable, although within the range from previous studies (Table 1). Finally, *L. bienne* + *L. usitatissimum* was estimated to diverge from all other species between the early Pliocene (4.5 Mya, nuclear supermatrix tree) and the Pleistocene (0.4 Mya; nuclear Astral tree). The older estimate (4.5Mya) apparently exceeded previous reports (Table 1). However, it was consistent with previous work when considering the age of the *L. villarianum + L. bienne* + *L. usitatissimum* crown node (Figure S3A).

**Table 1.** Summary of treePL dating (penalized likelihood) of the best maximum likelihood phylogenetic trees (plastid and nuclear Angiosperms353 supermatrix) and Astral tree (nuclear Angiosperms353 genes). The same fossil calibration was applied to the crown node of Linum in all analyses. At genus levels, we report the estimated crown node age for the main two *Linum* clades and for our node of interest (see Table S1 for definition of groups). Within the *L. bienne* node, we report the crown node age for *L. bienne* and *L. usitatissimum* and their sister species, *L. hologynum* and *L. marginale*, as well as for *L. bienne* and *L. usitatissimum* only. For this node, we also report the range of estimates from previous studies for comparison.

| Level | Crown Node | Divergence Time (Mya) |  |  |  |
| --- | --- | --- | --- | --- | --- |
|  |  | Plastid<br>(Supermatrix) | Nuclear<br>(Supermatrix) | Nuclear (Astral) | Previous<br>Reports* |
| *Linum* | Clade 1 | 12.5 | 19 | 7 | - |
|  | Clade 2 | 29 | 27 | 33 | - |
|  | *L. bienne* node | 18 | 15 | 18 | 15-20 |
| *L. bienne* node | *L. hologynum* +<br>*L. marginale* + *L.*<br>*biene* + *L.*<br>*usitatissimum* | 8.5 | 10 | 14 | 8-19 |
|  | *L. bienne* + *L.*<br>*usitatissimum* | 2.5 | 4.5 | 0.4 | 1-4 |

\* see Bolsheva et al. (2022), Maguilla et al. (2022), Villalvazo-Hernandez et al. (2022)

Finally, the genus level analysis also highlighted conflicting phylogenetic signal at intraspecific level between *L. bienne* and *L. usitatissimum*, with *L. bienne* individuals of different geographic origin included to represent the species distribution changing position relative to *L. usitatissimum* and low bootstrap support for internal nodes within this group for nuclear trees (Figure 1A-C; Figure S1A-B). Both the plastid and Astral nuclear phylogenetic trees consistently placed the *L. bienne* southwestern individual as sister to the southeastern and northwestern individuals (Figure 1A and 1C). However, the *L. usitatissimum* individual was placed as sister to the southwestern individual in the plastid phylogenetic tree, and as sister to the northwestern individual in the nuclear ASTRAL tree. The supermatrix nuclear tree (Figure 1B) had instead the northwestern *L. bienne* individual as sister to all other *L. bienne* and *L. usitatissimum* individuals, and *L. usitatissimum* in turn sister to the southeastern and northwestern *L. bienne* individual. The densitrees based on 1000 bootstrap trees for the plastid and nuclear supermatrix phylogenetic trees confirmed conflict at this node, with most conflict occurring due to the placement of *L. usitatissimum* (Figure S1A-B).

### Range-wide population genomics, plastome variation, and demographic history of *L. bienne*

Both the plastome phylogenetic tree (best ML tree from IQTREE) and the nuclear genetic distance tree (ngsDist) consistently identified four major *L. bienne* lineages with partitioned geographical distribution and high bootstrap support (Figure 3A; Figure S4A-B). The four *L. bienne* lineages included individuals from: (1) the western Mediterranean and the Canary Islands (SW lineage); (2) the centre of the Mediterranean (Central lineage); (3) the eastern Mediterranean (SE lineage); (4) northwestern Europe (NW lineage). The SW (southwestern) and Central lineages formed a clade sister to the SE (southeastern) and NW (northwestern) lineages. The NW lineage was nested within the SE lineage in the nuclear tree, corroborating patterns seen in the genus level analysis (Figure 1A and 1C). Younger nodes within the SE–NW lineage in the plastome tree had reduced bootstrap support (< 90%; Figure S4A-B). PCA and ngsADMIX analyses also corroborated this structure (Figure 3A; Figure S6A-B). The ancillary phylogenetic tree based on a small subset of plastid markers further placed *L. villarianum* as sister to *L. bienne*, and not as part of any of the *L. bienne* lineages, although bootstrap support (64%) was low (Figure S5).

**Figure 3.**
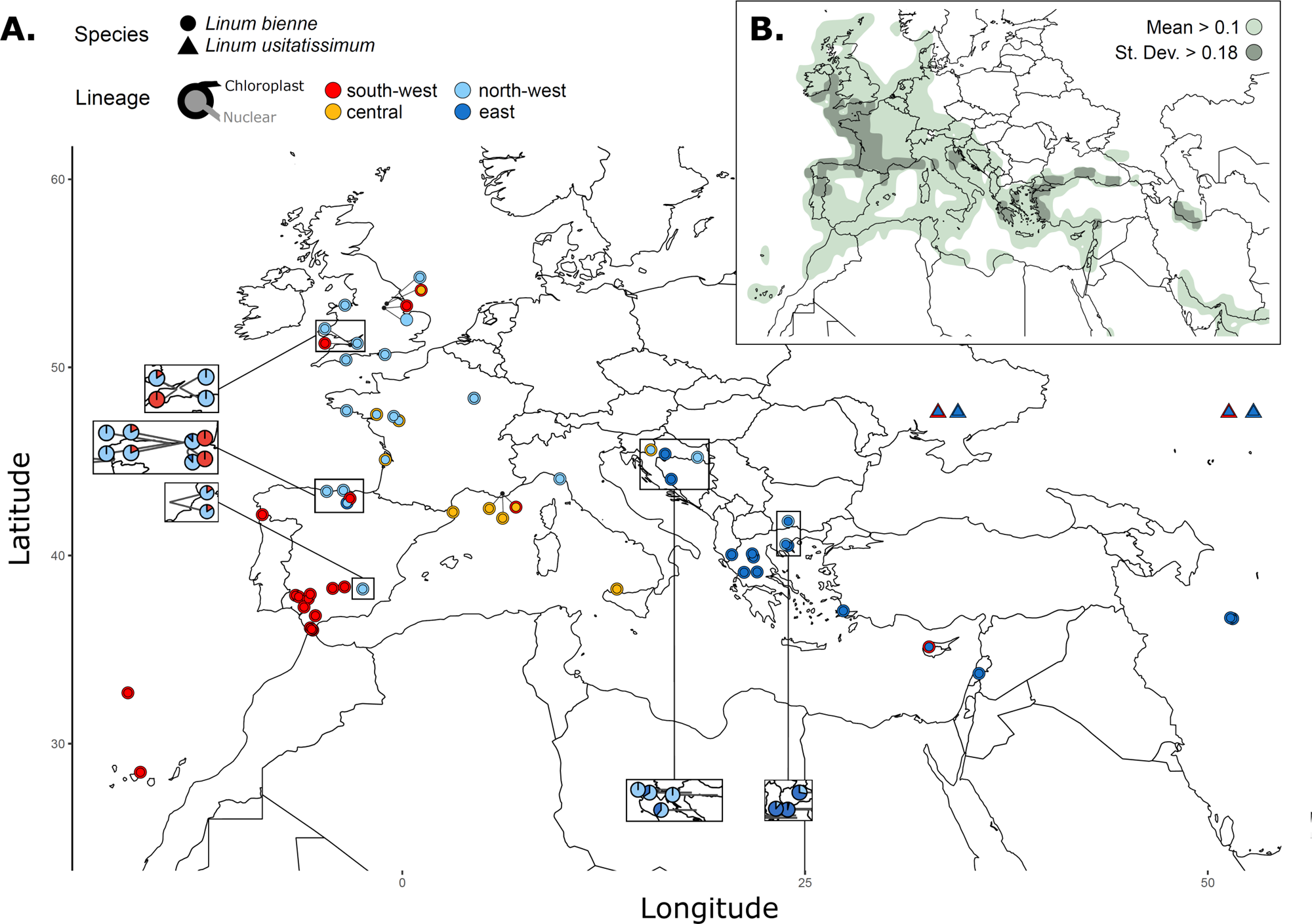
Geographic structure, cytonuclear discordance, and climatic favourability across the range of *Linum bienne*. (A) Geographic distribution of *L. bienne* lineages inferred from plastid and nuclear data. Circles represent *L. bienne* individuals and triangles represent *L. usitatissimum* individuals. Outer colours show lineage assignment based on plastome phylogenetic analyses, whereas inner colours show lineage assignment based on nuclear genetic distances. Differences between inner and outer colours indicate cytonuclear discordance. Insets show nuclear ancestry proportions inferred with ngsADMIX for regions where accessions showed mixed ancestry or discordant plastid and nuclear assignments. (B) Mean climatic favourability for *L. bienne* across the nine climatic periods analysed, from the early Pliocene to the present, estimated using GAM environmental niche modelling. Lighter shading indicates areas with mean favourability above 0.1. Darker shading indicates regions where favourability varied strongly across time periods, defined as standard deviation greater than 0.18. Regions of cytonuclear discordance and mixed ancestry spatially overlap with areas of high temporal variability in climatic favourability.

Despite clear spatial structuring of lineages across the range, several individuals were assigned to lineages inconsistent with their geographic location (Figure 3A; S4A-4B; FigureS6A). Some individuals from northern Spain and England grouped with the SW lineage, and one individual from southern Spain grouped with the NW lineage. Moreover, frequent cytonuclear discordance indicated chloroplast capture events, the most prominent involving Central vs. NW lineages (Figure 3A; Figure S4A-B): one individual from Croatia (L87) and six individuals from Atlantic France (L04, L05, L47–L50) carried Central plastid haplotypes but belonged to the NW lineage in the nuclear tree. Another example of cytonuclear discordance was within the SE–NW group: the nuclear tree included individuals from the SE clearly basal to NW individuals (except for L85 from Croatia), whereas plastid data did not clearly separate SE and NW individuals. These discordances were also reflected in ngsADMIX assignments (Figure 3A; Figure S6A-B), identifying individuals with mixed ancestry in northern Spain, England, northern Croatia, and Bulgaria. Overall, the NW lineage showed signs of cytonuclear discordance with both the SE and Central lineages, but ngsADMIX revealed multi-ancestry signal for SE-NW individuals only. Finally, *L. usitatissimum* also displayed cytonuclear discordance (Figure 3A; Figure S4A-B). It was monophyletic in the nuclear tree, but paraphyletic in the plastid tree, always displaying high bootstrap support. In the nuclear phylogenetic tree, all *L. usitatissimum* individuals were nested within the SE lineage (with one *L. bienne* individual from Cyprus, L91). In the plastid tree, two *L. usitatissimum* individuals (L71–72) grouped with SE *L. bienne* (Croatia, Greece), while the remaining individuals (cultivar, L70, L73–74) clustered with SW *L. bienne*, again with L91. Similar to the nuclear phylogenetic tree, *L. usitatissimum* finally formed a distinct genetic cluster based on ngsADMIX, although closer to SE and NW groups than others (Figure S6B). This aligns with the conflicting signal concerning *L.bienne - L. usitatissimum* found in the genus-level analysis (Figure 1A-C; Figure S1A-B).

Overall, the above analyses revealed gene flow between geographically partitioned *L. bienne* lineages, despite high selfing reported for its cultivated relative (Gutaker et al., 2019; Jhala et al., 2011). Within this context, pairwise genetic distances captured decreasing genetic diversity towards northern latitudes, both within the SW and the SE+NW lineages (Figure 4A). Thus, we proceeded with inferring the age of the main lineages, the time of possible expansions and contractions in population size, and secondary contact events between lineages. Overall, the origin and diversification of *L. bienne* was placed across Pliocene and Pleistocene, when climate progressively became more Mediterranean with enhanced seasonality, followed by ice-age cycles (Bache et al., 2011; Suc et al., 2018). Older divergence and diversification estimates derived from penalized likelihood dating of the nuclear or plastid supermatrix trees (Figure S3A-B) and BEAST plastome dating based on secondary calibration points derived from such trees (Figure S7A-B), while younger estimates derived from penalized likelihood dating of the nuclear Astral tree (Figure S3C), relative BEAST plastome dating (Figure S7C), as well as nuclear-based demographic analyses (Figure 4B).

**Figure 4.**
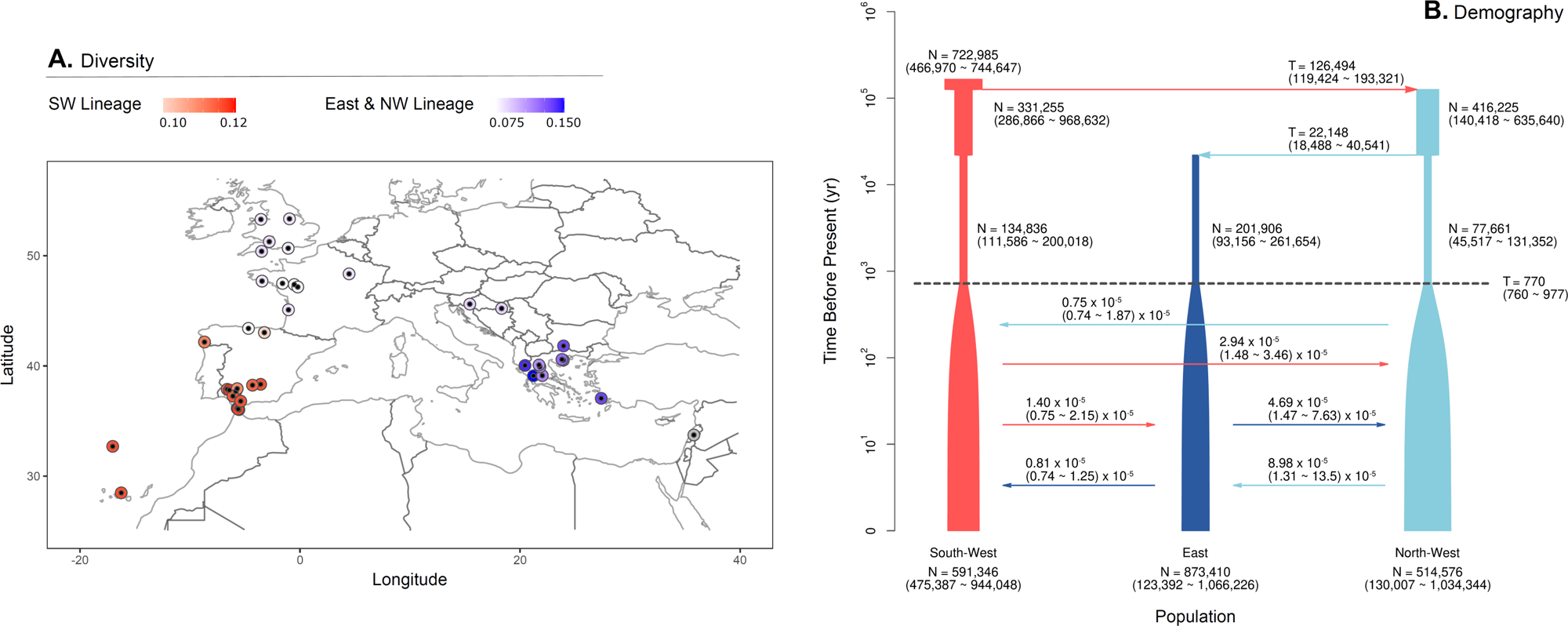
Genetic diversity and demographic history of *Linum bienne* lineages. (A) Spatial patterns of nuclear genetic diversity within the SW and SE–NW genetic clusters. Genetic diversity was estimated from pairwise nuclear genetic distances after excluding individuals with low assignment to the focal clusters and excluding the Central lineage because of its small sample size. Genetic diversity declines from south to north, particularly within the SE–NW cluster. (B) Demographic history inferred with fastsimcoal2.8 under the secondary-contact model. Colours indicate the SW, SE and NW lineages. T indicates divergence or demographic event times, and N indicates effective population size. The model supports lineage divergence during the late Pleistocene, post-split population contractions and subsequent expansions, with secondary contact among lineages.

More in detail, BEAST dating of the *L. bienne* plastome tree based on plastid (supermatrix) or nuclear (supermatrix, astral) secondary calibrations (Figure S7A-C) converged successfully (ESS > 200 for most parameters; Figure S8A-C), placing major splits in the late Miocene-Pliocene, soon after the reflooding of the Mediterranean (Bache et al., 2011), up to the early Pleistocene. The nuclear supermatrix calibration (Figure S7B) resulted in the *L. bienne* crown node at ∼5.4 My (95% HPD: 3.9–7.1 My). The SW+Central and SE+NW crown nodes were ∼4.1–4.4 My (95% HPD SW+Central = 2.8–6.2 My; SE+NW = 2.5–6.0 My), with SW and Central lineages at ∼3.1–3.7 My and the younger NW lineage at ∼1.8 Ma (95% HPD: 0.9–3.0 My). Similar results were obtained based on plastid supermatrix calibrations (Figure S7A). Conversely, secondary calibrations based on the nuclear Astral tree (Figure S7C) yielded node ages roughly 10 times younger and larger 95% HPD intervals, placing most events in the Pleistocene while recovering an anomalously young *L. narbonense*–*L. bienne* crown node (0.76 My, 95% HPD: 0.01–2.24 My) relative to other analyses (Figure S3).

Demographic modelling of nuclear data in fastsimcoal2 aligned with younger estimates for *L. bienne* divergence and diversification (Figure 4B), placed during the Pleistocene. Note that we calibrated fastsimcoal2 divergence estimates by assigning an average modal divergence of 0.244 substitutions per site (±0.002 s.d., N=4 genome pairs; see Supplementary Information) to the divergence time of 37 Mya, as estimated in our genus level analysis (Figure S3). Based on this, the secondary contact (SC) scenario had the highest maximum likelihood (ML estimated −292749.83, observed = −280466.505) compared to SI (estimated = −296715.23) or IM (estimated = −298432.44). The SC scenario supported an expansion with gene flow across all three lineages ∼770 ka, followed by an initial split (SW vs. SE+NW) ∼126 ka and a second split (SE vs. NW) ∼22 ka. All lineages exhibited post-split contractions (∼70,000–200,000 individuals) before re-expanding to ∼500,000–800,000 individuals (Figure 4B). Additionally, to capture complex dynamics and resolve the Central lineage, MSMC2 and MSMC-IM were also applied (Figure S9). These analyses supported a relatively older separation of SW+Central from SE+NW ∼400 ka, with cross-coalescence declining after ∼80 ka. Divergence of SE and NW occurred ∼40–5 ka with high persistent migration. Nuclear ngsADMIX multi-ancestry patterns (Figure S6) and cytonuclear discordance (Figures 3A, S4A-B) confirm recent gene flow or incomplete divergence among NW–SW, NW–SE, and NW–Central lineages. MSMC2 also revealed a pronounced bottleneck in all lineages between ∼60–10 ka, strongest in the NW lineage during its divergence from SE. Combined with the latitudinal decline in genetic diversity (Figure 4A), these dynamics indicate repeated range shifts during Pleistocene climate oscillations.

### The environmental niche of *L. bienne* from Pliocene to present

Based on the Pliocene-Pleistocene diversification timeline of *L. bienne*, ENM was conducted based on the present species distribution and the model was then used to predict where the species could likely occur in the past. MESS detected the highest climate dissimilarity between the training region (Iberian Peninsula and France) and north-eastern Europe (Figure S10D). Excluding this area, present predictions across the species range overlapped substantially across GLM, GAM, and Maxent, with GAM selected due to a higher TSS (Figure S10E) and predictions were also calculated for past climates. When averaging across present and past predictions, mean favourability exceeded 0.1 across Europe and the Mediterranean Basin (Figures S3B, S11A). Within this range, high variability (SD > 0.18) in the northern Iberian Peninsula, Atlantic France, and along the Balkan and Turkish coasts marked these areas as climatically unstable over time. Instead, low variability areas (SD < 0.18) such as the Mediterranean peninsulas, might have worked as stable refugia throughout *L. bienne* evolution (Figure 3B). In fact, from the early Pliocene to the present, favourability alternated dynamically, with higher latitudes being more variable (Figure 3B, S11). Early and mid-Pleistocene predictions—coinciding with the species’ inferred origin based on coalescent analyses of low-depth nuclear genomes—showed favourable conditions across the range that contracted toward the Mediterranean Basin around the mid-Pleistocene. This was followed by a post-LGM expansion of favourable climate across Europe during the mid-Holocene (∼8 ka), which remains similar to present-day distributions (Figure S11A-B).

## Discussion

### Conflicting phylogenetic signal across the genus *Linum* and the origin of *L. bienne*

*Linum* diversified during major Oligocene to Pliocene geological and climatic changes in the Mediterranean Basin, where repeated isolation and secondary contact have shaped many plant lineages (Cherchi & Montadert, 1982; Nieto Feliner, 2014). This temporal and geographical context coincides with a complex history of chromosome evolution and WGDs around the Miocene–Pliocene transition (Cai et al., 2019; Sveinsson et al., 2014; Wang et al., 2012; Valdés-Florido et al., 2023). In our dated plastid and nuclear phylogenies (Figure S3A-C), several deeper nodes fall within this window, suggesting that genome duplication formed part of the broader evolutionary backdrop to lineage diversification in *Linum*. Reticulation has long been considered an important component of the evolutionary history of the genus (Fu & Allaby, 2010), whereas WGD does not alone explain diversification patterns in the genus (Valdés-Florido et al., 2023).

Our phylogenomic analyses helped distinguish the evolutionary mechanisms underlying phylogenetic discordance in *Linum*. We identified conflicting nodes across the genus, with discordance being more frequent at shallower nodes (Figure 1A-C; Figure S1 and S2) and at nodes characterized by WGD or high chromosome numbers (Valdes et al., 2023; Cai et al., 2019; Bolsheva et al., 2017, 2022; Sveinsson et al. 2014; Wang et al., 2012; Ockendon, 1968; Table S1). Together, these results indicate that phylogenetic discordance is widespread across our genus-level sampling, with much of this discordance consistent with ILS.

At the node including *L. bienne*, cultivated flax, and their closest relatives, phylogenetic discordance involved an additional signal of gene flow acting together with ILS. Branch length and network analyses indicated that discordance at this node cannot be explained by ILS alone, showing that the origin of *L. bienne* and close relatives was not strictly bifurcating (Figure 2; Figure S2B). This supports previous hypotheses of *L. bienne* and *L. usitatissimum* being meso-allopolyploids (Bolsheva et al., 2017; Wang et al., 2012). Although cytonuclear discordance is often expected under hybridization, plastid and nuclear trees agreed at the *L. bienne* node (Figure 1). However, this agreement does not rule out reticulation. ML network analyses suggest that hybridizing ancestors of *L. bienne* had already diverged from *L. narbonense* and were more closely related to the *L. decumbens* + *L. grandiflorum* lineage (Bolsheva et al., 2017). This could explain why reticulation was detected via network analyses despite the lack of an obvious plastid-nuclear topology swap (Figure 2D-E). Furthermore, many plastid genes supported an alternative topology placing *L. narbonense* closer to *L. bienne*+*L. marginale* than *L. decumbens*+*L. grandiflorum* (Table S5). This marker-specific signal, alongside ITS marker variation, likely explains the swapped topologies recovered in previous phylogenetic studies (Table S6).

Topological discrepancies among previous studies were not restricted to the *L. bienne* node. Our study comparison showed recurrent variation within several sections of the genus, particularly between studies using few plastid or ITS markers and those using broader nuclear or plastome datasets (Table S6). Generally, extensive genome sampling retrieved similar topologies, whereas fewer markers yielded more variable topologies at different nodes (Table S6). This suggests locus-specific signal and limited marker sampling may contribute to past discrepancies, as supported by our finding that individual plastid genes recover alternative topologies at the *L. bienne* node (Table S5). Crucially, topological variation alone should not be interpreted as evidence of gene flow. In our analyses, support for gene flow was concentrated at the *L. bienne* node, whereas we did not find strong evidence of gene flow across other sections in *Linum* (Figure S2B). However, outside this focal clade, our taxon sampling was not designed to test reticulation exhaustively, especially given the pervasive genus-wide ILS (Figure S2A-B). Broader taxonomic and genomic sampling remains necessary to evaluate how chromosome evolution, WGD, ILS, and reticulation interact across other *Linum* sections (Tank et al., 2015; Ren et al., 2018; Huynh et al., 2019; Murillo et al., 2022; He et al., 2024; Xu et al., 2024; Jin et al., 2025; Zhao et al., 2025; Sun et al., 2026).

The phylogenetic complexity is also central to interpreting the differences in divergence time estimates for the crown node of *L. bienne* and other nodes (Figure S3; Figure S7). Rather than a single discrete speciation event, these estimates may capture different stages in the species’ evolutionary history. Our supermatrix nuclear phylogeny and plastid-based dating yielded older, late Miocene to Pliocene estimates, whereas coalescent nuclear analyses and demographic inference pointed to a younger Pleistocene diversification of extant *L. bienne* lineages. Discrepancies among nuclear, plastid or mitochondrial markers are not uncommon in complex lineages (Wahlberg et al., 2009; Huang et al., 2014; Middleton et al., 2014; Xu et al., 2021; Pokorny et al., 2024). Here, evidence for ILS and reticulation suggests that different datasets and dating approaches may be capturing different depths of the evolutionary history of *L. bienne*. Older estimates likely reflect the deep history of an ancestral reticulate lineage—including *L. bienne*, *L. usitatissimum*, and potentially the North African *L. villarianum* (Ruiz-Martín et al., 2018; Maguilla et al., 2021; Villalvazo-Hernández et al., 2022)—originating during Late Miocene climatic and sea-level changes (Bache et al., 2011; Nieto Feliner, 2014; Suc et al., 2018). This lineage might have originated from hybridization and WGD previously described for *L. bienne* and *L. usitatissimum* (Bolsheva et al., 2017; Wang et al. 2012). Conversely, the younger estimates based on coalescence better reflect divergence and subsequent diversification of *L. bienne* during the Pleistocene, an interpretation supported by environmental niche modeling showing that climatic favourability for *L. bienne* persisted through the Pleistocene despite areas characterized by higher variability (Figure S11; see discussion below).

### The diversification of *L. bienne* into geographically partitioned lineages with secondary contact

The diversification of *L. bienne* into four geographically structured lineages reflects population differentiation and range shifts during Pleistocene climatic oscillations which strongly shaped European flora (Hewitt, 1999; Nieto Feliner, 2014; Kadereit & Abbott, 2021). The SW + Central and SE + NW groups suggest differentiation within at least two major refugia: the western Mediterranean, and the Balkans/Anatolia, respectively. The distinct Central Mediterranean lineage may indicate an additional area of persistence, though confirming this requires deeper sampling across Mediterranean islands and the Apennine Peninsula. These regions are classic long-term diversity reservoirs for Mediterranean floras (Médail & Diadema, 2009), with comparable longitudinal genetic structuring reported for other Mediterranean plant species (Baumen et al., 2022; Campos et al., 2023). The diversification of *L. bienne* appears to have occurred in response to successive glacial-interglacial cycles interacting with complex Mediterranean orography. Demographic analyses placed the divergence of the SW lineage during the Last Interglacial, whereas the NW lineage diverged from the SE lineage during the Last Glacial Maximum (Figure 4B and S9). This asynchronous divergence shows that lineage splitting tracked major climatic transitions. Furthermore, the south-to-north decline in genetic diversity (Figure 4A) matches postglacial expansions following latitudinal range contractions (Excoffier et al., 2009). Environmental niche modelling supports this, showing that southwestern regions of the Mediterranean peninsulas and northwest Africa remained relatively favourable for *L. bienne* across climatic periods, whereas northern regions experienced stronger fluctuations in suitability (Figure 2B and S11).

Despite the clear geographic structure, divergence among *L. bienne* lineages was not strictly allopatric. Although plastid and nuclear trees agreed at genus level, cytonuclear discordance was evident at intraspecific level in *L. bienne*. Several individuals carried plastid haplotypes from one lineage but mapped to another in the nuclear tree, with some individuals showing admixed ancestry in *ngsADMIX* (Figure 3A; Figure S4A-B). These patterns point to secondary contact and gene flow among previously diverged lineages. ENM indicates that climatically favourable areas expanded northwards during the Holocene, within the last 10,000 years (Figure S11). This expansion matches demographic inferences of gene flow between the NW lineage and the SW and Central lineages (Figure S9), suggesting that postglacial range expansions facilitated secondary contact. Additionally, the concordant plastid and nuclear assignment of certain individuals to lineages far outside their typical geographic range (e.g., southern England; Figure 3A) suggests more recent long-distance dispersal events hypothesized in *Arabidopsis thaliana* (Alonso-Blanco et al., 2016).

Secondary contact likely had consequences beyond neutral demographic mixing. Populations within the NW lineage of *L. bienne* frequently require vernalization to flower (Landoni et al., 2024), and previous work suggests transfer of flowering-time alleles between northwestern and southeastern Europe (Gutaker et al. 2019). In other systems, introgression during postglacial expansions has been shown to facilitate adaptation, particularly through the transfer of alleles conferring tolerance to novel or extreme conditions (Lovell et al., 2021). Although our data do not directly test for adaptive introgression, gene flow among *L. bienne* lineages may have contributed to introduce adaptive variation during range expansions into climatically distinct parts of its range. The similar ecology and evolutionary history of *L. lewisii*, a close North American relative (Innes et al., 2025), suggests that comparative studies across *Linum* could yield valuable insights into parallel evolutionary responses to environmental gradients.

### Conflicting phylogenetic signal affects *L. bienne* and cultivated flax

Domestication is increasingly recognized as a complex evolutionary process, involving recurrent gene flow between crops and their wild relatives (Purugganan, 2019). To place cultivated flax within the broader evolutionary history of *L. bienne*, we included several *L. usitatissimum* accessions in our analyses. In the nuclear phylogeny, all crop accessions clustered together with a Cypriot *L. bienne* individual within the SE lineage (Figure S4), supporting an eastern Mediterranean origin, likely near present-day Turkey (Gutaker et al., 2019; Weiss & Zohary, 2011). The phylogenetic placement of the crop is informative considering the recent demographic history of *L. bienne*. The SE–NW lineage likely expanded northwards during the Holocene, within the last 10,000 years (Figure 4), a timeframe broadly overlapping with flax domestication (Gutaker et al., 2019). This temporal alignment raises the possibility that early domesticated flax expanded in parallel and in continued contact with wild *L. bienne* populations. This consistent link matches evidence that introgression from *L. bienne* facilitated crop adaptation to northern latitudes (Gutaker et al., 2019). Despite our limited crop sampling, these results suggest the early spread of cultivated flax was closely linked to its wild relative’s expansion.

Given the history of interaction, the cytonuclear discordance in *L. usitatissimum* is expected. Nuclear data placed all crop individuals within the SE lineage, whereas plastid phylogenies revealed that some *L. usitatissimum* accessions carry haplotypes associated with the SW lineage (Figure S4). Because the SW and SE lineages diverged early (Figure 4; Figure S9), this conflict cannot be explained by ILS alone. Assuming a single domestication center in the eastern Mediterranean (Allaby et al., 2005), this discordance likely points to post-domestication gene flow with geographically differentiated wild lineages. Alternatively, a lineage closely related to the SW group may have historically occupied the domestication region. Distinguishing these scenarios requires denser geographic sampling of wild and crop populations, especially near early agricultural contact zones.

In conclusion, the evolutionary history of *L. bienne* cannot be understood from a strictly bifurcating perspective. The *L. bienne* node retains distinct signatures of hybridization and ILS overlaid across a broader history of genome duplication. At shallower timescales, intraspecific *L. bienne* lineages diversified primarily during the Pleistocene, when climatic oscillations promoted persistence in southern refugia, northward expansions, and repeated secondary contact. Consequently, the contemporary geographic structure of *L. bienne* reflects both historical isolation and recurrent gene flow. This combination of a deep reticulate history and recent phylogeographic differentiation explains why the species harbours substantial genetic diversity, which is directly relevant for flax crop improvement and conservation under climate change (Fu, 2023). More broadly, our results demonstrate how evolutionary complexity in *Linum* emerges from the dynamic interplay between lineage divergence, reticulation, and climate-driven range shifts across multiple timescales.

## Acknowledgements

RPB was supported by an IBBS Research Grant from the University of Portsmouth (UoP) and by the Spanish Ministry of Science and Innovation (grant PID2021-127264NB-I00 funded by MCIN/AEI/10.13039/501100011033 ERDF A way of making Europe). BL was supported by her PhD bursary provided by the UoP and funding from the Botanical Society of the British Isles (BSBI) to conduct sequencing for three high depth *L. bienne* genomes. JV was supported by RYC2023-042611-I by MCIU/AEI/10.13039/501100011033 and FSE+. We also thank the authors of Villalvazo-Hernandez et al. (2022) for clarifying details about their analyses.

## Author Contributions

RPB, JV, and BL devised the study and experimental design. Some of the French and UK samples of *L. bienne* were provided by ACB. Eastern *L. bienne* and *L. usitatissimum* samples were provided by RGA. The DNA extractions and preparation of genomic libraries were conducted by JV. All genome sequencing was outsourced from external companies. BL and JV designed the pipeline to retrieve the plastid genomes. YB designed the pipeline to retrieve low-depth nuclear data covering the whole nuclear genome of *L. bienne* and conducted the corresponding demographic analyses. BL conducted all other analyses and produced all the figures. BL and RPB wrote the manuscript. All authors reviewed the manuscript afterwards.

## Code & Data Availability

All scripts are available at https://github.com/YannBourgeois/Scripts_Linum (*L. bienne* nuclear genome analyses) and https://github.com/beaLando/linomics.git (all other analyses). All genomic data were deposited in ENA under project name PRJEB86756 which will be publicly available upon publication of the manuscript.

## REFERENCES

Abadi, S., Azouri, D., Pupko, T. et al. (2019). Model selection may not be a mandatory step for phylogeny reconstruction. Nature Commununications 10, 934. 10.1038/s41467-019-08822-w

Ana Afonso, João Loureiro, Juan Arroyo, Erika Olmedo-Vicente, Sílvia Castro, Cytogenetic diversity in the polyploid complex Linum suffruticosum s.l. (Linaceae), Botanical Journal of the Linnean Society, Volume 195, Issue 2, February 2021, Pages 216–232, 10.1093/botlinnean/boaa060

Afonso A, Castro S, Loureiro J, Arroyo J, Figueiredo A, Lopes S and Castro M (2023) Ecological niches in the polyploid complex Linum suffruticosum s.l.. Front. Plant Sci. 14:1148828. doi: 10.3389/fpls.2023.1148828

Allaby, R. G., Peterson, G. W., Merriwether, D. A., & Fu, Y.-B. (2005). Evidence of the domestication history of flax (Linum usitatissimum L.) from genetic diversity of the sad2 locus. Theoretical and Applied Genetics, 112(1), 58–65. 10.1007/s00122-005-0103-3

Alonso-Blanco, C., Andrade, J., Becker, C., Bemm, F., Bergelson, J., Borgwardt, K. M.,… & Zhou, X. (2016). 1,135 genomes reveal the global pattern of polymorphism in Arabidopsis thaliana. Cell, 166(2), 481–491.

Andrews, S. (2010). FastQC: a quality control tool for high throughput sequence data. http://www.bioinformatics.babraham.ac.uk/projects/fastqc/

Ansell, S. W., Stenøien, H. K., Grundmann, M., Schneider, H., Hemp, A., Bauer, N., Russell, S. J., & Vogel, J. C. (2010). Population structure and historical biogeography of European Arabidopsis lyrata. Heredity, 105(6), 543–553. 10.1038/hdy.2010.10

Armbruster, W. S., Pérez-Barrales, R., Arroyo, J., Edwards, M. E., & Vargas, P. (2006). Three-dimensional reciprocity of floral morphs in wild flax (Linum suffruticosum): A new twist on heterostyly. New Phytologist, 171(3), 581–590. 10.1111/j.1469-8137.2006.01749.x

Bache, F., Popescu, S.-M., Rabineau, M., Gorini, C., Suc, J.-P., Clauzon, G., Olivet, J.-L., Rubino, J.-L., Melinte-Dobrinescu, M.C., Estrada, F., Londeix, L., Armijo, R., Meyer, B., Jolivet, L., Jouannic, G., Leroux, E., Aslanian, D., Reis, A.T.D., Mocochain, L., Dumurdžanov, N., Zagorchev, I., Lesić, V., Tomić, D., Namık Çağatay, M., Brun, J.-P., Sokoutis, D., Csato, I., Ucarkus, G. and Çakır, Z. (2012), A two-step process for the reflooding of the Mediterranean after the Messinian Salinity Crisis. Basin Research 24, 125–153. 10.1111/j.1365-2117.2011.00521.x

Baker, W. J., Bailey, P., Barber, V., Barker, A., Bellot, S., Bishop, D., Botigué, L. R., Brewer, G., Carruthers, T., Clarkson, J. J., Cook, J., Cowan, R. S., Dodsworth, S., Epitawalage, N., Françoso, E., Gallego, B., Johnson, M. G., Kim, J. T., Leempoel, K.,… Forest, F. (2022). A Comprehensive Phylogenomic Platform for Exploring the Angiosperm Tree of Life. Systematic Biology, 71(2), 301–319. 10.1093/sysbio/syab035

Barreto, E., Holden, P. B., Edwards, N. R., & Rangel, T. F. (2023). PALEO-PGEM-Series: A spatial time series of the global climate over the last 5 million years (Plio-Pleistocene). Global Ecology and Biogeography, 32, 1034–1045, doi:10.1111/geb.13683

Baumel, A., Nieto Feliner, G., Médail, F., La Malfa, S., Di Guardo, M., Bou Dagher Kharrat, M., Lakhal-Mirleau, F., Frelon, V., Ouahmane, L., Diadema, K., Sanguin, H., & Viruel, J. (2022). Genome-wide footprints in the carob tree (Ceratonia siliqua) unveil a new domestication pattern of a fruit tree in the Mediterranean. Molecular Ecology, 31, 4095–4111. 10.1111/mec.16563

Bolsheva, N. L., Melnikova, N. V., Kirov, I. V., Speranskaya, A. S., Krinitsina, A. A., Dmitriev, A. A., Belenikin, M. S., Krasnov, G. S., Lakunina, V. A., Snezhkina, A. V., Rozhmina, T. A., Samatadze, T. E., Yurkevich, O. Yu., Zoshchuk, S. A., Amosova, А. V., Kudryavtseva, A. V., & Muravenko, O. V. (2017). Evolution of blue-flowered species of genus Linum based on high-throughput sequencing of ribosomal RNA genes. BMC Evolutionary Biology, 17(S2), 253. 10.1186/s12862-017-1105-x

Bolsheva, N. L., Melnikova, N. V., Dvorianinova, E. M., Mironova, L. N., Yurkevich, O. Y., Amosova, A. V., Krasnov, G. S., Dmitriev, A. A., & Muravenko, O. V. (2022). Clarification of the Position of Linum stelleroides Planch. within the Phylogeny of the Genus Linum L. Plants, 11(5), 652. 10.3390/plants11050652

Brown, J. W., Walker, J. F., & Smith, S. A. (2017). Phyx: Phylogenetic tools for unix. Bioinformatics, 33(12), 1886–1888. 10.1093/bioinformatics/btx063

Burgarella, C., Barnaud, A., Kane, N. A., Jankowski, F., Scarcelli, N., Billot, C., Vigouroux, Y., Berthouly-Salazar C. (2019). Adaptive Introgression: An Untapped Evolutionary Mechanism for Crop Adaptation. Frontiers in Plant Science 10. 10.3389/fpls.2019.00004

Miguel Campos, Emma Kelley, Barbara Gravendeel, Frédéric Médail, J M Maarten Christenhusz, Michael F Fay, Pilar Catalán, Ilia J Leitch, Félix Forest, Paul Wilkin, Juan Viruel, Genomic, spatial and morphometric data for discrimination of four species in the Mediterranean Tamus clade of yams (Dioscorea, Dioscoreaceae), Annals of Botany, Volume 131, Issue 4, 14 March 2023, Pages 635–654, 10.1093/aob/mcad018

Cai, L., Xi, Z., Amorim, A. M., Sugumaran, M., Rest, J. S., Liu, L., & Davis, C. C. (2019). Widespread ancient whole-genome duplications in Malpighiales coincide with Eocene global climatic upheaval. New Phytologist, 221(1), 565–576. 10.1111/nph.15357

Cai, L., Xi, Z., Lemmon, E. M., Lemmon, A. R., Mast, A., Buddenhagen, C. E., Liu, L. & Davis, C. C. (2021). The Perfect Storm: Gene Tree Estimation Error, Incomplete Lineage Sorting, and Ancient Gene Flow Explain the Most Recalcitrant Ancient Angiosperm Clade, Malpighiales. Systematic Biology, 70 (3), 491–507. 10.1093/sysbio/syaa083

Capella-Gutierrez, S., Silla-Martinez, J. M., & Gabaldon, T. (2009). trimAl: A tool for automated alignment trimming in large-scale phylogenetic analyses. Bioinformatics, 25(15), 1972–1973. 10.1093/bioinformatics/btp348

Capistrano-Gossmann, G. G., Ries, D., Holtgräwe, D., Minoche, A., Kraft, T., Frerichmann, S. L. M., Rosleff Soerensen, T., Dohm, J. C., González, I., Schilhabel, M., Varrelmann, M., Tschoep, H., Uphoff, H., Schütze, K., Borchardt, D., Toerjek, O., Mechelke, W., Lein, J. C., Schechert, A. W.,… Kopisch-Obuch, F. J. (2017). Crop wild relative populations of Beta vulgaris allow direct mapping of agronomically important genes. Nature Communications, 8. 10.1038/ncomms15708

Cavagnetto, C., & Anadón, P. (1996). Preliminary palynological data on floristic and climatic changes during the Middle Eocene-Early Oligocene of the eastern Ebro Basin, northeast Spain. Review of Palaeobotany and Palynology, 92(3–4), 281–305. 10.1016/0034-6667(95)00096-8

Chamberlain S, Barve V, Mcglinn D, Oldoni D, Desmet P, Geffert L, Ram K (2025). rgbif: Interface to the Global Biodiversity Information Facility API. R package version 3.8.3, https://CRAN.R-project.org/package=rgbif.

Chauvet, Stéphanie, El Oualidi, Jalal, Fennane, Mohamed, Ibn Tattou, Mohamed, Khamar, Hamid, & Taleb, Mohamed Sghir. (2012). Checklist des endémiques et spécimens types de la flore vasculaire de l’Afrique du Nord. Rabat : Institut Scientifique, Universite Mohammed V - Agdal. https://www.biodiversitylibrary.org/bibliography/77150

Chen, S., Zhou, Y., Chen, Y., & Gu, J. (2018). fastp: an ultra-fast all-in-one FASTQ preprocessor. Bioinformatics, 34(17), i884–i890.

Chen, H., Almeida-Silva, F., Logghe, G., Maere, S., Bonte, D., Van de Peer, Y. (2024). The Rise of Polyploids During Environmental Upheaval. bioRxiv. 10.1101/2024.11.22.624806

Chen, X., Dadole, R., Avia, K., Venon, A., Brisson, M., Remoué, C.,… & Cornille, A. (2026). Gene flow from the European wild apple and selection shaped the domesticated apple genome. Current Biology, 36(8), 2104–2118.

Cherchi, A., & Montadert, L. (1982). Oligo-Miocene rift of Sardinia and the early history of the Western Mediterranean Basin. Nature, 298(5876), 736–739. 10.1038/298736a0

Comes, H. P., & Abbott, R. J. (2001). Molecular phylogeography, reticulation, and lineage sorting in mediterranean Senecio sect. Senecio (Asteraceae). Evolution, 55(10), 1943–1962. 10.1111/j.0014-3820.2001.tb01312.x

Corriveau, J. L., & Coleman, A. W. (2016). Rapid Screening Method to Detect Potential Biparental Inheritance of Plastid DNA and Results for Over 200 Angiosperm Species Author ( s ): Joseph L. Corriveau and Annette W. Coleman Published by: Botanical Society of America, Inc. Stable URL : *http:/*. 75(10), 1443–1458.

Cortés, A. J., Monserrate, F. A., Ramírez-Villegas, J., Madriñán, S., & Blair, M. W. (2013). Drought Tolerance in Wild Plant Populations: The Case of Common Beans (Phaseolus vulgaris L.). PLoS ONE, 8(5). 10.1371/journal.pone.0062898

Davies, R., Flint, J., Myers, S. et al. (2016). Rapid genotype imputation from sequence without reference panels. Nature Genetics 48, 965–969. 10.1038/ng.3594

de Santana Lopes, A., Pacheco, T.G., Santos, K.G.d., et al. (2018). The Linum usitatissimum L. plastome reveals atypical structural evolution, new editing sites, and the phylogenetic position of Linaceae within Malpighiales. Plant Cell Rep 37, 307–328. 10.1007/s00299-017-2231-z

Díaz-Pérez, A., López-Álvarez, D., Sancho, R., & Catalán, P. (2018). Reconstructing the origins and the biogeography of species’ genomes in the highly reticulate allopolyploid-rich model grass genus Brachypodium using minimum evolution, coalescence and maximum likelihood approaches. Molecular Phylogenetics and Evolution, 127(June), 256–271. 10.1016/j.ympev.2018.06.003

Diez, C.M., Trujillo, I., Martinez-Urdiroz, N., Barranco, D., Rallo, L., Marfil, P. and Gaut, B.S. (2015). Olive domestication and diversification in the Mediterranean Basin. New Phytologist 206. 436–447. 10.1111/nph.13181

Drummond, A. J., & Rambaut, A. (2007). BEAST: Bayesian evolutionary analysis by sampling trees. BMC Evolutionary Biology, 7(1), 214. 10.1186/1471-2148-7-214

Nathaniel B. Edelman et al. (2019). Genomic architecture and introgression shape a butterfly radiation. Science 366,594–599. DOI:10.1126/science.aaw2090

Edgar, R. C. (2004). MUSCLE: multiple sequence alignment with high accuracy and high throughput. Nucleic acids research, 32(5), 1792–1797.

Edwards, S. V., Robin, V. V., Ferrand, N., Moritz, C. (2022). The Evolution of Comparative Phylogeography: Putting the Geography (and More) into Comparative Population Genomics. Genome Biology and Evolution, 14(1). 10.1093/gbe/evab176

Excoffier, L., Foll, M., Petit, R. J. (2009). Genetic Consequences of Range Expansions. ANNUAL REVIEW OF ECOLOGY, EVOLUTION, AND SYSTEMATICS, 40, 481–501. 10.1146/annurev.ecolsys.39.110707.173414

Excoffier, L., Dupanloup, I., Huerta-Sánchez, E., Sousa, V. C., & Foll, M. (2013). Robust demographic inference from genomic and SNP data. PLoS genetics, 9(10), e1003905.

Fennane M. and Rejdali M., Aromatic and medicinal plants of Morocco: richness,diversity and threats, Bull. l’institut Sci. Rabat, Sect. Sci. la Vie. (2016) 38, 27–42.

Fick, S. E., & Hijmans, R. J. (2017). WorldClim 2: New 1-km spatial resolution climate surfaces for global land areas. International Journal of Climatology, 37(12), 4302–4315. 10.1002/JOC.5086

Fu, Y. B., & Allaby, R. G. (2010). Phylogenetic network of Linum species as revealed by non-coding chloroplast DNA sequences. Genetic Resources and Crop Evolution, 57(5), 667–677. 10.1007/s10722-009-9502-7

Fu, Y. B., Dong, Y., & Yang, M. H. (2016). Multiplexed shotgun sequencing reveals congruent three-genome phylogenetic signals for four botanical sections of the flax genus Linum. Molecular Phylogenetics and Evolution, 101, 122–132. 10.1016/j.ympev.2016.05.010

Fu, YB. (2023). Pale Flax (Linum Bienne): an Underexplored Flax Wild Relative. In: You, F.M., Fofana, B. (eds) The Flax Genome. Compendium of Plant Genomes. Springer, Cham. 10.1007/978-3-031-16061-5_3

García-Verdugo, C., Mairal, M., Tamaki, I. and Msanda, F. (2021), Phylogeography at the crossroad: Pleistocene range expansion throughout the Mediterranean and back-colonization from the Canary Islands in the legume Bituminaria bituminosa. J Biogeogr, 48: 1622–1634. 10.1111/jbi.14100

Grabowski, P. P., Dang, P., Jenkins, J. J., Sreedasyam, A., Webber, J., Lamb, M., Zhang, Q., Sanz-Saez, A., Feng, Y., Bunting, V., Talag, J., Clevenger, J., Ozias-Akins, P., Holbrook, C. C., Chu, Y., Grimwood, J., Schmutz, J., Chen, C., Lovell, J. T. (2024). Relics of interspecific hybridization retained in the genome of a drought-adapted peanut cultivar. G3 Genes|Genomes|Genetics, 14(11). 10.1093/g3journal/jkae208

Guggisberg, A., Mansion, G., Kelso, S., & Conti, E. (2006). Evolution of biogeographic patterns, ploidy levels, and breeding systems in a diploid–polyploid species complex of Primula. New Phytologist, 171(3), 617–632. 10.1111/j.1469-8137.2006.01722.x

Gutaker, R. M., Zaidem, M., Fu, Y., Diederichsen, A., Smith, O., Ware, R., & Allaby, R. G. (2019). Flax latitudinal adaptation at LuTFL1 altered architecture and promoted fiber production. Scientific Reports, 9(1), 976. 10.1038/s41598-018-37086-5

Gutiérrez-Valencia J, Fracassetti M, Berdan EL, Bunikis I, Soler L, Dainat J, Kutschera VE, Losvik A, Désamoré A, Hughes PW, Foroozani A, Laenen B, Pesquet E, Abdelaziz M, Pettersson OV, Nystedt B, Brennan AC, Arroyo J, Slotte T. Genomic analyses of the Linum distyly supergene reveal convergent evolution at the molecular level. Curr Biol. 2022 Oct 24;32(20):4360–4371.e6. doi: 10.1016/j.cub.2022.08.042

Hanghøj, K., Moltke, I., Andersen, P. A., Manica, A., Korneliussen, T. S. Fast and accurate relatedness estimation from high-throughput sequencing data in the presence of inbreeding. GigaScience, 2025, 8 (5), pp.giz034. 10.1093/gigascience/giz034

Harris, R. S. (2007). Improved pairwise alignment of genomic DNA. The Pennsylvania State University.

Guan-Hao He, Ying Meng, Meng-Hua Zhang, Da Wang, Ran Meng, Lei Zhang, Zhao-Fu Chu, Jun Wen, Ze-Long Nie, Extensive genome-wide phylogenetic discordance is due to incomplete lineage sorting in the rapidly radiated East Asian genus Nekemias (Vitaceae), Annals of Botany, Volume 135, Issue 5, 30 April 2025, Pages 925–934, 10.1093/aob/mcae224

Hewitt, G. M. (1999). Post-glacial re-colonization of European biota. Biological Journal of the Linnean Society, 68(1–2), 87–112. 10.1111/J.1095-8312.1999.TB01160.X

Holden, P. B., Edwards, N. R., Rangel, T. F., Pereira, E. B., Tran, G. T., and Wilkinson, R. D. (2019): PALEO-PGEM v1.0: a statistical emulator of Pliocene–Pleistocene climate, Geosci. Model Dev., 12, 5137–5155, doi:10.5194/gmd-12-5137-2019.

Hollister, J. D. (2015). Polyploidy: Adaptation to the genomic environment. New Phytologist, 205(3), 1034–1039. 10.1111/nph.12939

Hong-Yun Shang, Kai-Hua Jia, Nai-Wei Li, Min-Jie Zhou, Hao Yang, Xiao-Ling Tian, Yong-Peng Ma, Ren-Gang Zhang (2025). Phytop: a tool for visualizing and recognizing signals of incomplete lineage sorting and hybridization using species trees output from ASTRAL, Horticulture Research, Volume 12, Issue 3, uhae330, 10.1093/hr/uhae330

Huang, D.I., Hefer, C.A., Kolosova, N., Douglas, C.J. and Cronk, Q.C.B. (2014), Whole plastome sequencing reveals deep plastid divergence and cytonuclear discordance between closely related balsam poplars, Populus balsamifera and P. trichocarpa (Salicaceae). New Phytol, 204: 693–703. 10.1111/nph.12956

Hufford, M. B., Lubinksy, P., Pyhäjärvi, T., Devengenzo, M. T., Ellstrand, N. C., & Ross-Ibarra, J. (2013). The Genomic Signature of Crop-Wild Introgression in Maize. PLoS Genetics, 9(5), e1003477. 10.1371/journal.pgen.1003477

Stella Huynh, Thomas Marcussen, François Felber, Christian Parisod, Hybridization preceded radiation in diploid wheats, Molecular Phylogenetics and Evolution, Volume 139,2019,106554,ISSN 1055-7903,10.1016/j.ympev.2019.106554.

Inkscape Project. (2020). Inkscape. https://inkscape.org

Innes, P.A., Smart, B. C., Barham, J. A. M., Hulke, B. S., Kane, N. C. (2023). Chromosome-scale Genome Assembly of Lewis Flax (Linum lewisii Pursh.). biorXiv. 10.1101/2023.10.10.561607

Innes, P. A., Marcus, Z., Hulke, B. S., Kane, N. C. (2025). Glaciation history and geographic barriers shape population genetic structure and diversity of Lewis flax across western North America. bioRxiv. 10.1101/2025.08.18.670546

Jhala, A. J., Bhatt, H., Topinka, K., & Hall, L. M. (2011). Pollen-mediated gene flow in flax (*Linum usitatissimum L.*): Can genetically engineered and organic flax coexist? Heredity, 106(4), 557–566. 10.1038/hdy.2010.81

Jin, Z.-T., Lin, X.-H., Ma, D.-K., Xie, S.-Y., Huang, J.-X., Ren, C., Zhao, L., Duan, L., Xu, C., Hodel, R.G.J., Wu, J. and Liu, B.-B. (2025), Unravelling the Web of Life: Incomplete Lineage Sorting and Hybridisation as Primary Mechanisms Over Polyploidisation in the Evolutionary Dynamics of Pear Species. Mol Ecol Resour, 25: e70029. 10.1111/1755-0998.70029

Johnson, M.G., Gardner, E.M., Liu, Y., Medina, R., Goffinet, B., Shaw, A.J., Zerega, N.J.C. and Wickett, N.J. (2016), HybPiper: Extracting coding sequence and introns for phylogenetics from high-throughput sequencing reads using target enrichment. Applications in Plant Sciences, 4, 1600016. 10.3732/apps.1600016

Johnson, Pokorny, Dodsworth, Botigué, Cowan, Devault, Eiserhardt, Epitawalage, Forest, Kim, Leebens-Mack, Leitch, Maurin, D Soltis, P Soltis, Wong, Baker, Wickett, A Universal Probe Set for Targeted Sequencing of 353 Nuclear Genes from Any Flowering Plant Designed Using k-Medoids Clustering, Systematic Biology, Volume 68, Issue 4, July 2019, Pages 594–606,

Kadereit, J. W., & Abbott, R. J. (2021). Plant speciation in the Quaternary. Plant Ecology & Diversity, 14(3–4), 105–142. 10.1080/17550874.2021.2012849

Katoh, K., & Standley, D. M. (2013). MAFFT Multiple Sequence Alignment Software Version 7: Improvements in Performance and Usability. Molecular Biology and Evolution, 30(4), 772–780. 10.1093/molbev/mst010

Keller, S. R., Sowell, D. R., Neiman, M., Wolfe, L. M., & Taylor, D. R. (2009). Adaptation and colonization history affect the evolution of clines in two introduced species. New Phytologist, 183(3), 678–690. 10.1111/j.1469-8137.2009.02892.x

Kreitschitz, A., Kovalev, A., & Gorb, S. N. (2015). Slipping vs sticking: Water-dependent adhesive and frictional properties of Linum usitatissimum L. seed mucilaginous envelope and its biological significance. Acta Biomaterialia, 17, 152–159. 10.1016/j.actbio.2015.01.042

Korneliussen, T.S., Albrechtsen, A. & Nielsen, R. (2014). ANGSD: Analysis of Next Generation Sequencing Data. BMC Bioinformatics 15, 356. 10.1186/s12859-014-0356-4

Landoni, B., Suárez-Montes, P., Habeahan, R. H. F., Brennan, A. C., Pérez-Barrales, R. (2024). Local climate and vernalization sensitivity predict the latitudinal patterns of flowering onset in the crop wild relative Linum bienne Mill. Annals of Botany, 134 (1), 117–130. 10.1093/aob/mcae040

Langmead, B., & Salzberg, S. L. (2013). Fast gapped-read alignment with Bowtie 2. Nature Methods, 9(4), 357–359. 10.1038/nmeth.1923.Fast

Leroy, T., Louvet, J.-M., Lalanne, C., Le Provost, G., Labadie, K., Aury, J.-M., Delzon, S., Plomion, C. and Kremer, A. (2020), Adaptive introgression as a driver of local adaptation to climate in European white oaks. New Phytologist, 226: 1171-1182. 10.1111/nph.16095

Li, H., Handsaker, B., Wysoker, A., Fennell, T., Ruan, J., Homer, N., Marth, G., Abecasis, G., & Durbin, R. (2009). The Sequence Alignment/Map format and SAMtools. Bioinformatics, 25(16), 2078–2079. 10.1093/bioinformatics/btp352

Liston, A., Cronn, R., & Ashman, T. L. (2014). Fragaria: A genus with deep historical roots and ripe for evolutionary and ecological insights. American Journal of Botany, 101(10), 1686–1699. 10.3732/ajb.1400140

López-Jurado, J., Mateos-Naranjo, E. and Balao, F. (2019). Niche divergence and limits to expansion in the high polyploid Dianthus broteri complex. New Phytologist, 222: 1076-1087. 10.1111/nph.15663

Lovell, J. T., MacQueen, A. H., Mamidi, S., Bonnette, J., Jenkins, J., Napier, J. D., Sreedasyam, A., Healey, A., Session, A., Shu, S., Barry, K., Bonos, S., Boston, L. B., Daum, C., Deshpande, S., Ewing, A., Grabowski, P. P., Haque, T., Harrison, M.,… Schmutz, J. (2021). Genomic mechanisms of climate adaptation in polyploid bioenergy switchgrass. Nature, 590(7846), 438–444. 10.1038/s41586-020-03127-1

MacLay, T. G., Birch, J. L., Gunn, B. F., Ning, W., Tate, J. A., Nauheimer, L.,… & Jackson, C. J. (2021). New targets acquired: Improving locus recovery from the Angiosperms353 probe set. Applications in plant sciences, 9(7).

Magri, D. (2008). Patterns of post-glacial spread and the extent of glacial refugia of European beech (Fagus sylvatica). Journal of Biogeography, 35(3), 450–463. 10.1111/j.1365-2699.2007.01803.x

Magri, D., Vendramin, G. G., Comps, B., Dupanloup, I., Geburek, T., Gömöry, D., Latałowa, M., Litt, T., Paule, L., Roure, J. M., Tantau, I., Van Der Knaap, W. O., Petit, R. J., & De Beaulieu, J. L. (2006). A new scenario for the Quaternary history of European beech populations: Palaeobotanical evidence and genetic consequences. New Phytologist, 171(1), 199–221. 10.1111/j.1469-8137.2006.01740.x

Maguilla, E., Escudero, M., Ruíz-Martín, J., Arroyo, J., & Schneeweiss, G. (2021). Origin and diversification of flax and their relationship with heterostyly across the range. Journal of Biogeography, 48(8), 1994–2007. 10.1111/jbi.14129

Marcet-Houben, M., & Gabaldón, T. (2015). Beyond the whole-genome duplication: Phylogenetic evidence for an ancient interspecies hybridization in the baker’s yeast lineage. PLoS Biology, 13(8), 1–26. 10.1371/journal.pbio.1002220

Marshall, D. L., Avritt, J. J., Maliakal-Witt, S., Medeiros, J. S., Shaner, M. G. M. (2010). The impact of plant and flower age on mating patterns. Annals of Botany, 105(1), 7–22. 10.1093/aob/mcp260

McDill, J., Repplinger, M., Simpson, B. B., & Kadereit, J. W. . (2009). The Phylogeny of Linum and Linaceae Subfamily Linoideae, with Implications for Their Systematics, Biogeography, and Evolution of Heterostyly. 34(2), 386–405.

Médail, F. and Diadema, K. (2009). Glacial refugia influence plant diversity patterns in the Mediterranean Basin. Journal of Biogeography, 36: 1333-1345. 10.1111/j.1365-2699.2008.02051.x

Meisner J, Albrechtsen A. (2018). Inferring Population Structure and Admixture Proportions in Low-Depth NGS Data. Genetics, 210(2), 719-731. doi: 10.1534/genetics.118.301336

Middleton CP, Senerchia N, Stein N, Akhunov ED, Keller B, et al. (2014) Sequencing of Chloroplast Genomes from Wheat, Barley, Rye and Their Relatives Provides a Detailed Insight into the Evolution of the Triticeae Tribe. PLOS ONE 9(3): e85761. 10.1371/journal.pone.0085761

Miller, P. (1768). The Gardeners Dictionary. 8th edition, 372–374. 10.5962/bhl.title.541

Migliore, J., Baumel, A., Juin, M., & Médail, F. (2012). From Mediterranean shores to central Saharan mountains: Key phylogeographical insights from the genus Myrtus. Journal of Biogeography, 39(5), 942–956. 10.1111/j.1365-2699.2011.02646.x

Migliore, J., Baumel, A., Leriche, A., Juin, M., & Médail, F. (2018). Surviving glaciations in the Mediterranean region: An alternative to the long-term refugia hypothesis. Botanical Journal of the Linnean Society, 187(4), 537–549. 10.1093/botlinnean/boy032

Morales-Briones, D. F., Liston, A., & Tank, D. C. (2018). Phylogenomic analyses reveal a deep history of hybridization and polyploidy in the Neotropical genus Lachemilla (Rosaceae). New Phytologist, 218(4), 1668–1684. 10.1111/nph.15099

Mousavi-Derazmahalleh, M., Bayer, P.E., Nevado, B. et al. (2018). Exploring the genetic and adaptive diversity of a pan-Mediterranean crop wild relative: narrow-leafed lupin. Theoretical and Applied Genetics, 131, 887–901. 10.1007/s00122-017-3045-7

Murillo-A., J., J. Valencia-D., C. I. Orozco, C. Parra-O., and K.M. Neubig. 2022. Incomplete lineage sorting and reticulate evolution mask species relationships in Brunelliaceae, an Andean family with rapid, recent diversification. American Journal of Botany109(7): 1139–1156. 10.1002/ajb2.16025

Myers, N., Mittermeier, R., Mittermeier, C. et al. (2000). Biodiversity hotspots for conservation priorities. Nature, 403, 853–858. 10.1038/35002501

Nguyen, L.-T., Schmidt, H. A., von Haeseler, A., & Minh, B. Q. (2015). IQ-TREE: A Fast and Effective Stochastic Algorithm for Estimating Maximum-Likelihood Phylogenies. Molecular Biology and Evolution, 32(1), 268–274. 10.1093/molbev/msu300

Nieto Feliner, G. (2014). Patterns and processes in plant phylogeography in the Mediterranean Basin. A review. Perspectives in Plant Ecology, Evolution and Systematics, 16(5), 265–278. 10.1016/j.ppees.2014.07.002

Nieto Feliner, G., Cellinese, N., Crowl, A. A., Frajman B. (2023). Editorial: Understanding plant diversity and evolution in the Mediterranean Basin. Frontiers in Plant Science. 10.3389/fpls.2023.1152340

Ockendon, D.J. (1968), BIOSYSTEMATIC STUDIES IN THE LINUM PERENNE GROUP. New Phytologist, 67: 787–813. 10.1111/j.1469-8137.1968.tb06396.x

Paradis, E., & Schliep, K. (2019). ape 5.0: An environment for modern phylogenetics and evolutionary analyses in R. Bioinformatics, 35(3), 526–528. 10.1093/bioinformatics/bty633

Perret, j., Cobelli, O., Taudière, A., Andrieu, J., Aumeeruddy-Thomas, Y., Souissi, J. B., Besnard, G., Casazza, G., Crochet, P-A., Decaëns, T., Denis, F., Geniez, P., Loizides, M., Médail, F., Pasqualini, V., Speciale, C., Battesti, V., Chevaldonné, P., Lejeusne, C., Richard, F. (2023). Time to refine the geography of biodiversity hotspots by integrating molecular data: The Mediterranean Basin as a case study. Biological Conservation, 284. 10.1016/j.biocon.2023.110162.

Pokorny L, Pellicer J, Woudstra Y, Christenhusz MJM, Garnatje T, Palazzesi L, Johnson MG, Maurin O, Françoso E, Roy S, Leitch IJ, Forest F, Baker WJ and Hidalgo O. (2024). Genomic incongruence accompanies the evolution of flower symmetry in Eudicots: a case study in the poppy family (Papaveraceae, Ranunculales). Front. Plant Sci. 15:1340056. doi: 10.3389/fpls.2024.1340056

Evolutionary consequences of repeated loss of distyly in Linum. Zoé Postel, Panagiotis-Ioannis Zervakis, Marco Fracassetti, Aleksandra Losvik, Matias Wanntorp, Lucile Soler, Allison Churcher, Aelys M. Humphreys, Tanja Slotte. bioRxiv 2026.03.03.709227; doi: 10.64898/2026.03.03.709227

Purugganan, M. D. (2019). Evolutionary Insights into the Nature of Plant Domestication. Current Biology, 29(14), R705–R714. 10.1016/j.cub.2019.05.053

Razanamaro, O. H. M., Randriatsitohaina, R. D., Jean Michel, L. P. T., Ramiliarisona, L. F., Nantenaina, R. H., Raoelinjanakolona, N. N.,… & Viruel, J. (2025). Integrating genomics and habitat surveys to uncover population structure and regeneration challenges in Adansonia suarezensis (Malvaceae). Annals of Botany, mcaf320.

Rasmussen, M. S., Garcia-Erill, G., Korneliussen, T. S., Wiuf, C., & Albrechtsen, A. (2022). Estimation of site frequency spectra from low-coverage sequencing data using stochastic EM reduces overfitting, runtime, and memory usage. Genetics, 222(4), iyac148.

Ren R, Wang H, Guo C et al. (2018). Widespread Whole Genome Duplications Contribute to Genome Complexity and Species Diversity in Angiosperms. Molecular Plant, 2018; 11, 414–428

Revell, L. J. (2012). phytools: An R package for phylogenetic comparative biology (and other things): *phytools: R package*. Methods in Ecology and Evolution, 3(2), 217–223. 10.1111/j.2041-210X.2011.00169.x

Ruiz-Martín, J., Santos-Gally, R., Escudero, M., Midgley, J. J., Pérez-Barrales, R., & Arroyo, J. (2018). Style polymorphism in Linum (Linaceae): A case of Mediterranean parallel evolution? Plant Biology, 20(4), 100–111. 10.1111/plb.12670

Sancho, R., Cantalapiedra, C. P., López-Alvarez, D., Gordon, S. P., Vogel, J. P., Catalán, P., & Contreras-Moreira, B. (2018). Comparative plastome genomics and phylogenomics of Brachypodium: Flowering time signatures, introgression and recombination in recently diverged ecotypes. New Phytologist, 218(4), 1631–1644. 10.1111/nph.14926

Santos-Gally, R., Vargas, P. and Arroyo, J. (2012). Insights into Neogene Mediterranean biogeography based on phylogenetic relationships of mountain and lowland lineages of Narcissus (Amaryllidaceae). Journal of Biogeography, 39: 782-798. 10.1111/j.1365-2699.2011.02526.x

Scheunert, A., & Heubl, G. (2014). Diversification of Scrophularia (Scrophulariaceae) in the Western Mediterranean and Macaronesia—Phylogenetic relationships, reticulate evolution and biogeographic patterns. Molecular Phylogenetics and Evolution, 70(1), 296–313. 10.1016/j.ympev.2013.09.023

Schiffels, S., & Wang, K. (2020). MSMC and MSMC2: the multiple sequentially Markovian coalescent. In Statistical population genomics (pp. 147–165). Humana.

Skotte L, Korneliussen TS, Albrechtsen A. Estimating individual admixture proportions from next generation sequencing data. Genetics. 2013 Nov;195(3):693–702. doi: 10.1534/genetics.113.154138

Smith, S. A., & O’Meara, B. C. (2012). treePL: Divergence time estimation using penalized likelihood for large phylogenies. Bioinformatics, 28(20), 2689–2690. 10.1093/bioinformatics/bts492

Smýkal, P., Hradilová, I., Trněný, O. et al. (2017). Genomic diversity and macroecology of the crop wild relatives of domesticated pea. Scientific Reports, 7. 10.1038/s41598-017-17623-4

Solís-Lemus C, Ané C (2016) Inferring Phylogenetic Networks with Maximum Pseudolikelihood under Incomplete Lineage Sorting. PLOS Genetics 12(3): e1005896. 10.1371/journal.pgen.1005896

Soto-Cerda, B. J., Diederichsen, A., Duguid, S., Booker, H., Rowland, G., & Cloutier, S. (2014). The potential of pale flax as a source of useful genetic variation for cultivated flax revealed through molecular diversity and association analyses. Molecular Breeding, 34(4), 2091–2107. 10.1007/s11032-014-0165-5

Stull, G.W., Pham, K.K., Soltis, P.S. and Soltis, D.E. (2023), Deep reticulation: the long legacy of hybridization in vascular plant evolution. The Plant Journal, 114: 743-766. 10.1111/tpj.16142

Suc, J-P., Popescu, S-M., Fauquette, S., Bessedik, M., Jiménez-Moreno, G., Bachiri Taoufiq, N., Zheng, Z., Médail, F., Klotz, S. (2018). Reconstruction of Mediterranean flora, vegetation and climate for the last 23 million years based on an extensive pollen dataset. Ecologia Mediterranea, 44. 10.3406/ecmed.2018.2044.

Shan-Shan Sun, Shu-Han Xu, Wen-Jie Yi, Shi-Long Chen, Pan Li, Daiki Takahashi, Harue Abe, Xiao-Lei Ma, Peng-Cheng Fu, Incomplete lineage sorting, hybridization and polyploidization blurred phylogenetic relationships of gentians from the Qinghai-Tibetan Plateau, Molecular Phylogenetics and Evolution, Volume 214, 2026, 108476, ISSN 1055-7903, 10.1016/j.ympev.2025.108476.

Sveinsson, S., McDill, J., Wong, G. K. S., Li, J., Li, X., Deyholos, M. K., & Cronk, Q. C. B. (2014). Phylogenetic pinpointing of a paleopolyploidy event within the flax genus (Linum) using transcriptomics. Annals of Botany, 113(5), 753–761. 10.1093/aob/mct306

Tank, D.C., Eastman, J.M., Pennell, M.W., Soltis, P.S., Soltis, D.E., Hinchliff, C.E., Brown, J.W., Sessa, E.B. and Harmon, L.J. (2015), Nested radiations and the pulse of angiosperm diversification: increased diversification rates often follow whole genome duplications. New Phytol, 207: 454–467. 10.1111/nph.13491

Than, C., Ruths, D. & Nakhleh, L. (2008), PhyloNet: a software package for analyzing and reconstructing reticulate evolutionary relationships. BMC Bioinformatics 9, 322. 10.1186/1471-2105-9-322

Tillich, M., Lehwark, P., Pellizzer, T., Ulbricht-Jones, E. S., Fischer, A., Bock, R., & Greiner, S. (2017). GeSeq – versatile and accurate annotation of organelle genomes. Nucleic Acids Research, 45(W1), W6–W11. 10.1093/nar/gkx391

Toledo, B., Marcer, A., Méndez-Vigo, B., Alonso-Blanco, C., & Picó, F. X. (2020). An ecological history of the relict genetic lineage of Arabidopsis thaliana. Environmental and Experimental Botany, 170(June 2019), 103800. 10.1016/j.envexpbot.2019.103800

Uysal, H., Fu, Y. B., Kurt, O., Peterson, G. W., Diederichsen, A., & Kusters, P. (2010). Genetic diversity of cultivated flax (Linum usitatissimum L.) and its wild progenitor pale flax (Linum bienne Mill.) as revealed by ISSR markers. Genetic Resources and Crop Evolution, 57(7), 1109–1119. 10.1007/s10722-010-9551-y

Uysal, H., Kurt, O., Fu, YB. et al. (2012). Variation in phenotypic characters of pale flax (Linum bienne Mill.) from Turkey. Genetic Resources and Crop Evolution, 59, 19–30. 10.1007/s10722-011-9663-z

Valdés-Florido, A., Tan, L., Maguilla, E., Simón-Porcar, V. I., Zhou, Y-H., Arroyo, J., Escudero, M. (2023). Drivers of diversification in Linum (Linaceae) by means of chromosome evolution: correlations with biogeography, breeding system and habit. Annals of Botany, 132 (5), 949–962. 10.1093/aob/mcad139

Valdés-Florido, A., González-Toral, C., Maguilla, E., Cires, E., Díaz-Lifante, Z., Andrés-Camacho, C., et al. (2024a). Polyploidy and hybridization in the Mediterranean: unravelling the evolutionary history of Centaurium (Gentianaceae). Annals of Botany, 134, 247–262. doi: 10.1093/aob/mcae066

Valdés-Florido A., Valcárcel V., Maguilla E., Díaz-Lifante Z., Andrés-Camacho C., Zeltner L., Coca-de-la-Iglesia M., Medina N. G., Arroyo J., Escudero M. (2024). The interplay between climatic niche evolution, polyploidy and reproductive traits explains plant speciation in the Mediterranean Basin: a case study in Centaurium (Gentianaceae). Frontiers in Plant Sciences. 15:1439985. doi: 10.3389/fpls.2024.1439985

Vanrell, M. A., Novaes, L. R., Afonso, A., Arroyo, J., Simón-Porcar, V. (2024b). Ecological correlates of population genetics in Linum suffruticosum, an heterostylous polyploid and taxonomic complex endemic to the Western Mediterranean Basin. Annals of Botany PLANTS, 16(4). 10.1093/aobpla/plae027

Vieira, F. G., Lassalle, F., Korneliussen, T. S., & Fumagalli, M. (2016). Improving the estimation of genetic distances from Next-Generation Sequencing data: Genetic Distances from NGS Data. Biological Journal of the Linnean Society, 117(1), 139–149. 10.1111/bij.12511

Villalvazo-Hernández, A., Burgos-Hernández, M., & González, D. (2022). Phylogenetic Analysis and Flower Color Evolution of the Subfamily Linoideae (Linaceae). Plants, 11(12), 1579. 10.3390/plants11121579

Vincent, H., Amri, A., Castaneda-Alvarez, N. P., Dempewolf, A., Dulloo, E., Guarino, L., Hole, D., Mba, C., Toledo, A., & Maxted, N. (2019). Modeling of crop wild relative species identifies areas globally for in situ conservation. Communications Biology, 2(1). 10.1038/s42003-019-0372-z

Viruel J, Conejero M, Hidalgo O, Pokorny L, Powell RF, Forest F, Kantar MB, Soto Gomez M, Graham SW, Gravendeel B, Wilkin P and Leitch IJ (2019) A Target Capture-Based Method to Estimate Ploidy From Herbarium Specimens. Front. Plant Sci. 10:937. doi: 10.3389/fpls.2019.00937

Viruel, J., Kantar, M. B., Gargiulo, R., Hesketh-Prichard, P., Leong, N., Cockel, C., Forest, F., Gravendeel, B., Pérez-Barrales, R., Leitch, I. J., & Wilkin, P. (2020a). Crop wild phylorelatives (CWPs): Phylogenetic distance, cytogenetic compatibility and breeding system data enable estimation of crop wild relative gene pool classification. Botanical Journal of the Linnean Society, Dd, 1–33. 10.1093/botlinnean/boaa064

Viruel, J., Le Galliot, N., Pironon, S., Nieto Feliner, G., Suc, J., Lakhal-Mirleau, F., Juin, M., Selva, M., Bou Dagher Kharrat, M., Ouahmane, L., La Malfa, S., Diadema, K., Sanguin, H., Médail, F., & Baumel, A. (2020b). A strong east–west Mediterranean divergence supports a new phylogeographic history of the carob tree ( *Ceratonia siliqua*, Leguminosae) and multiple domestications from native populations. Journal of Biogeography, 47(2), 460–471. 10.1111/jbi.13726

Xu, L.-L., Yu, R.-M., Lin, X.-R., Zhang, B.-W., Li, N., Lin, K., Zhang, D.-Y. and Bai, W.-N. (2021), Different rates of pollen and seed gene flow cause branch-length and geographic cytonuclear discordance within Asian butternuts. New Phytol, 232: 388–403. 10.1111/nph.17564

Xu Y, Wei Y, Zhou Z, et al. (2023). Widespread incomplete lineage sorting and introgression shaped adaptive radiation in the Gossypium genus. Plant Communications, 2023; 5

Wagner, F., Ott, T., Zimmer, C., Reichhart, V., Vogt, R. and Oberprieler, C. (2019), ‘At the crossroads towards polyploidy’: genomic divergence and extent of homoploid hybridization are drivers for the formation of the ox-eye daisy polyploid complex (Leucanthemum, Compositae-Anthemideae). New Phytolologist, 223: 2039-2053. 10.1111/nph.15784

Wahlberg, N., Weingartner, E., Warren, A.D. et al. Timing major conflict between mitochondrial and nuclear genes in species relationships of Polygoniabutterflies (Nymphalidae: Nymphalini). BMC Evol Biol 9, 92 (2009). 10.1186/1471-2148-9-92

Wang, Z., Hobson, N., Galindo, L., Zhu, S., Shi, D., McDill, J., Yang, L., Hawkins, S., Neutelings, G., Datla, R., Lambert, G., Galbraith, D. W., Grassa, C. J., Geraldes, A., Cronk, Q. C., Cullis, C., Dash, P. K., Kumar, P. A., Cloutier, S.,… Deyholos, M. K. (2012). The genome of flax (Linum usitatissimum) assembled de novo from short shotgun sequence reads. Plant Journal, 72(3), 461–473. 10.1111/j.1365-313X.2012.05093.x

Wang, K., Mathieson, I., O’Connell, J., & Schiffels, S. (2020). Tracking human population structure through time from whole genome sequences. PLoS genetics, 16(3), e1008552.

Dan L. Warren, Anthony J. Geneva, Robert Lanfear, RWTY (R We There Yet): An R Package for Examining Convergence of Bayesian Phylogenetic Analyses, Molecular Biology and Evolution, Volume 34, Issue 4, April 2017, Pages 1016–1020, 10.1093/molbev/msw279

Weiss, E., & Zohary, D. (2011). The Neolithic Southwest Asian Founder Crops. Current Anthropology, 52(S4), S237–S254. 10.1086/658367

Wickham, H. (2016). ggplot2: Elegant Graphics for Data Analysis. Springer-Verlag. https://ggplot2.tidyverse.org

Wickham, H., & Girlich, M. (2022). tidyr: Tidy Messy Data (1.3.1). https://tidyr.tidyverse.org, https://github.com/tidyverse/tidyr

Xi, Z., Ruhfel, B. R., Schaefer, H., Amorim, A. M., Sugumaran, M., Wurdack, K. J., Endress, P. K., Matthews, M. L., Stevens, P. F., Mathews, S., & Davis, C. C. (2012). Phylogenomics and a posteriori data partitioning resolve the Cretaceous angiosperm radiation Malpighiales. Proceedings of the National Academy of Sciences of the United States of America, 109(43), 17519–17524. 10.1073/pnas.1205818109

Zhang, C., Rabiee, M., Sayyari, E., & Mirarab, S. (2018). ASTRAL-III: Polynomial time species tree reconstruction from partially resolved gene trees. BMC Bioinformatics, 19(S6), 153. 10.1186/s12859-018-2129-y

Zhao, Y., Yu, D., Kuo, W., Huang, J., Guo, J., Sun, M., Hu, Y., Soltis, D. E., Soltis, P. S., Ma, H., et al. (2025). Nuclear phylogenomics provide evidence to clarify key morphological evolution and whole-genome duplication across rosids. J. Integr. Plant Biol.67: 2704–2730.

Zizka A, Silvestro D, Andermann T, Azevedo J, Duarte Ritter C, Edler D, Farooq H, Herdean A, Ariza M, Scharn R, Svanteson S, Wengstrom N, Zizka V, Antonelli A (2019). CoordinateCleaner: standardized cleaning of occurrence records from biological collection databases. Methods in Ecology and Evolution. doi:10.1111/2041-210X.13152

Zunino L, Cubry P, Sarah G, Mournet P, El Bakkali A, Aqbouch L, Sidibé-Bocs S, Costes E, Khadari B.(2024). Genomic evidence of genuine wild versus admixed olive populations evolving in the same natural environments in western Mediterranean Basin. PLoS One, 19(1). doi: 10.1371/journal.pone.0295043

Zuntini, A.R., Carruthers, T., Maurin, O. et al. (2024). Phylogenomics and the rise of the angiosperms. Nature 629, 843–850. 10.1038/s41586-024-07324-0

