## Supplementary Information for "Reticulate evolution and climate driven diversification shaped the origin and geographic structure of *Linum bienne*"

**New Phytologist Supporting Information**

**TITLE:**

Yann Bourgeois^2,6^ (0000-0002-1809-387X)

Robin G. Allaby^7^ (0000-0001-5046-002X)

Adrian C. Brennan^8^ (ORCID: 0000-0002-8171-769X)

^1^ Department of Biosciences, University of Milan. 20133 Milano, IT.

^2^ School of Biological Sciences, University of Portsmouth. PO12DY Portsmouth, UK

^3^ Technological College, University of Zaragoza. 22071 Zaragoza, ES.

^4^ Royal Botanic Gardens, Kew. TW9 3AB Richmond, UK.

^5^ Institute for Biocomputation and Physics of Complex Systems, University of Zaragoza. 50018 Zaragoza, ES

^6^ UMR DIADE, University of Montpellier. CIRAD, IRD, 34394 Montpellier Cedex 5, FR

^7^ Faculty of Science, Engineering and Medicine, School of Life Sciences, Gibbet Hill Campus, University of Warwick. CV4 7AL Coventry, UK.

^8^ Biosciences Department, Durham University. DH1 3LE Durham, UK

^9^ Botany Department, University of Granada. 18071 Granada, ES.

* corresponding authors

The following Supporting Information is available for this article:

**Figure S1.** Bootstrap-tree conflict in plastid and nuclear genus-level phylogenies of *Linum*.

**Figure S2.** Incomplete lineage sorting and introgression across genus-level *Linum* sampling.

**Figure S3.** Divergence-time estimates for genus-level plastid and nuclear phylogenies of *Linum*.

**Figure S4.** Plastome and nuclear genomic relationships among *Linum bienne* and *L. usitatissimum* accessions.

**Figure S5.** Plastid-gene phylogeny including *Linum villarianum*.

**Figure S6.** Nuclear population structure of *Linum bienne*.

**Figure S7.** Plastome-based BEAST dating of *Linum bienne* lineages.

**Figure S8.** BEAST parameter distributions and convergence diagnostics.

**Figure S9.** Demographic history and migration inferred with MSMC-IM based on nuclear genomes.

**Figure S10.** Present-day environmental niche model performance for *Linum bienne*.

**Figure S11.** Past and present climatic favourability predictions for *Linum bienne*.

**Figure S12.** Accuracy of genotype imputation with STITCH.

**Table S1.** Taxon sampling and gene recovery for genus-level phylogenomic analyses in *Linum*.

**Table S2.** HybPiper recovery statistics for nuclear Angiosperms353 loci and plastid genes.

**Table S3.** Range-wide sampling of *Linum bienne* and *L. usitatissimum* individuals used for plastome and nuclear genomic analyses.

**Table S4.** QuIBL test results for ILS and introgression based on triplets of *Linum* individuals.

**Table S5.** Individual plastid-gene support for the two alternative topologies at the *L. bienne* node retrieved from the literature.
**Table S6.** Comparison of published phylogenetic topologies at selected nodes within *Linum*.

**Methods S1.** Hybpiper pipeline details for Angiosperms353 nuclear loci and plastid loci

**Methods S2.** Assessing the nature of conflicting phylogenetic signal across *Linum* and for the *Linum bienne* node using Angiosperms353 and plastid genes
**Methods S3.** Nuclear genome analyses in *Linum bienne*

**Methods S4.** Environmental niche of *Linum bienne* from the Pliocene to present

*Note that Supplementary Figures are provided in a separate powerpoint file, Supplementary Tables are provided in a separate excel file, and Supplementary Methods in a separate word file.*

**Extended captions**

**Figure S1.** Bootstrap-tree conflict in plastid and nuclear genus-level phylogenies of *Linum*. DensiTree plots summarizing 1000 unrooted bootstrap trees for the (A) plastid and (B) nuclear Angiosperms353 supermatrix datasets. Branches in black follow the main consensus topology, whereas branches in red represent alternative topologies recovered among bootstrap replicates. The plastid dataset showed conflict around the *L. bienne* node, both among species and among *L. bienne* and *L. usitatissimum* accessions. The nuclear dataset showed conflict mainly among *L. bienne* and *L. usitatissimum* accessions, and around the *L. flavum* node. Supplementary Table 1 includes information about all samples.

**Figure S2.** Incomplete lineage sorting and introgression across genus-level *Linum* sampling. (A) Phytop quartet-topology analysis using the ASTRAL nuclear tree as the guide tree. Bars show the relative frequency of the three most common quartet topologies across nuclear gene trees for each node. Similar frequencies of the three topologies are consistent with incomplete lineage sorting, whereas prevalence on the main topology and a single alternative topology may indicate hybridization or introgression. (B) QuIBL analysis testing whether gene-tree discordance is better explained by incomplete lineage sorting alone (null model) or by incomplete lineage sorting plus introgression (alternative model), using branch-length information from nuclear gene trees. Average introgression fractions are shown by tile colour, with darker colours meaning higher introgression fraction. Statistically supported ILS+introgression models were recovered only for triplets involving the *L. bienne* node. Supplementary Table 1 includes information about all samples.

**Figure S3.** Divergence-time estimates for genus-level plastid and nuclear phylogenies of *Linum*. Penalized-likelihood dated phylogenies inferred from (A) the plastid supermatrix, (B) the nuclear Angiosperms353 supermatrix, and (C) the nuclear ASTRAL tree. The same fossil calibration was applied to the crown node of *Linum* in all analyses. Major nodes within *Linum* were generally placed within similar broad time windows, but age estimates differed for the *L. bienne* crown node: the ASTRAL nuclear tree recovered the youngest estimate, the nuclear supermatrix recovered the oldest estimate, and the plastid tree recovered an intermediate estimate. Boxes in (A) indicate comparable age estimates from previous studies for nodes relevant to the evolutionary history of *L. bienne*. Supplementary Table 1 includes information about all samples.

**Figure S4.** Plastome and nuclear genomic relationships among *Linum bienne* and *L. usitatissimum* accessions. (A) Plastome phylogeny inferred with maximum likelihood in IQ-TREE (GTR+G). (B) Nuclear genetic-distance tree inferred with ngsDist from low-depth nuclear genome data. *Linum narbonense* was used as outgroup in both analyses (outgroup was pruned for representation purposes). Bootstrap (1000) is represented by light blue circles. Light grey lines connect the same individuals between trees; darker lines indicate *L. usitatissimum* individuals. Colours indicate the geographically structured lineages recovered in the plastome and nuclear datasets: SW (red), Central (yellow), SE (dark blue) and NW (light blue), with the SW+central sister to the SE+NW, and NW embedded within SE. Most individuals were assigned to the same lineage in both datasets, but several showed cytonuclear discordance, with plastid and nuclear data assigning the same individual to different lineages. Supplementary Table 3 includes information about all samples.

**Figure S5.** Plastid-gene phylogeny including *Linum villarianum*. IQtree maximum-likelihood phylogeny (GTR+G) inferred from a reduced plastid-gene alignment including *L. bienne* individuals and one *L. villarianum* accession retrieved from published sequence data. The analysis was used to assess whether *L. villarianum* falls within any of the major *L. bienne* lineages or is recovered as sister to *L. bienne*. Supplementary Table 3 includes information about all *L. bienne* samples.

**Figure S6.** Nuclear population structure of *Linum bienne*. (A) ngsADMIX ancestry coefficients inferred from low-depth nuclear genome data across values of K. (B) Principal component analysis based on nuclear genotype likelihoods. Colours correspond to the major *L. bienne* lineages recovered from nuclear and plastid analyses. *Linum usitatissimum* accessions formed a separate genetic cluster, shown in grey. Supplementary Table 3 includes information about all samples.

**Figure S7.** Plastome-based BEAST dating of *Linum bienne* lineages. Maximum clade credibility trees inferred from the *L. bienne* and *L. usitatissimum* plastome alignment, using *L. narbonense* as outgroup. Three BEAST analyses were run using alternative secondary calibration points derived from genus-level treePL analyses based on (A) plastid, (B) nuclear supermatrix, and (C) nuclear ASTRAL phylogenies. Horizontal bars indicate uncertainty around node-age estimates. Plastid and nuclear-supermatrix calibrations produced older age estimates than the nuclear-ASTRAL calibration. Overall, it was estimated that *L. bienne* started to diverge from other *Linum* species between the early to mid-Pliocene. Horizontal bars represent posterior probabilities for the age of each node and they are coloured with: black (*L. bienne* crown node), red (southwestern lineage), yellow (central lineage), blue (southeastern + northwestern lineages), grey (any other node). Supplementary Table 3 includes information about all samples.

**Figure S8.** BEAST parameter distributions and convergence diagnostics. BEAST parameter distribution from 1,000,000,000runs (10% burn-in in black) for the dating of the *L. bienne* plastome phylogeny using differing secondary calibration points for the *L. bienne* crown node and the overall root using *L. narbonense* as outgroup. Parameters are shown for BEAST run with secondary calibration points from the *Linum* phylogenetic trees dated with treePL based on: plastid (A), nuclear supermatrix (B), and nuclear-Astral (C) datasets. All three types of calibration resulted in high ESS (>200) for all parameters, except UCLD mean and standard deviation for the plastid calibration which had ESS < 200.

**Figure S9.** Demographic history and migration inferred with MSMC-IM based on nuclear genomes. Genetic clusters are indicated with colours and correspond to columns. Time (ca. 1Mya to present) is on the x-axis. The first row of panels represents effective population size for all combinations of clusters with lineages recovering from a large drop in population size around 10000 years ago (Holocene). The second row of panels represent cumulative migration probability, which is clearly higher between the NW and SE genetic clusters, but in general decreases from past to present highlighting when lineages started to diverge. The third row of panels represents migration rates. Migration rates are again quite high again for the NW and SE genetic clusters even in recent times, however a smaller spike was also found around the mid-Holocene between the SW and NW lineages, and possibly the same pattern is present between the central and NW lineages.

**Figure S10.** Present-day environmental niche model performance for *Linum bienne*. Present-day climatic suitability or favourability predicted using (A) Maxent, (B) GLM and (D) GAM models fitted with *L. bienne* occurrence data extracted from GBIF and WorldClim bioclimatic variables. (C) MESS was used to show climatic dissimilarity between the model training region (indicated by a square and including the Iberian Peninsula and France, excluding islands) and the broader projection area. In the analysis, low values (below zero – blue) indicate the highest dissimilarity in climate between the training area and the predicted area, while high values (above zero – yellow) indicate high climate similarity. Model performance assessed using AUC (> 0.7), MCS (0.5 < MCS < 1.5), and TSS (> 0.4). GAM and Maxent showed broadly similar performance, with GAM selected for past-climate projections, while GLM performed the worst. AUC = Area Under the Curve; MCS = Miller Calibration Score; TSS = True Skill Statistic.

**Figure S11.** Past and present climatic favourability for Linum bienne. GAM-based projections of climatic favourability for *L. bienne* (using GBIF occurrence <https://doi.org/10.15468/dl.nhbuhr> accessed on 03/07/2025, and WorldClim and pastclim bioclimatic variables) from the present to the early Pliocene. Panels show: present (A), Mid-Holocene – 8000 years ago (B), Last Glacial Maximum – 21000 years ago (C), transition between Last Interglacial and Last Glacial Maximum – 70000 years ago (D), Last Interglacial – 130000 years ago, Mid Pleistocene – 700000 years ago (F), Early Pleistocene – 1300000 years ago (G), Late Pliocene – 2700000 years ago (H), Early Pliocene – 4500000 years ago (I). Darker colours indicate higher predicted favourability, whereas lighter colours indicate lower predicted favourability. Across periods of reduced favourability, northern and higher-latitude regions were generally less favourable than southern Mediterranean regions.

**Figure S12.** Accuracy of genotype imputation with STITCH. Confusion matrix comparing observed high-depth genotypes with genotypes imputed from low-depth data using STITCH. The x-axis shows observed genotypes and the y-axis shows imputed genotypes. Warmer colours indicate more genotype calls. Higher counts along the diagonal indicate that imputed genotypes generally matched observed genotypes.

**Table S1.** Taxon sampling and gene recovery for genus-level phylogenomic analyses in *Linum*. All main sections in *Linum* (see Ruiz-Martin et al., 2018) were sampled except for section Cathartolinum. For each individual included, the table reports the main clade to which the species belong, species identity, chromosome number (retrieved from Bolsheva et al. (2017), Ruiz-Martin et al. (2018), and the Chromosome Count Database http://ccdb.tau.ac.il/), sample code, and data source (downloaded from Tree of Life or newly sequence in this study), country of origin and locality of collection (available for the newly sequence samples). All samples collected by the authors and relative reads were uploaded to the European Nucleotide Archive (ENA) under the project PRJEB86756. Under the column "ENA ID", ENA sample IDs are reported for samples collected by the authors, while under the column "Sample ID", IDs are reported for Tree of Life samples, together with sample aliases for samples collected by the authors. For both plastid and nuclear genes, samples were used for phylogenetic inference if they passed thresholds based on hybpiper statistics. The number of putative duplicate genes based on hybpiper statistics is also reported here.

**Table S2.** HybPiper recovery statistics for nuclear Angiosperms353 loci and plastid genes. For each accession, the table reports HybPiper recovery statistics for nuclear Angiosperms353 loci and plastid genes, including the number of reads mapped, percentage on target, number of genes recovered at different sequence-length thresholds, or paralogy warnings, stitched-contig information, chimera warnings, and total bases recovered. These statistics were used to assess sample and locus recovery before filtering for downstream phylogenetic analyses. Sample information is available in Table S1.

**Table S3.** Range-wide sampling of *Linum bienne* and *L. usitatissimum* individuals used for plastome and nuclear genomic analyses. For each individual, the table reports sample source (individual collected by Juan Viruel JV, Rocio Perez-Barrales RPB, Adrian C. Brennan ACB, Beatrice Landoni BL, Robin Allaby RA, or collected from herbaria sheet from Kew Garden), species identity, population or family code, sample ID, ENA accession when available, country, locality, geographic coordinates, altitude, plastid lineage assignment, nuclear lineage assignment, and sequencing depth. Samples were obtained from field collections by the authors or from herbarium material. When precise locality information was unavailable, city or country centroids were used to approximate geographic coordinates. Lineage assignments correspond to the plastome and nuclear analyses shown in Figure 3 and in Figure S4. All samples and relative reads were uploaded to the European Nucleotide Archive (ENA) under project PRJEB86756 and the relative sample IDs are reported. Note that 7 samples marked with (*) had previously been uploaded to NCBI under project PRJNA580472 and the NCBI codes are reported instead. Data for one sample were not published because sequencing yielded bad quality reads, while one sample was not published because the species could not be determined (not *L. bienne*).

**Table S4.** QuIBL test results for ILS and introgression based on triplets of Linum individuals. The table was formatted using the R package QuiblR. Column isDiscordant indicates whether the topology retrieved for the triplet is the same (0) or not (1) as the species tree provided to QuIBL (Astral tree). The column isSignificant indicates whether an ILS-only (0) or ILS+introgression model (1) better explained the triplet observed. In this case 0 and 1 were assigned based on whether the difference between BIC1Dist (ILS+introgression model) and BIC2Dist (ILS-only model) was > 10 (BIC1 > BIC2). The total introgression fraction was calculated in QuiblR as (mixprop2*count*isDiscordant)/totalTrees, with count corresponding to the number of trees supporting that topology, and total trees the total number of input gene trees (n = 190). The significant fraction (fraction of trees supporting with deltaBIC > 10 the discordant topology) was calculated as (isSignificant * count * isDiscordant) / totalTrees. For a detailed description of other columns please see QuIBL's manual at <https://github.com/miriammiyagi/QuIBL>. Sample information is available in Table S1.

**Table S5.** Individual plastid-gene support for the two alternative topologies at the *L. bienne* node retrieved from the literature. For alternative topologies and relative references also see Table S6. Out of the 36 plastid genes, 20 genes supported topology 1, while 15 genes support topology 2, based on likelihood, and one gene did not support any of the two topologies. This results in genes supporting topology 1 being only 0.3 times more frequent than genes supporting topology 2. However, considering informative genes only (delta LL > 1.5; all but one were longer than 550bp), the genes supporting topology 1 are 1.5 times more frequent than genes supporting topology 2. Indeed, the AU (Approximately Unbiased) test run in IQtree does not rule out topology 2 as a plausible topology for this node based on the markers used. Overall, this suggests that retrieval of short plastid sequences (or sampling of a small number of plastid genes) and sampling of specific genes might result in different topologies which might explain variation in topology found at this node in previous studies. Interestingly, matk, one of the genes sampled here and in previous studies was quite uninformative for topology 1 VS topology 2 (not shown).

**Table S6.** Comparison of published phylogenetic topologies at selected nodes within *Linum*. Comparisons were drawn based on: this study, McDill et al. (2009), Bolsheva et al. (2017), Bolsheva et al. (2022), Ruiz-Martin et al. (2018), Villalvazo et al. (2022), Patel et al. (2026). Comparisons concerned conflicting nodes found in *Linum* based on the Angiosperm353 phylogenetic tree shown in Figure 1 and/or other studies. Other studies were considered to be included in this comparison (Fu & Allaby, 2010; Fu et al. 2016), but these did not present all the species of interest or phylogenetic relationship in a tree format. Further studies (i.e. Maguilla et al. (2021)) used the same markers, samples, and methods to retrieve the phylogenetic trees as in Ruiz-Martin et al. (2018), so they were redundant and were not included. Also, Villalvazo et al. (2022) et al. used the same markers as Ruiz-Martin (2018), but they present phylogenetic trees obtained both by combining nuclear and plastid markers (the unique presentation in Ruiz-Martin, 2018), as well as keeping them separated. Bolsheva et al. (2017) and Patel et al (2026) did not include all species in (A), McDill (2009) did not include all species in (B), and Sveinsson et al. (2014) did not include all species in (C), so they are missing from (A), (B), and (C), respectively. Furthermore, topologies that were supported by one study only are not shown. This happened for Ruiz-Martin et al. (2018) in (B) for the chloroplast+ITS based phylogenetic tree, and for Villalvazo et al. (2022) in (B), where species were paraphyletic, and (C) for the chloroplast-based phylogenetic tree. (*) means that both chloroplast and nuclear markers were used together for phylogenetic inference. (°) appears when multiple samples were included for the same species in the phylogenetic tree, and some deviant samples occurred in the phylogenetic tree relative to the topology presented here. Specifically, Ruiz-martin et al. (2018) uses multiple *L. tenue* samples and one falls far away from the other *L. tenue* samples (not considered here). Bolsheva et al. (2017) uses multiple L. perenne samples and one also falls far away from the other *L. perenne* samples in the phylogenetic tree (not considered here). Overall, it seems like the topology retrieved by Ruiz-Martin et al. (2018) is mostly due to the ITS marker sampled which based on this summary seems to pull the topology of the tree to the alternative topology, since topologies retrieved by Villalvazo et al. (2023) based on solely plastid markers retrieve the topology encountered in this and other studies. Moreover, studies that make use of a higher number of nuclear and plastid markers (this study, Sveinsson et al. (2014), Patel et al. (2026)) always converge to the same topology at these conflicting nodes.
