## Supplementary Figures for "Reticulate evolution and climate driven diversification shaped the origin and geographic structure of *Linum bienne*"

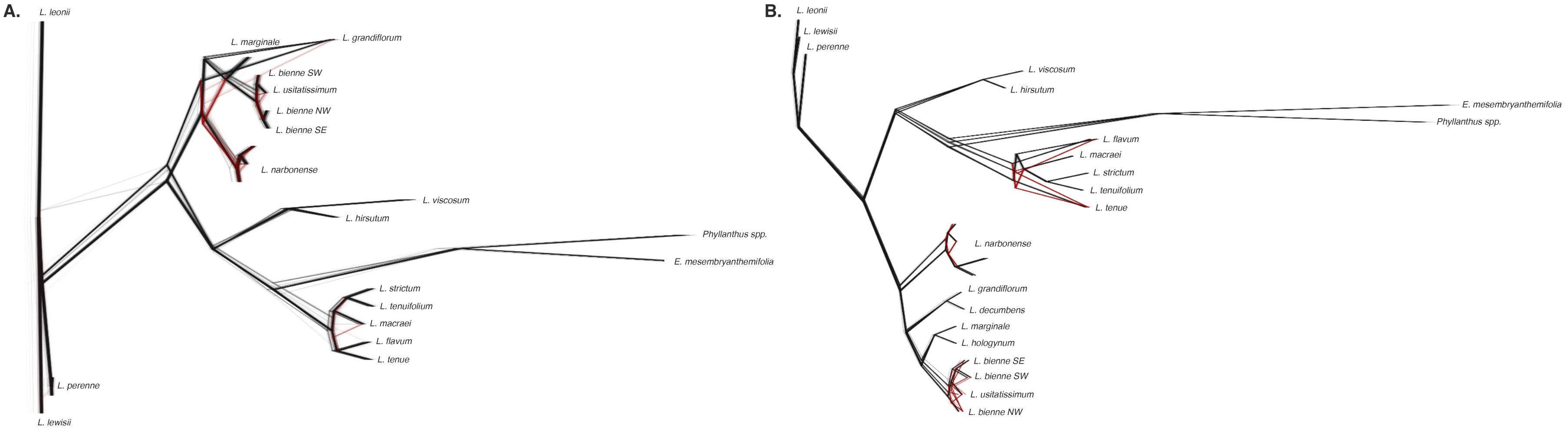


**Figure S1.** Bootstrap-tree conflict in plastid and nuclear genus-level phylogenies of Linum. DensiTree plots summarizing 1000 unrooted bootstrap trees for the (A) plastid and (B) nuclear Angiosperms353 supermatrix datasets. Branches in black follow the main consensus topology, whereas branches in red represent alternative topologies recovered among bootstrap replicates. The plastid dataset showed conflict around the L. bienne node, both among species and among L. bienne and L. usitatissimum accessions. The nuclear dataset showed conflict mainly among L. bienne and L. usitatissimum accessions, and around the L. flavum node. Supplementary Table 1 includes information about all samples.

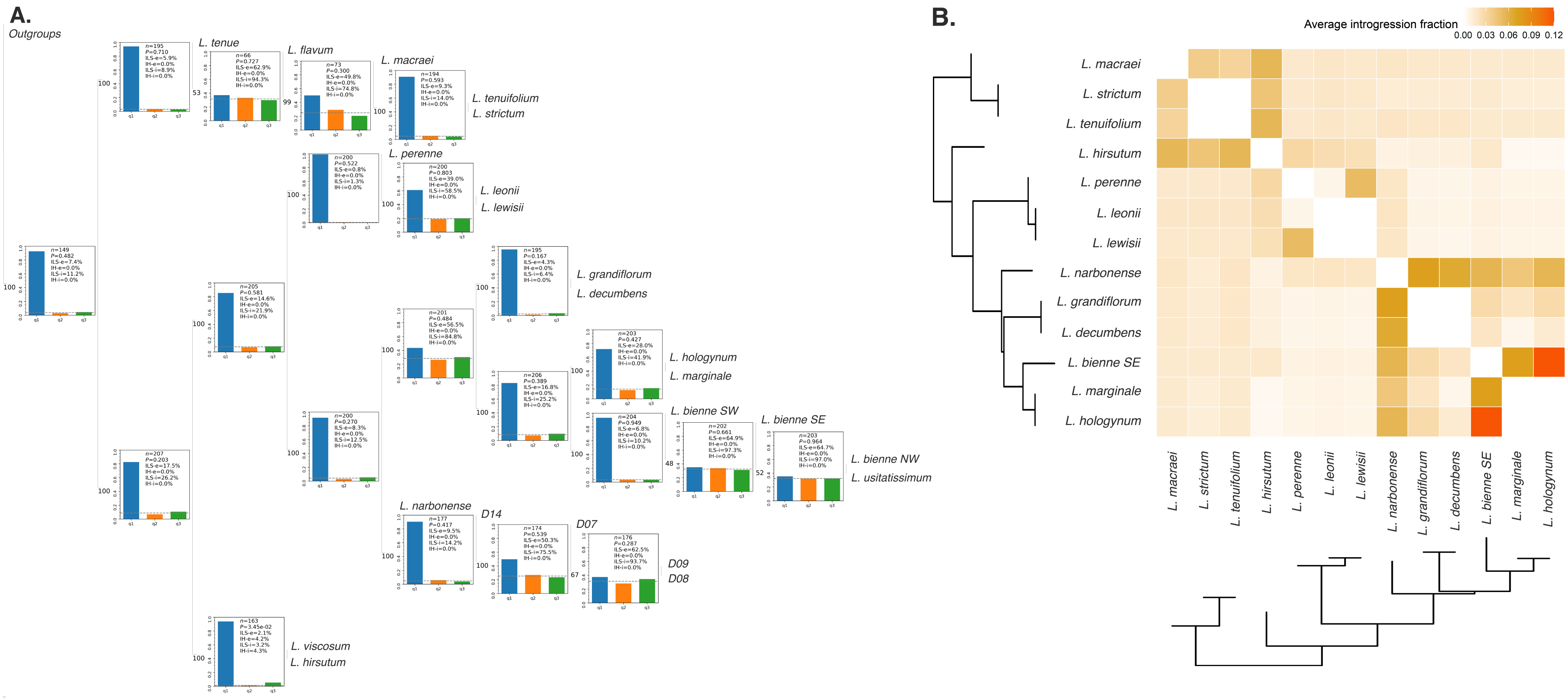


**Figure S2.** Incomplete lineage sorting and introgression across genus-level Linum sampling. (A) Phytop quartet-topology analysis using the ASTRAL nuclear tree as the guide tree. Bars show the relative frequency of the three most common quartet topologies across nuclear gene trees for each node. Similar frequencies of the three topologies are consistent with incomplete lineage sorting, whereas prevalence on the main topology and a single alternative topology may indicate hybridization or introgression. (B) QuIBL analysis testing whether gene-tree discordance is better explained by incomplete lineage sorting alone (null model) or by incomplete lineage sorting plus introgression (alternative model), using branch-length information from nuclear gene trees. Average introgression fractions are shown by tile colour, with darker colours meaning higher introgression fraction. Statistically supported ILS+introgression models were recovered only for triplets involving the L. bienne node. Supplementary Table 1 includes information about all samples.

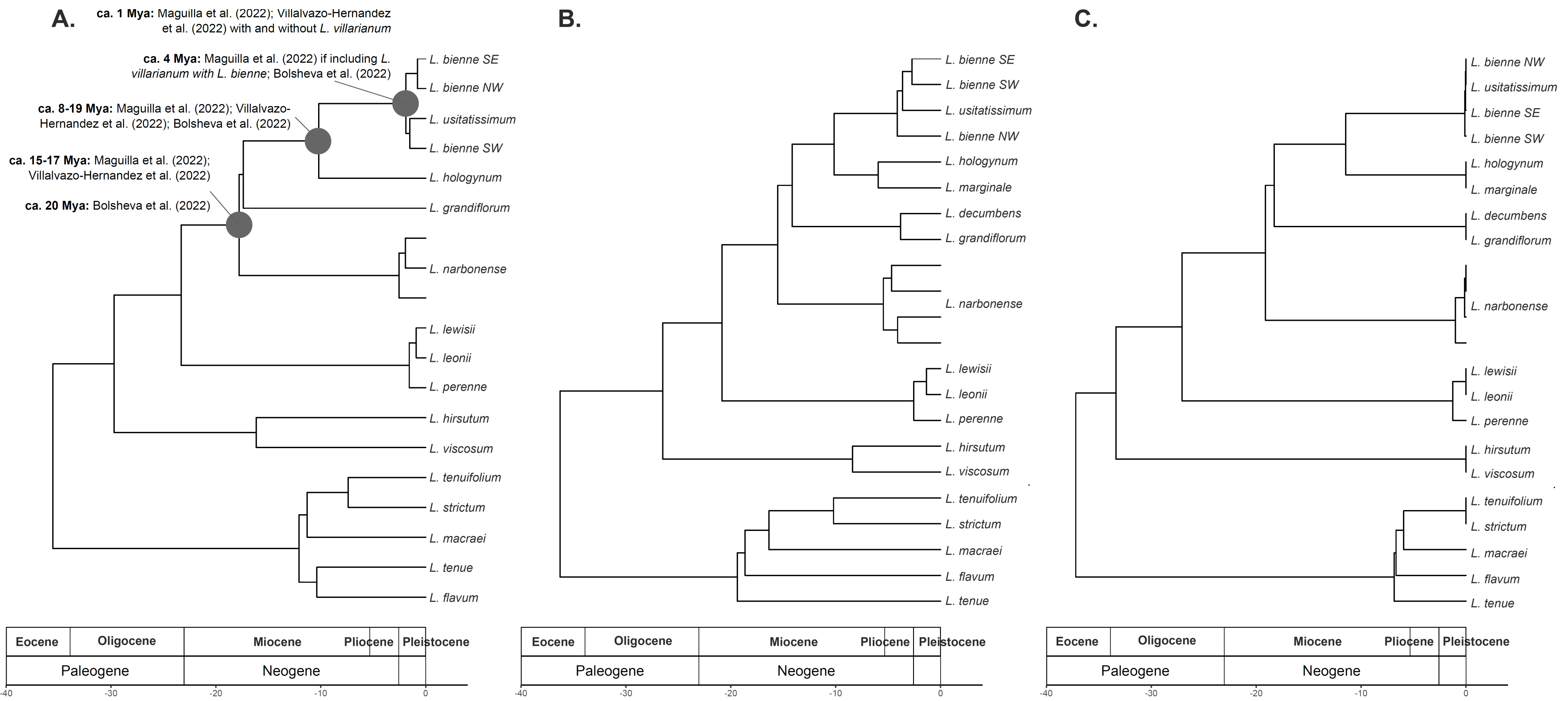


**Figure S3.** Divergence-time estimates for genus-level plastid and nuclear phylogenies of Linum. Penalized-likelihood dated phylogenies inferred from (A) the plastid supermatrix, (B) the nuclear Angiosperms353 supermatrix, and (C) the nuclear ASTRAL tree. The same fossil calibration was applied to the crown node of Linum in all analyses. Major nodes within Linum were generally placed within similar broad time windows, but age estimates differed for the L. bienne crown node: the ASTRAL nuclear tree recovered the youngest estimate, the nuclear supermatrix recovered the oldest estimate, and the plastid tree recovered an intermediate estimate. Boxes in (A) indicate comparable age estimates from previous studies for nodes relevant to the evolutionary history of L. bienne. Supplementary Table 1 includes information about all samples.

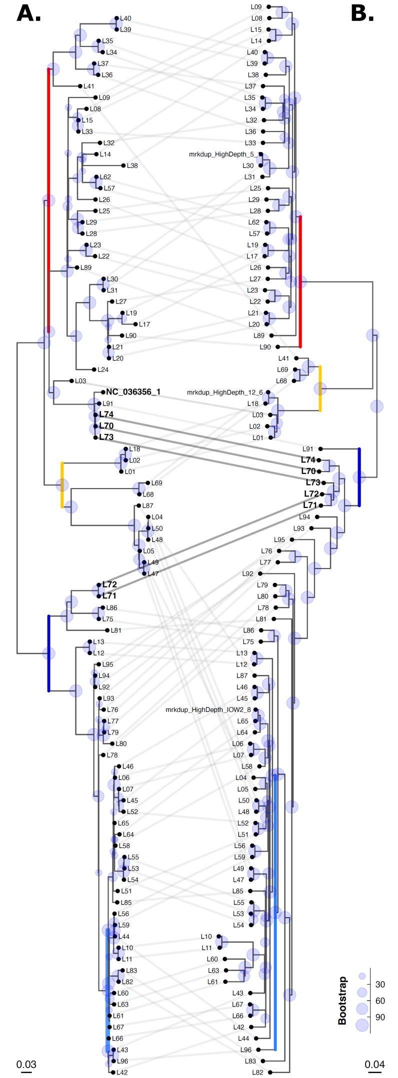


**Figure S4.** Plastome and nuclear genomic relationships among *Linum bienne* and *L. usitatissimum* accessions. (A) Plastome phylogeny inferred with maximum likelihood in IQ-TREE (GTR+G). (B) Nuclear genetic-distance tree inferred with ngsDist from low-depth nuclear genome data. *Linum narbonense* was used as outgroup in both analyses (outgroup was pruned for representation purposes). Bootstrap (1000) is represented by light blue circles. Light grey lines connect the same individuals between trees; darker lines indicate *L. usitatissimum* individuals. Colours indicate the geographically structured lineages recovered in the plastome and nuclear datasets: SW (red), Central (yellow), SE (dark blue) and NW (light blue), with the SW+central sister to the SE+NW, and NW embedded within SE. Most individuals were assigned to the same lineage in both datasets, but several showed cytonuclear discordance, with plastid and nuclear data assigning the same individual to different lineages. Supplementary Table 3 includes information about all samples.

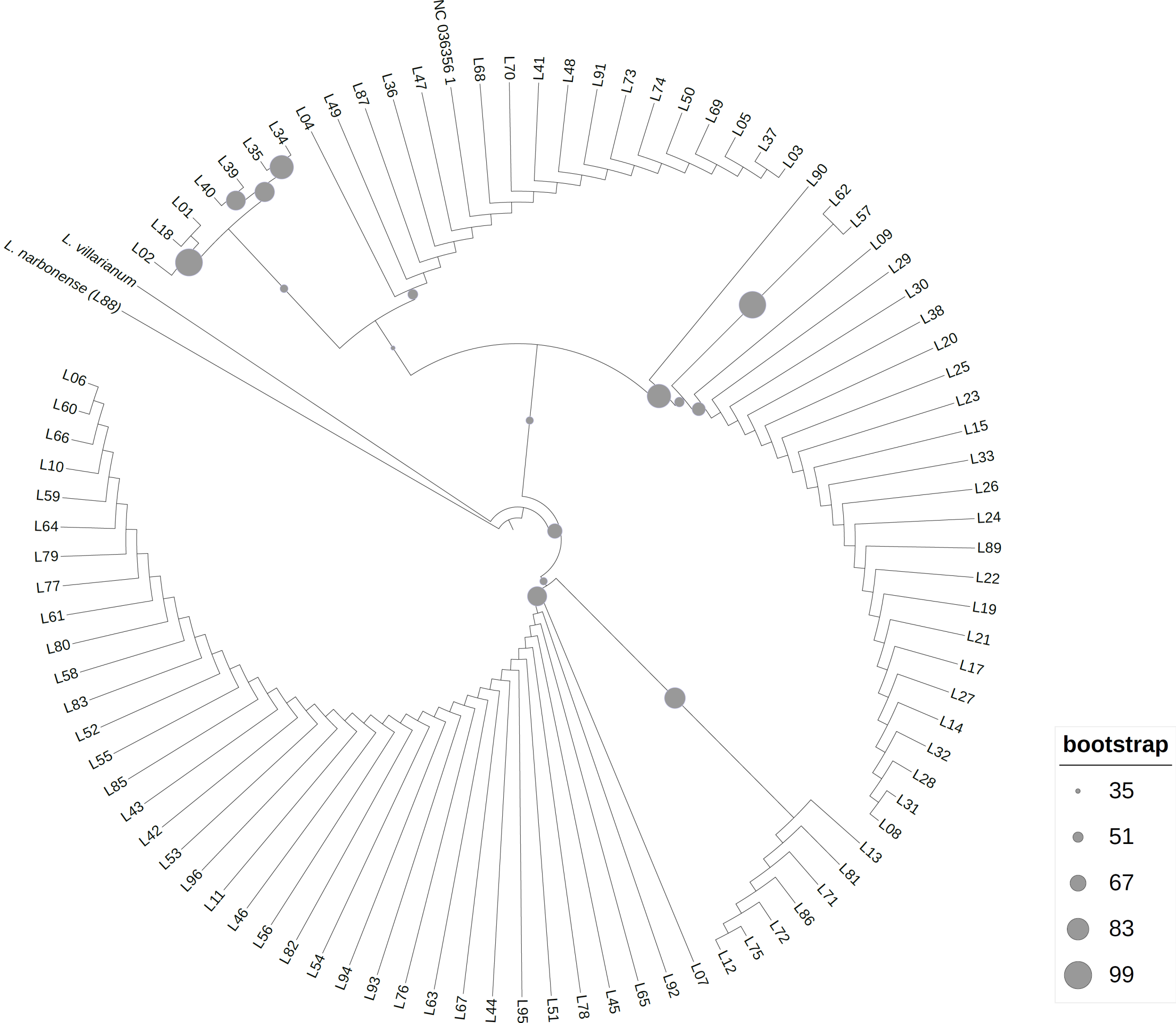


**Figure S5.** Plastid-gene phylogeny including *Linum villarianum*. IQtree maximum-likelihood phylogeny (GTR+G) inferred from a reduced plastid-gene alignment including *L. bienne* individuals and one *L. villarianum* accession retrieved from published sequence data. The analysis was used to assess whether *L. villarianum* falls within any of the major *L. bienne* lineages or is recovered as sister to *L. bienne*. Supplementary Table 3 includes information about all *L. bienne* samples.

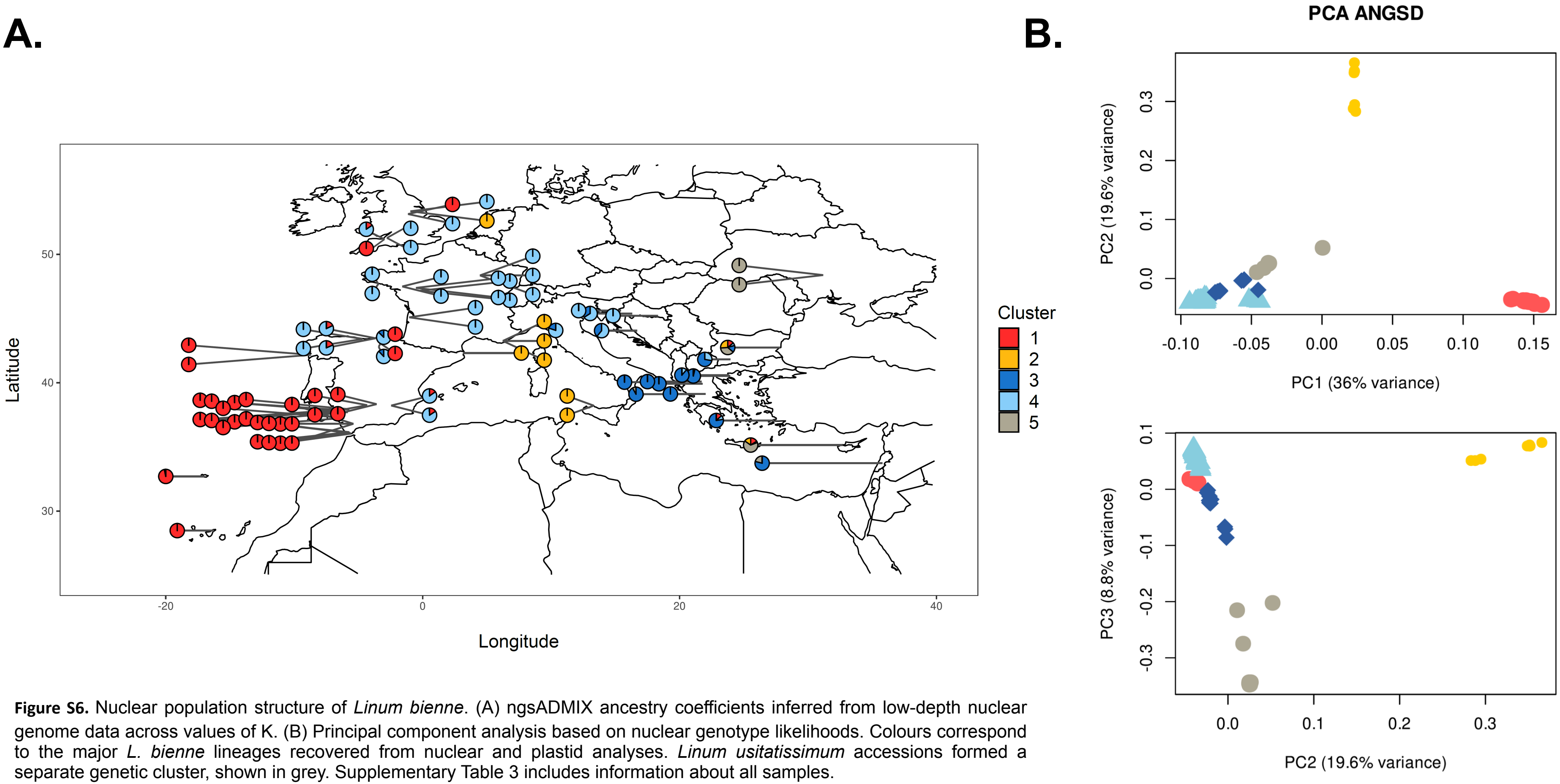


**Figure S6.** Nuclear population structure of *Linum bienne*. (A) ngsADMIX ancestry coefficients inferred from low-depth nuclear genome data across values of K. (B) Principal component analysis based on nuclear genotype likelihoods. Colours correspond to the major *L. bienne* lineages recovered from nuclear and plastid analyses. *Linum usitatissimum* accessions formed a separate genetic cluster, shown in grey. Supplementary Table 3 includes information about all samples.

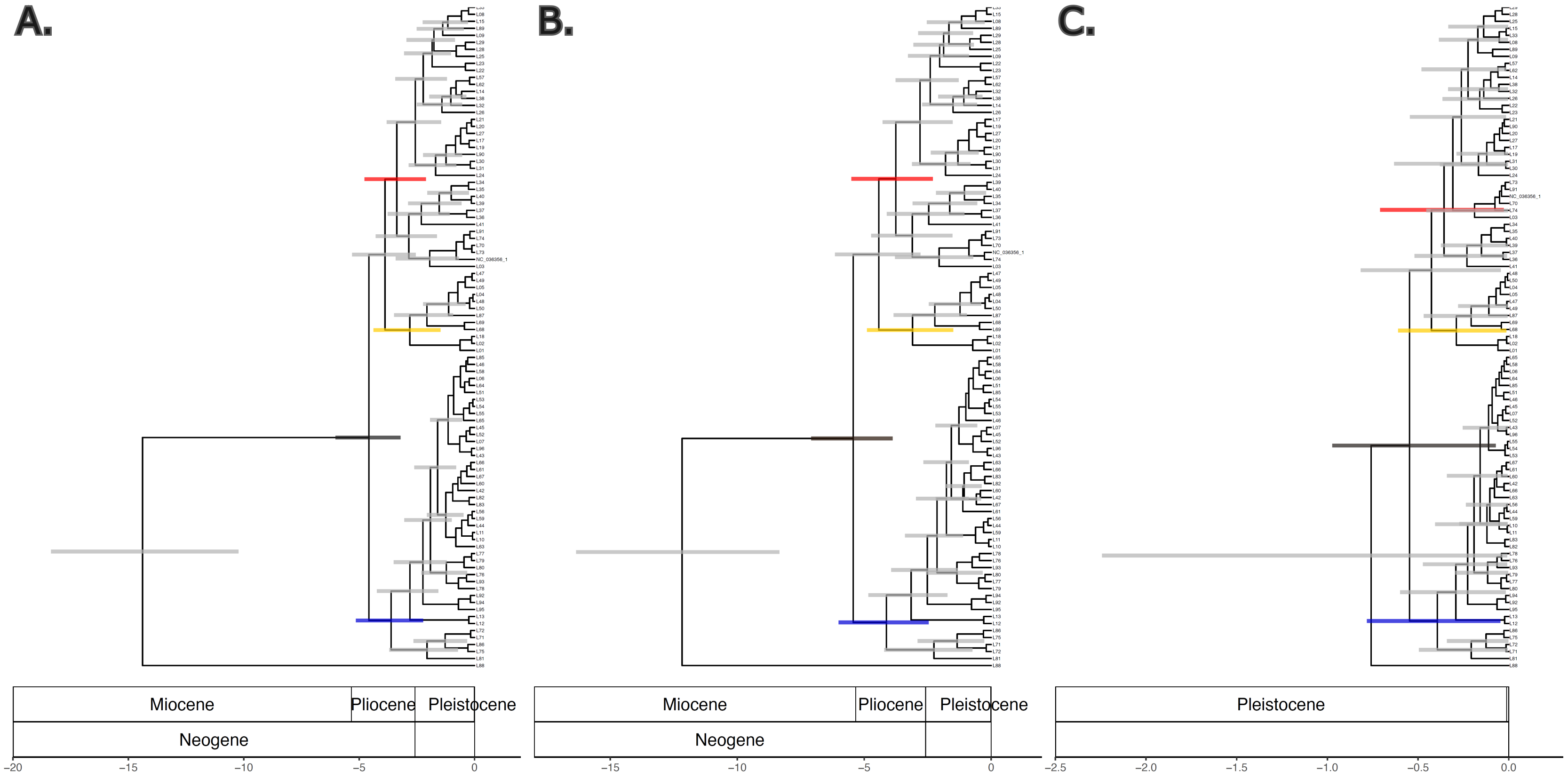


**Figure S7.** Plastome-based BEAST dating of Linum bienne lineages. Maximum clade credibility trees inferred from the L. bienne and L. usitatissimum plastome alignment, using L. narbonense as outgroup. Three BEAST analyses were run using alternative secondary calibration points derived from genus-level treePL analyses based on (A) plastid, (B) nuclear supermatrix, and (C) nuclear ASTRAL phylogenies. Horizontal bars indicate uncertainty around node-age estimates. Plastid and nuclear-supermatrix calibrations produced older age estimates than the nuclear-ASTRAL calibration. Overall, it was estimated that L. bienne started to diverge from other Linum species between the early to mid-Pliocene. Horizontal bars represent posterior probabilities for the age of each node and they are coloured with: black (L. bienne crown node), red (southwestern lineage), yellow (central lineage), blue (southeastern + northwestern lineages), grey (any other node). Supplementary Table 3 includes information about all samples.

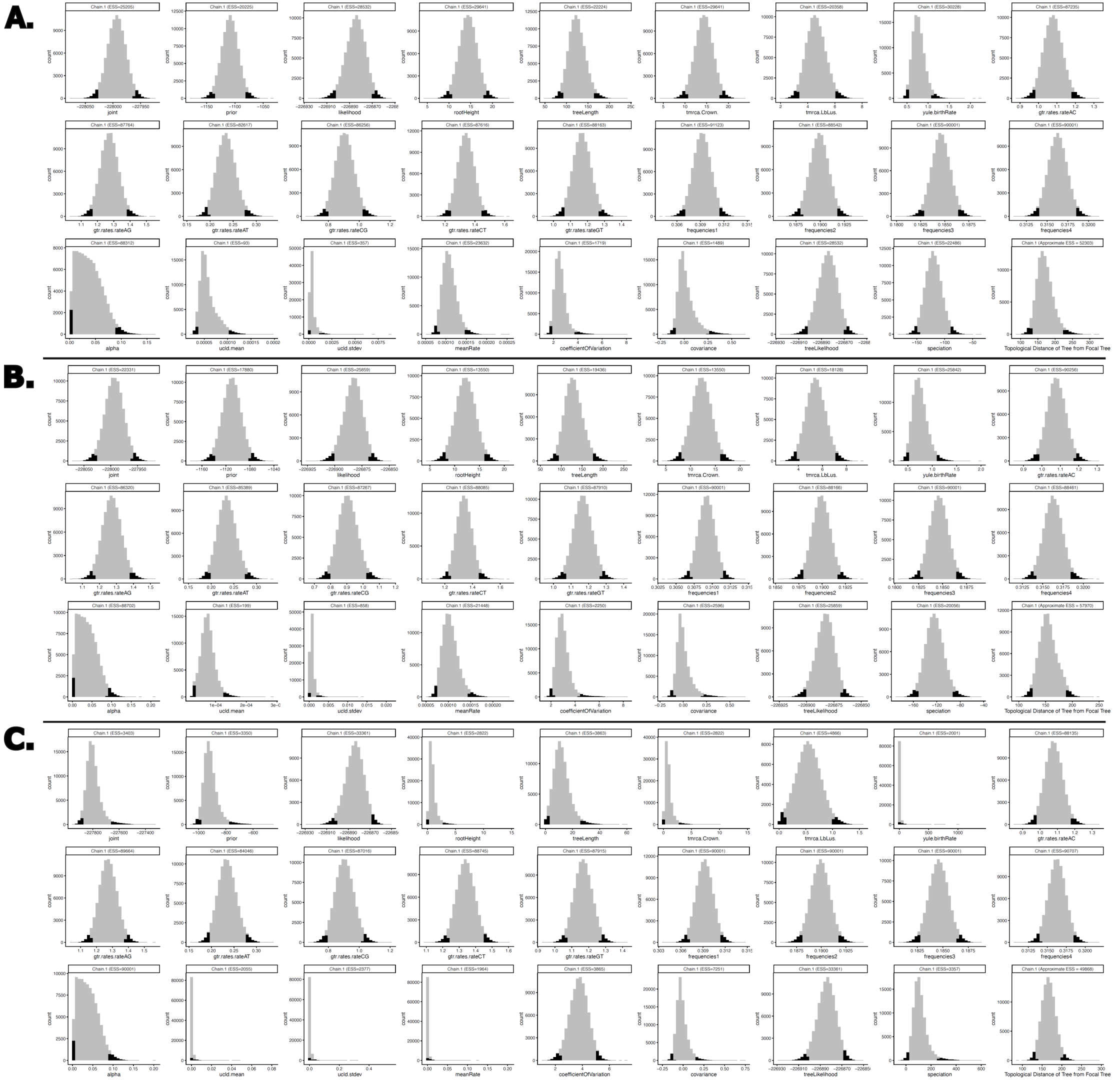


**Figure S8.** BEAST parameter distributions and convergence diagnostics. BEAST parameter distribution from 1,000,000,000 runs (10% burn-in in black) for the dating of the *L. bienne* plastome phylogeny using differing secondary calibration points for the *L. bienne* crown node and the overall root using *L. narbonense* as outgroup. Parameters are shown for BEAST run with secondary calibration points from the *Linum* phylogenetic trees dated with treePL based on: plastid (A), nuclear supermatrix (B), and nuclear-Astral (C) datasets. All three types of calibration resulted in high ESS (>200) for all parameters, except UCLD mean and standard deviation for the plastid calibration which had ESS < 200.

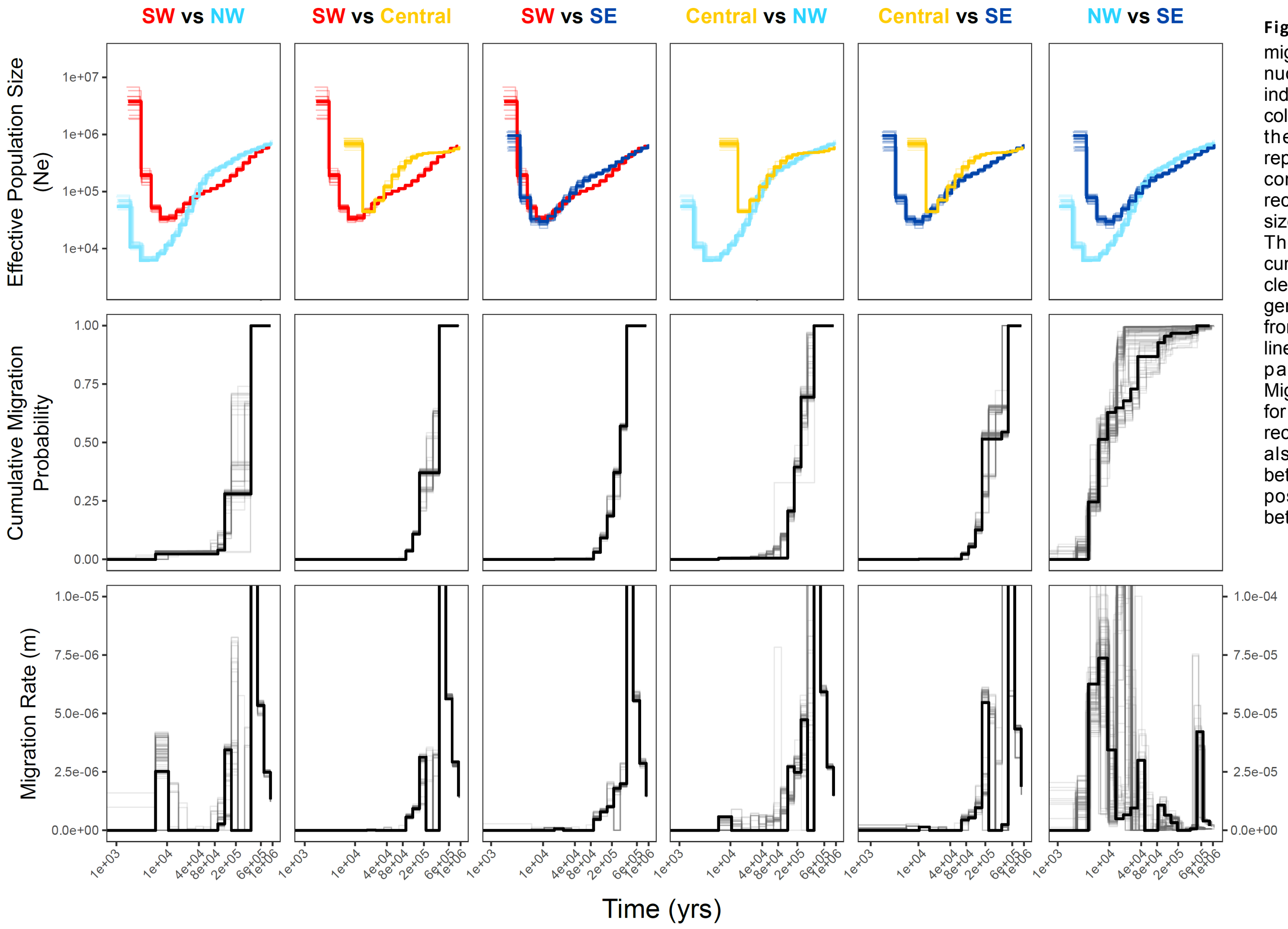


**Figure S9.** Demographic history and migration inferred with MSMC-IM based on nuclear genomes. Genetic clusters are indicated with colours and correspond to columns. Time (ca. 1Mya to present) is on the x-axis. The first row of panels represents effective population size for all combinations of clusters with lineages recovering from a large drop in population size around 10000 years ago (Holocene). The second row of panels represent cumulative migration probability, which is clearly higher between the NW and SE genetic clusters, but in general decreases from past to present highlighting when lineages started to diverge. The third row of panels represents migration rates. Migration rates are again quite high again for the NW and SE genetic clusters even in recent times, however a smaller spike was also found around the mid-Holocene between the SW and NW lineages, and possibly the same pattern is present between the central and NW lineages.

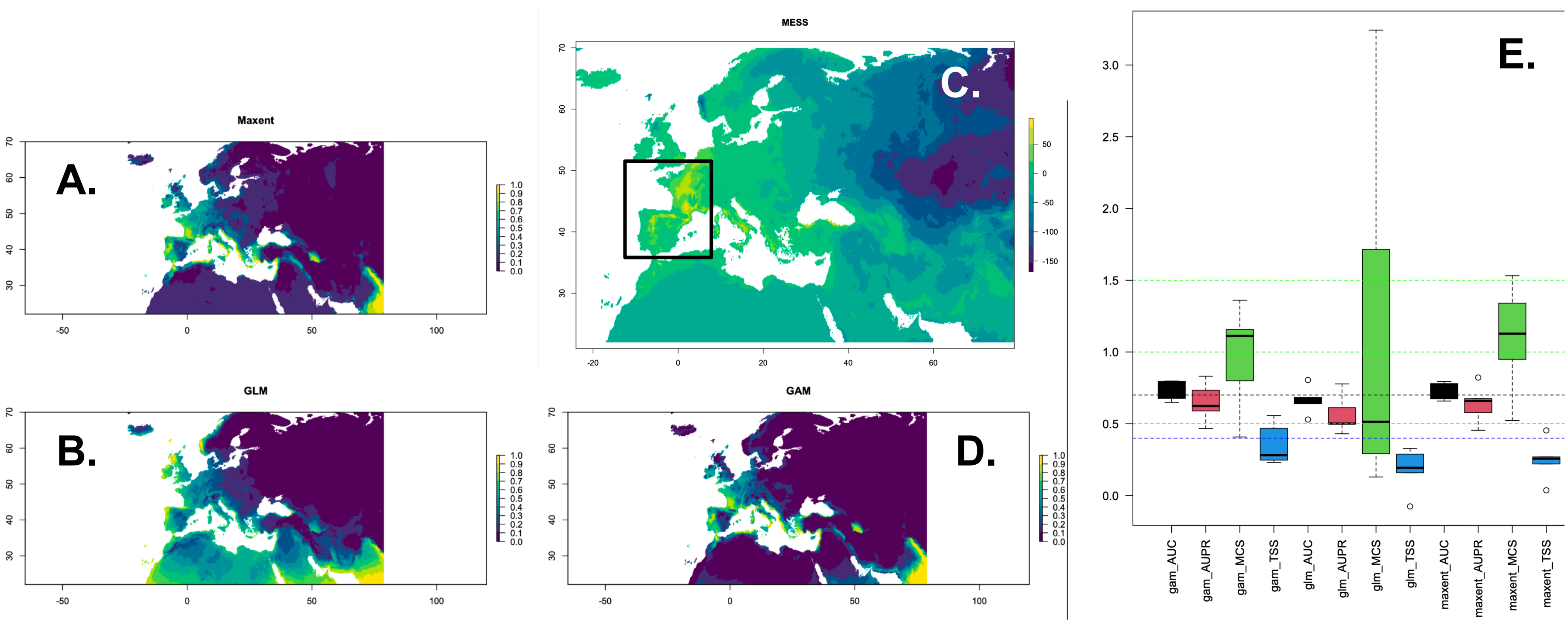


**Figure S10.** Present-day environmental niche model performance for Linum bienne. Present-day climatic suitability or favourability predicted using (A) Maxent, (B) GLM and (D) GAM models fitted with L. bienne occurrence data extracted from GBIF and WorldClim bioclimatic variables. (C) MESS was used to show climatic dissimilarity between the model training region (indicated by a square and including the Iberian Peninsula and France, excluding islands) and the broader projection area. In the analysis, low values (below zero – blue) indicate the highest dissimilarity in climate between the training area and the predicted area, while high values (above zero – yellow) indicate high climate similarity. Model performance assessed using AUC (> 0.7), MCS (0.5 < MCS < 1.5), and TSS (> 0.4). GAM and Maxent showed broadly similar performance, with GAM selected for past-climate projections, while GLM performed the worst. AUC = Area Under the Curve; MCS = Miller Calibration Score; TSS = True Skill Statistic.

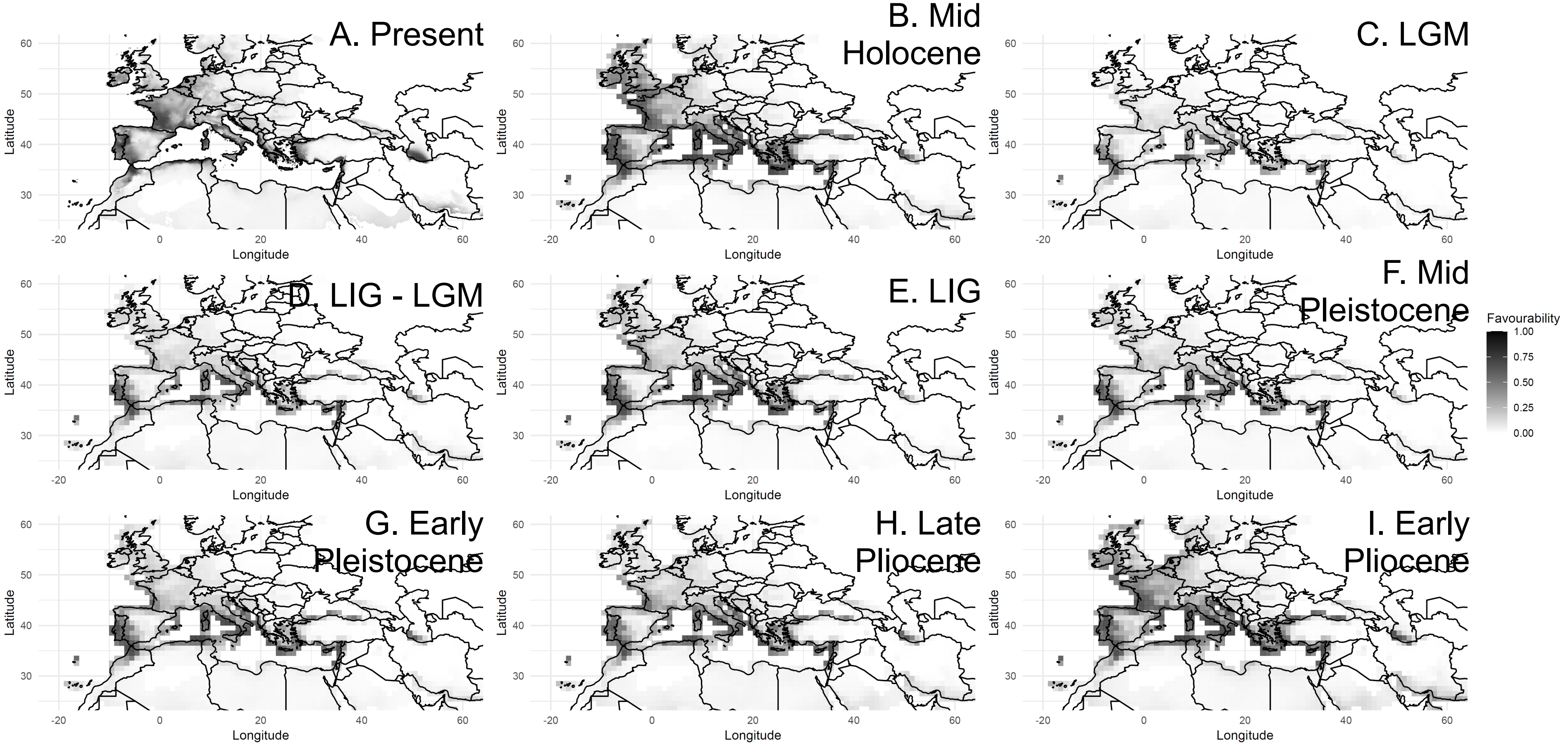


**Figure S11**. Past and present climatic favourability for Linum bienne. GAM-based projections of climatic favourability for L. bienne (using GBIF occurrence https://doi.org/10.15468/dl.nhbuhr accessed on 03/07/2025, and WorldClim and pastclim bioclimatic variables) from the present to the early Pliocene. Panels show: present (A), Mid-Holocene – 8000 years ago (B), Last Glacial Maximum – 21000 years ago (C), transition between Last Interglacial and Last Glacial Maximum – 70000 years ago (D), Last Interglacial – 130000 years ago, Mid Pleistocene – 700000 years ago (F), Early Pleistocene – 1300000 years ago (G), Late Pliocene – 2700000 years ago (H), Early Pliocene – 4500000 years ago (I). Darker colours indicate higher predicted favourability, whereas lighter colours indicate lower predicted favourability. Across periods of reduced favourability, northern and higher-latitude regions were generally less favourable than southern Mediterranean regions.

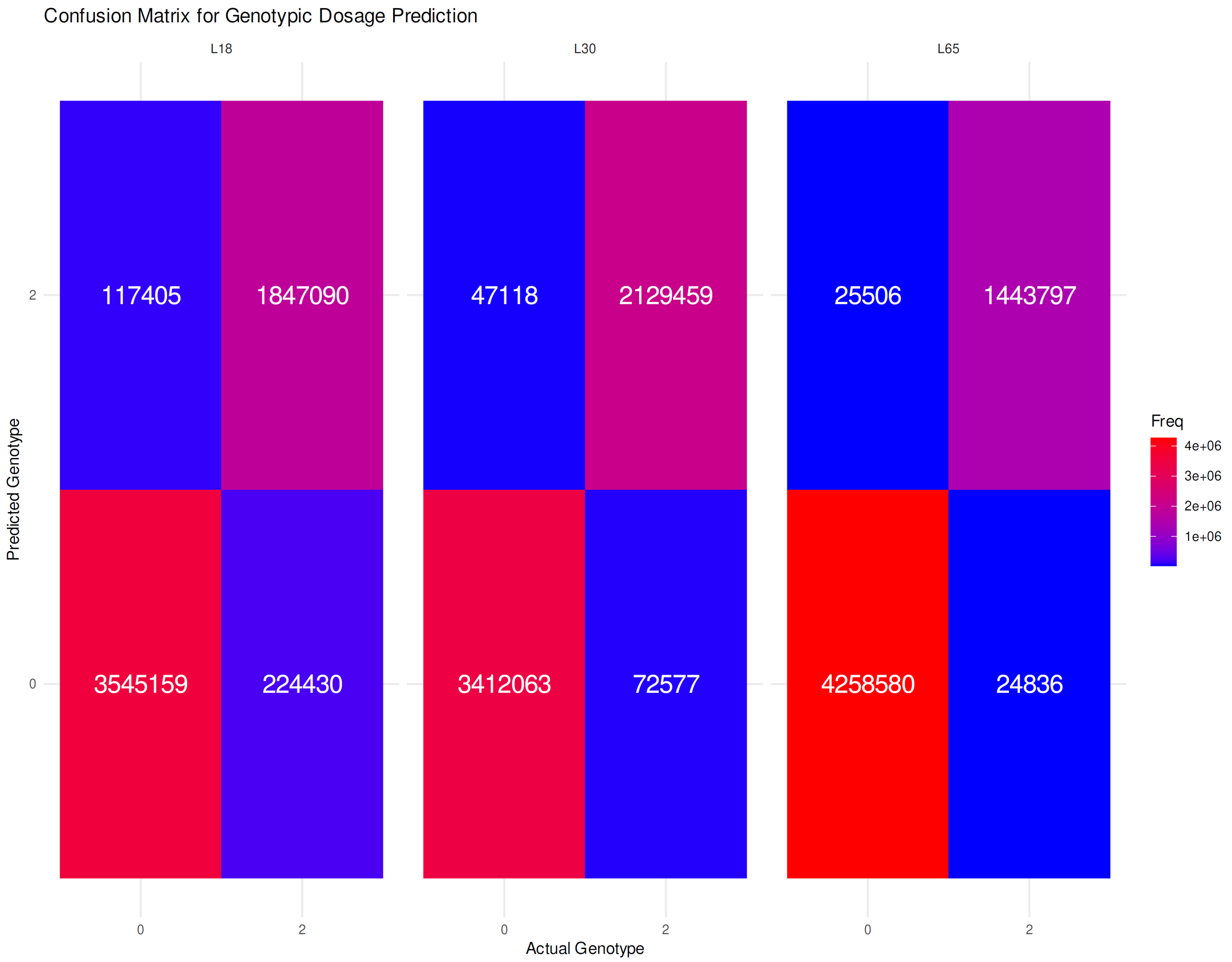


**Figure S12**. Accuracy of genotype imputation with STITCH. Confusion matrix comparing observed high-depth genotypes with genotypes imputed from low-depth data using STITCH. The x-axis shows observed genotypes and the y-axis shows imputed genotypes. Warmer colours indicate more genotype calls. Higher counts along the diagonal indicate that imputed genotypes generally matched observed genotypes.
