## Supplementary Methods for "Reticulate evolution and climate driven diversification shaped the origin and geographic structure of *Linum bienne*"

**S1. Hybpiper pipeline details for Angiosperms353 nuclear loci and plastid loci**

The HybPiper pipeline (v2.3.2; Johnson et al., 2016) was used to recover plastid and single-copy nuclear genes (Johnson et al., 2019) from 19 *Linum* species represented by 26 individuals (Table S1).

For Angiosperms 353 nuclear genes (Johnson et al., 2019), the mega353 bait file (<https://github.com/chrisjackson-pellicle/NewTargets>) was enriched in silico for Malpighiales-specific baits using the function filter_mega353.py found at the same repository which should have improved retrieval of Angiosperms353 loci (MacLay et al., 2021). After running HybPiper on nuclear data, we recovered on average 150 Kbp for nuclear genes (≈65% of the Angiosperms353 targets). Two individuals with <50 genes retrieved at 50% sequence length were excluded (samples C95, D04), but all 19 intended species remained represented (Table S1-S2). Further, genes were excluded from subsequent analyses: if their retrieved length was < 400bp in at least half of the samples, if they were not retrieved for any of the two outgroups, and if hybpiper flagged them with a paralog warning across more than one taxon. Paralog warnings were especially frequent in *L. bienne, L. usitatissimum, L. marginale, L. hologynum, L. macraei* (Table S1-S2). This finally resulted in 208 nuclear genes out of 353 and 24 samples out of 26 retained for subsequent analyses.

For plastid genes, the bait file available at https://github.com/mossmatters/plastidTargets (Pokorny et al., 2024) was used for sequence retrieval of 72 genes. After running HybPiper, we retained only individuals for which at least 20 genes were recovered for at least 50% of their length (samples D06, D04, D08 were discarded; Table S1-S2). However, although *L. hologynum* and *L. decumbens* (samples D02, D03) recovered 30 genes or more at 50%, the percentage on target was among the lowest for retained samples (0.3%; Table S1-S2) and the two individuals formed a monophyletic group subtending a long branch if kept for phylogenetic inference (not shown) also these were discarded. Further, genes were excluded from subsequent analyses using the same criteria as for nuclear genes. This resulted in 36 out of 72 plastid genes and 21 out of 26 individuals for subsequent analyses.

**S2. Assessing the nature of conflicting phylogenetic signal across *Linum* and for the *L. bienne* node using Angiosperms353 and plastid genes**

We assessed the extent and likely origin of conflicting phylogenetic signals in *Linum*. implementing complementary approaches using the nuclear Angiosperm 353 gene trees. Phytop v0.3.2 (Hong-Yun et al., 2025) and QuIBL (Edelman et al., 2019) were used to explore whether nuclear gene-tree discordance was more consistent with incomplete lineage sorting (ILS) or with hybridization/introgression at the genus level. Phytop evaluates discordance from quartet topology frequencies, whereas QuIBL incorporates branch-length information. We then used PhyloNet v3.8.2 to further explore patterns of reticulation at the *L. bienne* node (Table S1) (Than et al., 2008). These methods are usually applied to larger datasets (> 500 gene trees; Morales-Briones et al., 2022; Cui et al., 2025; Xu et al., 2024; Wu et al., 2024) and might be sensitive to sample size, sequence length, high mutation rate variation, and/or less effective at confidently identifying introgression when ILS is high (Yu & Nakleh, 2015; Solis-Lemu & Ane, 2016; Hong-Yun et al., 2025; Yan et al., 2026; Galtier, 2024). We therefore interpret these analyses as evidence for the presence of dominant sources of phylogenetic discordance, rather than as tests that can exclude weaker or more localized signals of introgression.

For all analyses, the ASTRAL coalescent tree was used as the guide species tree. QuIBL further requires all ML best gene trees produced in IQTREE (input) and the guide tree to contain the same set of species, and it is sensitive to missing data and to the inclusion of multiple individuals per species. Thus, all guide and input gene trees were pruned before the analysis so that all trees retained had one representative per species and *E. mesembryanthemifolia* as outgroup. This also resulted in dropping some of the gene trees because some did not contain all taxa of interest (190 gene trees were retained out of 208). Consequently, only individuals appearing in at least 190 gene trees were retained. This filtering removed individuals AXPJ, D08, D09, D14, D10, D11, D12, while retaining all sampled *Linum* species (Table S1). Because QuIBL does not analyse sister lineages directly, the southeastern *L. bienne* accession was retained as a placeholder for the *L. bienne–L. usitatissimum* lineage.

Both QuIBL and Phytop suggested ILS was predominant relative to hybridization or introgression at the genus level (Figure S2, Table S4). However, at the *L. bienne* node (including *L. narbonense, L. decumbens, L.grandiflorum, L. hologynum, L. marginale, L. bienne)*, QuIBL identified triplets for which a model including both ILS and introgression was statistically supported. Most of these triplets had *L. bienne* as an outgroup, suggesting that its position was not fully resolved and could reflect a history of mixed ancestry. Moreover, previous studies have reported *L. bienne* and *L. usitatissimum* to be ancient or meso-allopolyploids with ancestors closely related to *L. decumbens-L. grandiflorum*, and/or *L. narbonense* (Bolsheva et al., 2017; Wang et al., 2012). Therefore, we applied a phylogenetic network approach – PhyloNet - to test the hypothesis of *L. bienne* having hybrid origin. For this, we used the InferNetwork_ML method, based on maximum likelihood, and a reduced set of taxa (*L. hirsutum* as outgroup, then *L. narbonense, L. grandiflorum, L. decumbens, L. bienne*) based on previous karyotype studies (Bolsheva et al., 2017). Note that *Linum hologynum* and *L. marginale* were excluded from these analyses because, although they are polyploids based on chromosome counts, their origin is unknown and their chromosome numbers differ from those of *L. bienne* and *L. usitatissimum* (Bolsheva et al., 2017; Table S1). *Linum usitatissimum*, sister to *L. bienne*, was also excluded from these analyses as *L. bienne* could work as a placeholder for both species based on our ASTRAL phylogenetic trees (see Results).

The 190 input best ML trees were pruned to contain *L. bienne, L. decumbens, L. grandiflorum, L. narbonense,* with *L. hirsutum* set as outgroup. The ASTRAL tree was set as the starting species tree, with 10 iterations for each run. To better capture the species diversity, *L. bienne* was represented by the southeastern and southwestern individuals (Table S1) and assigned to the same species using the -a option in PhyloNet. PhyloNet was run allowing the maximum number of hybrid nodes to vary from 0 to 3. Analyses were run twice: first without specifying any hybrid taxon, and second by specifying *L. bienne* as the hybrid taxon (setting the option -h). This allowed us to test whether likelihoods improved when *L. bienne* was constrained as deriving from a hybrid node, rather than allowing PhyloNet to place hybrid nodes freely. After running PhyloNet, log likelihoods were examined for both runs, to assess which number of hybrid nodes resulted in the optimal network solution. Adding one or two hybrid nodes improved the likelihood relative to the backbone tree, and a plateau was reached at three. We therefore examined the ten most likely networks for one and two hybrid nodes using PhyloNetworks (Solís-Lemus et al., 2017). Because PhyloNet can infer hybrid nodes with very low gamma values for one of the two donor lineages (gamma is the inheritance probability from the parents of the hybrid node), which might be a sign of a spurious hybrid node (for example in the presence of ILS), very ancient introgression signal, or minor gene flow (Kong et al., 2024; Morales-Briones et al., 2018; Pezzi et al., 2026). Thus, we paid particular attention to networks in which the both hybrid edges had γ > 0.1.

We also assessed whether plastid gene choice could have contributed to topological discrepancies among previous phylogenetic studies of *Linum*. Although plastomes are inherited as a single unit and should be treated as a single coalescent locus (Doyle, 2022), individual plastid genes can differ in length, substitution rate and phylogenetic signal (Walker et al., 2019; Gonçalves et al., 2019). Previous phylogenetic studies in *Linum* have used different subsets of plastid markers (Table S6), which might have contributed to differences in recovered topology. We then used our expanded plastid-gene dataset to test whether using specific plastid genes over others could lead to divergent phylogenetic signals at the same node. For plastid data, IQTREE was run again on plastid gene alignments with a GTR+G model, but this time separately for each gene and providing two alternative topologies to test based on the present and previous studies (Table S6). Differences in log-likelihood between topologies were used to assess whether particular plastid genes provided stronger support for one topology over the other. This analysis is summarized in Table S5.

**S3. Nuclear genome analyses in *Linum bienne***

*3.a. Low depth genotype likelihood estimates*

Low depth nuclear genotypes were retrieved by aligning trimmed short reads to the chromosome level assembly available for *L. usitatissimum* (ASM1066527v2 Zhang et al., 2020), using BWA MEM v0.7.17 (Li et al., 2009). The program ANGSD v0.940 (Korneliussen et al. 2014) was used to obtain genotype likelihoods and incorporate uncertainties due to low depth of coverage into downstream analyses. Filtering parameters common to all analyses were as follows: -GL 1 -uniqueOnly 1 -remove_bads 1 -trim 0 -C 50 -baq 1 -minMapQ 20 -minQ 20 -skipTriallelic 1. The SNP calling model was the same as in SAMTOOLS (GL 1) and only reads with a mapping quality of at least 20 and mapped at a single position along the genome were kept (minMapQ 20, uniqueOnly 1 and remove_bads 1). Bases with a quality lower than 20 were removed (minQ 20). No trimming of the ends of the reads was performed (trim 0), mapping quality for reads with more than 50 mismatches was adjusted (-C 50), as well as quality scores around indels (baq 1). Only biallelic loci were retained (skipTriallelic 1 ). Because L. bienne and cultivated flax are highly inbred, we incorporated inbreeding in likelihood inference. Selfing rate (*S*) in cultivated flax is in the 95 – 99% range (Dillman, 1938; Jhala et al., 2011), and we therefore set an inbreeding coefficient *F*=*S*/(2-*S*) = 0.95 using the -indF option. We generated a mask file to exclude regions of low confidence. We used genmap (Pockrandt et al., 2020) to estimate genome-wide mappability, using windows size of 50bp and allowing for up to two mismatches. Regions with a mappability lower than 1 were excluded. In highly selfing plants, heterozygosity is often found associated with regions with poor mappability or recent repeats (Jaegle et al., 2023). We identified these regions by resequencing three individuals representative of the main *Linum bienne* lineages in our analyses (L65, L30, L18). We called genotypes with freebayes v1.2.0 (Garrison & Marth, 2012) Shared heterozygosity is expected to be rare among these three high-depth individuals, given the high geographic distance of sample origin and high inbreeding. We therefore masked 5kb regions flanking SNPs found heterozygous in at least two individuals.

*3.b. Population structure and relatedness*

Polymorphic sites were filtered to have a minimal p-value of 1x10^-6^, and a maximum amount of missing data per site of 66%. Only biallelic loci were retained (skipTriallelic 1). After inspecting the distribution of depth of coverage over the entire dataset, we set a maximum bound on depth of 250X. We thinned the resulting dataset to sample one every 10kb, reducing linkage disequilibrium among markers. This resulted in 16,890 polymorphic sites distributed along the *L. bienne* nuclear genome. We used ngsDist (Vieira et al., 2016) with pairwise deletion of missing data to obtain a tree of genetic distances. We estimated branch support through 100 block-bootstrap replicates, with blocks of 20 SNPs each. We also used ngsADMIX (Skotte et al., 2013) to obtain a probability of assigning each individual to K clusters, with K = 1 to 9. We also used PCANGSD v1.36.1 (Meisner & Albrechtsen, 2018) to run a Principal Component Analysis based on genotype likelihoods. Demographic analyses are sensitive to the inclusion of closely related samples. We excluded close relatives from these analyses by estimating *R_AB_*, a relatedness coefficient that is robust to inbreeding (Hedrick & Lacy, 2015), using ngsRelate v2 (Hanghøj et al., 2019). *R_AB_* ranges from 0 (unrelated individuals) to 1 (clones). We excluded one individual of each related pair if they were above the 2^nd^ degree relative threshold (*R_AB_* > 0.25), retaining the individual with the highest depth of coverage for downstream demographic analyses. We assigned individuals to a genetic cluster if they showed a percentage of assignment larger than 60%. *Linum bienne* being a highly selfing species, individual heterozygosity cannot be used as a proxy for local genetic diversity. To visualize patterns of genetic diversity in space, we used the matrix of pairwise genetic distances generated by ngsDist and computed the average genetic distance between all individuals found in a predefined radius of five degrees, excluding putative clones (*R_AB_* > 0.75).

*3.c. Imputation of missing genotypes.*

To examine rates of (cross-)coalescence, we used MSMC-IM (Wang et al., 2020), which require phased, non-missing genotypes. To maximize the amount of information retrieved from low-depth genomes, we used STITCH v1.7.3 (Davies et al., 2016), a reference-panel–free imputation method that leverages haplotype structure and genomic alignments to infer genotypes from low-depth sequencing data. Imputation accuracy was assessed using high-depth resequencing data from three accessions (L65, L30, L18) representing the main *L. bienne* lineages in our analyses. We compared genotypes imputed from low-coverage genotypes to their high-coverage replicates to produce a confusion matrix, and estimated the correlation coefficient (*r²*) between imputed and true genotypes. STITCH accuracy is mainly governed by two parameters: *K*, the number of founder haplotypes, and *S*, the number of replicates used to average inference. We set the nGen parameter to 4 × *Nₑ* / *K* (with a long-term value for *Nₑ* = 50,000) to approximate the recombination rate between ancestral haplotypes. STITCH is expected to be robust to moderate misspecification of this parameter. We tested combinations of *K* (4–32) and *S* (1–10), and retained the configuration that optimized both runtime and imputation accuracy (S=10, K=16). Because of high selfing rates in flax, we used the diploid-inbred model, which assumes homozygous genotypes. Imputation showed good performance, with correlation coefficients between actual and imputed genotypes reaching 0.85 (Figure S12). We considered genotypes as already phased, with an allele dosage ≥1 and < 1 as homozygous for the alternate allele and the reference allele, respectively.

*3.d. Estimating the genome-wide substitution rate*

Demographic analyses require a substitution rate to obtain parameters such as times and effective population sizes. However, the set of nuclear baits used for phylogenetic reconstruction consists of highly conserved markers, and may not reflect the genome-wide nuclear rate. We took advantage of the recently published genomes of *Linum tenue* (Gutiérrez-Valencia et al., 2022)*, L. lewisii* (Innes et al., 2023)*,* and *L. trigynum* (Gutiérrez-Valencia et al., 2024)*,* a close relative to *L. tenue,* to obtain a substitution rate based on our divergence time estimates. We used LASTZ v1.04.22 (Harris, 2007) to obtain pairwise alignments of the genomes, comparing *Linum tenue* and *L. trigynum* to *L. usitatissimum* and *L. lewisii*. For the four pairs, we estimated the modal divergence for aligned fragments longer than 5kb, and divided the result by two times the time since divergence inferred in our phylogenetic analysis.

*3.e. Demographic inference based on the allele frequency spectrum (fastsimcoal2)*

Recent demographic events may be difficult to quantify under a phylogenetic framework. We therefore performed demographic inference and model comparison under the likelihood framework developed in fastsimcoal2.8 (Excoffier et al., 2013), using frequency spectra inferred by angsd from nuclear data. We obtained the spectra using ANGSD, filtering sites not found in at least 66% of individuals assigned to a lineage, and filtering sites with an average depth > 3X. We did not include the central lineage nor *L. usitatissimum* in this analysis, due to their low sample size after removing close relatives, which would have prevented an accurate estimate of the frequency spectrum. We extracted the joint minor allele frequency spectrum (MSFS) using WINSFS (Rasmussen et al., 2022), which considers genotypic uncertainties to directly output the most likely SFS. We compared three distinct demographic models, one with no gene flow between the three lineages, one allowing constant and asymmetric gene flow between populations, and a last model of secondary contact after the split between the eastern and north–western lineages. Population sizes could vary at each splitting time, and each lineage was assigned a specific effective population size. Parameters were estimated from the joint SFS using the likelihood approach implemented in fastsimcoal2.8 (Excoffier et al., 2013). We set inbreeding coefficients for all populations at 0.95. Based on interspecific whole genome alignments (see section above), we assumed a mutation rate of 3.3 x 10 ^– 9^ substitutions/site/generation (see Results) and a generation time of one year. Parameters with the highest likelihood were obtained after 40 cycles of the algorithm, starting with 100,000 coalescent simulations per cycle, and ending with 100,000 simulations. This procedure was replicated 50 times and the set of parameters with the highest likelihood was retained as the best point estimate. This estimate was then used as a set of starting parameters (--initValues option in fastsimcoal2) for a further round of 50 replicates. After confirming that likelihoods all converged to a narrow range, we used the estimate with the highest likelihood as the final set. We estimated 95% confidence intervals (CIs) using a nonparametric bootstrap procedure, using WINSFS to obtain block-bootstraps, sampling with replacement from 100 genome blocks. We repeated the parameter estimation procedure on these spectra, using the final inference obtained from the observed spectrum as starting parameters to reduce computation time.

*3.f. Demographic inference based on imputed genotypes (MSMC-IM)*

We used MSMC2 and MSMC-IM (Schiffels & Wang, 2020; Wang et al., 2020) on imputed genotypes to fit models of population size changes and migration rates through time. The method estimates rates of coalescence along genomes within and across populations, which can then be interpreted as changes in effective population sizes and migration rates through time. In addition to the genome masks already generated for heterozygosity and mappability, we masked regions for which the total depth of coverage was either lower than 100X or higher than 200X, retaining only sites with enough information for high-quality imputation. We ran MSMC2 using a time-patterning of 29 time segments with 27 free parameters (-p 1*2+25*1+1*2). We excluded from the analysis pairs of haplotypes belonging to the same individual, given their expected identity under selfing. For inference using haplotypes from the south-western and the north-western cluster, as well as for cross-coalescence analyses, we selected ten random pairs of haplotypes to reduce computational load. We used MSMC-IM with default parameters to estimate corrected rates of cross-coalescence between lineages. We converted coalescence rates to demographic parameters by assuming a mutation rate of 3.3 x 10 ^– 9^ substitutions/site/generation and a generation time of one year. To obtain an estimate of uncertainties on parameter estimates, we generated 20 bootstrapped datasets by randomizing both the ten pairs of haplotypes selected for inference and genome blocks of 250kb, sampled with replacement.

**S4. Environmental niche of *Linum bienne* from the Pliocene to present**

*Linum bienne* occurrences were retrieved with the R package rgbif (Chamberlain et al., 2025) from GBIF.org (<https://doi.org/10.15468/dl.nhbuhr> accessed on 03/07/2025) filtered in R using tidyverse (Wickham & Girlich, 2022) based on geographical coordinates approximating the species’ distribution area (longitude = [-25; 80], latitude = [21; 71]) and unlikely or impossible coordinates flagged by CoordinateCleaner (Zizka et al., 2019). Because occurrence density varied across the range, models were trained on France, Spain, and Portugal (high sampling intensity and representative climatic variation). WorldClim bioclimatic variables (Fick & Hijmans, 2017) show high correlation within the range of *L. bienne*, so we selected six related to temperature and precipitation seasonality (bio1, bio4, bio11, bio12, bio15, bio17), based on previous knowledge about the species (Landoni et al., 2024). Variables were extracted for nine periods: present; mid-Holocene (8 ka); LGM (22 ka); LIG–LGM transition (70 ka); Last Interglacial (130 ka); mid-Pleistocene (700 ka); early Pleistocene (1.3 Ma); late Pliocene (2.7 Ma); and early Pliocene (4.5 Ma). Present-day layers (10’ resolution) were obtained from WorldClim, and past climates (1 arc-degree resolution) via pastclim (Barreto et al., 2023; Holden et al., 2019). To further reduce multicollinearity the variance inflation factor (VIF) criterion was used to obtain a final predictor set of three variables including: bio4 (temperature seasonality), bio11 (coldest-quarter temperature), and bio12 (annual precipitation).

**REFERENCES**

Bolsheva, N. L., Melnikova, N. V., Kirov, I. V., Speranskaya, A. S., Krinitsina, A. A., Dmitriev, A. A., Belenikin, M. S., Krasnov, G. S., Lakunina, V. A., Snezhkina, A. V., Rozhmina, T. A., Samatadze, T. E., Yurkevich, O. Yu., Zoshchuk, S. A., Amosova, А. V., Kudryavtseva, A. V., & Muravenko, O. V. (2017). Evolution of blue-flowered species of genus Linum based on high-throughput sequencing of ribosomal RNA genes. BMC Evolutionary Biology, 17(S2), 253. https://doi.org/10.1186/s12862-017-1105-x

Cui, X., Li, E., He, J., Wang, Y., Shang, C., Zhong, B., Viruel, J., Dong, W. and Zhang, Z. (2025), Ancient hybridization drives arid adaptation and species diversification in Caragana (Fabaceae). New Phytol, 247: 2454-2472. https://doi.org/10.1111/nph.70360

Davies, R., Flint, J., Myers, S. et al. (2016). Rapid genotype imputation from sequence without reference panels. Nature Genetics 48, 965–969. https://doi.org/10.1038/ng.3594

Excoffier, L., Dupanloup, I., Huerta-Sánchez, E., Sousa, V. C., & Foll, M. (2013). Robust demographic inference from genomic and SNP data. PLoS genetics, 9(10), e1003905.

Fick, S. E., & Hijmans, R. J. (2017). WorldClim 2: New 1-km spatial resolution climate surfaces for global land areas. *International Journal of Climatology*, *37*(12), 4302–4315. https://doi.org/10.1002/JOC.5086

Galtier, N. (2024). An approximate likelihood method reveals ancient gene flow between human, chimpanzee and gorilla. Peer Community Journal, 4.

Garrison, E., & Marth, G. (2012). Haplotype-based variant detection from short-read sequencing. arXiv Preprint arXiv:1207.3907, 9. https://doi.org/arXiv:1207.3907 [q-bio.GN]

Goncalves, D. J., Simpson, B. B., Ortiz, E. M., Shimizu, G. H., & Jansen, R. K. (2019). Incongruence between gene trees and species trees and phylogenetic signal variation in plastid genes. Molecular phylogenetics and evolution, 138, 219-232.

Gutiérrez-Valencia J, Fracassetti M, Berdan EL, Bunikis I, Soler L, Dainat J, Kutschera VE, Losvik A, Désamoré A, Hughes PW, Foroozani A, Laenen B, Pesquet E, Abdelaziz M, Pettersson OV, Nystedt B, Brennan AC, Arroyo J, Slotte T. Genomic analyses of the Linum distyly supergene reveal convergent evolution at the molecular level. Curr Biol. 2022 Oct 24;32(20):4360-4371.e6. doi: 10.1016/j.cub.2022.08.042

Gutiérrez-Valencia, J., Fracassetti, M., Berdan, E. L., Bunikis, I., Soler, L., Dainat, J., Kutschera, V. E., Losvik, A., Désamoré, A., Hughes, P. W., Foroozani, A., Laenen, B., Pesquet, E., Abdelaziz, M., Pettersson, O. V., Nystedt, B., Brennan, A. C., Arroyo, J., & Slotte, T. (2022). Genomic analyses of the Linum distyly supergene reveal convergent evolution at the molecular level. Current Biology, 32(20), 4360-4371.e6. https://doi.org/10.1016/j.cub.2022.08.042

Gutiérrez-Valencia, J., Zervakis, P.-I., Postel, Z., Fracassetti, M., Losvik, A., Mehrabi, S., Bunikis, I., Soler, L., Hughes, P. W., Désamoré, A., Laenen, B., Abdelaziz, M., Pettersson, O. V., Arroyo, J., & Slotte, T. (2024). Genetic Causes and Genomic Consequences of Breakdown of Distyly in Linum trigynum. Molecular Biology and Evolution, 41(5), msae087. https://doi.org/10.1093/molbev/msae087

Hanghøj, K., Moltke, I., Andersen, P. A., Manica, A., Korneliussen, T. S. Fast and accurate relatedness estimation from high-throughput sequencing data in the presence of inbreeding. GigaScience, 2025, 8 (5), pp.giz034. 10.1093/gigascience/giz034

Hedrick, P. W., & Lacy, R. C. (2015). Measuring Relatedness between Inbred Individuals. Journal of Heredity, 106(1), 20–25. https://doi.org/10.1093/jhered/esu072

Hong-Yun Shang, Kai-Hua Jia, Nai-Wei Li, Min-Jie Zhou, Hao Yang, Xiao-Ling Tian, Yong-Peng Ma, Ren-Gang Zhang (2025). Phytop: a tool for visualizing and recognizing signals of incomplete lineage sorting and hybridization using species trees output from ASTRAL, Horticulture Research, Volume 12, Issue 3, uhae330, https://doi.org/10.1093/hr/uhae330

Incomplete lineage sorting and hybridization underlie of tree discordance in Petunia and related genera (Petunieae, Solanaceae). ecorRxiv (2026). Pedro H. Pezzi1,*, Lucas C. Wheeler2, Loreta B. Freitas1, Stacey D. Smith². https://doi.org/10.32942/X21W31

Innes, P.A., Smart, B. C., Barham, J. A. M., Hulke, B. S., Kane, N. C. (2023). Chromosome-scale Genome Assembly of Lewis Flax (Linum lewisii Pursh.). biorXiv. https://doi.org/10.1101/2023.10.10.561607

Jaegle, B., Pisupati, R., Soto-Jiménez, L. M., Burns, R., Rabanal, F. A., & Nordborg, M. (2023). Extensive sequence duplication in Arabidopsis revealed by pseudo-heterozygosity. Genome Biology, 24(1), 44. https://doi.org/10.1186/s13059-023-02875-3

Jeff J Doyle, Defining Coalescent Genes: Theory Meets Practice in Organelle Phylogenomics, Systematic Biology, Volume 71, Issue 2, March 2022, Pages 476–489, https://doi.org/10.1093/sysbio/syab053

Johnson, M.G., Gardner, E.M., Liu, Y., Medina, R., Goffinet, B., Shaw, A.J., Zerega, N.J.C. and Wickett, N.J. (2016), HybPiper: Extracting coding sequence and introns for phylogenetics from high-throughput sequencing reads using target enrichment. *Applications in Plant Sciences, 4*, 1600016. <https://doi.org/10.3732/apps.1600016>

Johnson, Pokorny, Dodsworth, Botigué, Cowan, Devault, Eiserhardt, Epitawalage, Forest, Kim, Leebens-Mack, Leitch, Maurin, D Soltis, P Soltis, Wong, Baker, Wickett, A Universal Probe Set for Targeted Sequencing of 353 Nuclear Genes from Any Flowering Plant Designed Using k-Medoids Clustering, Systematic Biology, Volume 68, Issue 4, July 2019, Pages 594–606,

Landoni, B., Suárez-Montes, P., Habeahan, R. H. F., Brennan, A. C., Pérez-Barrales, R. (2024). Local climate and vernalization sensitivity predict the latitudinal patterns of flowering onset in the crop wild relative Linum bienne Mill. *Annals of Botany, 134 (1)*, 117–130. https://doi.org/10.1093/aob/mcae040

Li, H., Handsaker, B., Wysoker, A., Fennell, T., Ruan, J., Homer, N., Marth, G., Abecasis, G., & Durbin, R. (2009). The Sequence Alignment/Map format and SAMtools. Bioinformatics, 25(16), 2078–2079. https://doi.org/10.1093/bioinformatics/btp352

McLay, T. G., Birch, J. L., Gunn, B. F., Ning, W., Tate, J. A., Nauheimer, L., ... & Jackson, C. J. (2021). New targets acquired: Improving locus recovery from the Angiosperms353 probe set. Applications in plant sciences, 9(7).

Meisner, J., & Albrechtsen, A. (2018). Inferring population structure and admixture proportions in low-depth NGS data. Genetics, 210(2), 719–731. https://doi.org/10.1534/genetics.118.301336

Morales-Briones, D. F., N.Lin, E. Y.Huang, D. L.Grossenbacher, J. M.Sobel, C. D.Gilmore, D. C.Tank, and Y.Yang. 2022. Phylogenomic analyses in Phrymaceae reveal extensive gene tree discordance in relationships among major clades. American Journal of Botany 109(6): 1035–1046. https://doi.org/10.1002/ajb2.1860

Morales-Briones, D.F., Liston, A. and Tank, D.C. (2018), Phylogenomic analyses reveal a deep history of hybridization and polyploidy in the Neotropical genus Lachemilla (Rosaceae). New Phytol, 218: 1668-1684. https://doi.org/10.1111/nph.15099

Nathaniel B. Edelman et al. (2019). Genomic architecture and introgression shape a butterfly radiation. Science 366,594-599. DOI:10.1126/science.aaw2090

Pockrandt, C., Alzamel, M., Iliopoulos, C. S., & Reinert, K. (2020). GenMap: Ultra-fast computation of genome mappability. Bioinformatics, 36(12), 3687–3692. https://doi.org/10.1093/bioinformatics/btaa222

Pokorny L, Pellicer J, Woudstra Y, Christenhusz MJM, Garnatje T, Palazzesi L, Johnson MG, Maurin O, Françoso E, Roy S, Leitch IJ, Forest F, Baker WJ and Hidalgo O. (2024). Genomic incongruence accompanies the evolution of flower symmetry in Eudicots: a case study in the poppy family (Papaveraceae, Ranunculales). Front. Plant Sci. 15:1340056. doi: 10.3389/fpls.2024.1340056

R Core Team. (2021). R: A Language and Environment for Statistical Computing. R Foundation for Statistical Computing. https://www.R-project.org/

Rasmussen, M. S., Garcia-Erill, G., Korneliussen, T. S., Wiuf, C., & Albrechtsen, A. (2022). Estimation of site frequency spectra from low-coverage sequencing data using stochastic EM reduces overfitting, runtime, and memory usage. Genetics, 222(4), iyac148.

S. Kong,C. Solís-Lemus, & G.P. Tiley, Phylogenetic networks empower biodiversity research, Proc. Natl. Acad. Sci. U.S.A. 122 (31) e2410934122, https://doi.org/10.1073/pnas.2410934122 (2025).

Skotte L, Korneliussen TS, Albrechtsen A. Estimating individual admixture proportions from next generation sequencing data. Genetics. 2013 Nov;195(3):693-702. doi: 10.1534/genetics.113.154138

Solís-Lemus C, Ané C (2016) Inferring Phylogenetic Networks with Maximum Pseudolikelihood under Incomplete Lineage Sorting. PLOS Genetics 12(3): e1005896. https://doi.org/10.1371/journal.pgen.1005896

Solís-Lemus, C., Bastide, P., & Ané, C. (2017). PhyloNetworks: a package for phylogenetic networks. Molecular biology and evolution, 34(12), 3292-3298.

Than, C., Ruths, D. & Nakhleh, L. (2008), PhyloNet: a software package for analyzing and reconstructing reticulate evolutionary relationships. BMC Bioinformatics 9, 322. https://doi.org/10.1186/1471-2105-9-322

Walker JF, Walker-Hale N, Vargas OM, Larson DA, Stull GW. 2019. Characterizing gene tree conflict in plastome-inferred phylogenies. PeerJ 7:e7747 https://doi.org/10.7717/peerj.7747

Wang, K., Mathieson, I., O’Connell, J., & Schiffels, S. (2020). Tracking human population structure through time from whole genome sequences. PLoS genetics, 16(3), e1008552.

Wickham, H., & Girlich, M. (2022). *tidyr: Tidy Messy Data* (1.3.1). https://tidyr.tidyverse.org, https://github.com/tidyverse/tidyr

Wu, T., Xu, A.N., Lei, Y. and Song, H. (2025), Ancient Hybridisation Fuelled Diversification in Acropora Corals. Mol Ecol, 34: e17615. https://doi.org/10.1111/mec.17615

Xu Y, Wei Y, Zhou Z et al. (2023). Widespread incomplete lineage sorting and introgression shaped adaptive radiation in the Gossypium genus. Plant Communications, 2023; 5. 10.1016/j.xplc.2023.100728

Yan, P., Guo, C., Cui, X., Li, E., Bai, Y., Roncal-Rabanal, M. R., ... & Dong, W. (2026). Phylogenomics unravels the early divergence and diversification in Bignoniaceae. Plant Diversity.

Yu, Y., Nakhleh, L. A maximum pseudo-likelihood approach for phylogenetic networks. BMC Genomics 16 (Suppl 10), S10 (2015). <https://doi.org/10.1186/1471-2164-16-S10-S10>

Zizka A, Silvestro D, Andermann T, Azevedo J, Duarte Ritter C, Edler D, Farooq H, Herdean A, Ariza M, Scharn R, Svanteson S, Wengstrom N, Zizka V, Antonelli A (2019). CoordinateCleaner: standardized cleaning of occurrence records from biological collection databases. *Methods in Ecology and Evolution*. [doi:10.1111/2041-210X.13152](https://doi.org/10.1111/2041-210X.13152)
